# Longitudinal structural variant phylogenies define tumor evolution under therapeutic selection pressure in metastatic prostate cancer

**DOI:** 10.1101/2025.02.01.636056

**Authors:** Yunzhou Liu, Jiaying Lai, Yi Yang, Mark C. Markowski, Emmanuel S. Antonarakis, Angelo M. De Marzo, Srinivasan Yegnasubramanian, Laura D. Wood, Laura A. Sena, Rachel Karchin

## Abstract

Therapeutic pressure shapes tumor evolution by selecting for subclones with distinct genomic architectures. In metastatic castration-resistant prostate cancer (mCRPC), structural variants (SVs) are central to this process but rarely incorporated into clonal reconstruction from bulk DNA sequencing. We describe SVCFit, a computational framework that estimates the cellular fraction of diverse SV classes from whole-genome sequencing and uses these estimates to reconstruct clonal evolutionary relationships. SVCFit explicitly models SV-type-specific breakpoint and breakend patterns and accounts for local copy-number context, enabling accurate SV cellular-fraction estimation across heterogeneous tumor genomes. On simulated datasets and in silico mixtures of metastatic prostate cancer samples, SVCFit achieves lower estimation error than SVclone, the current state-of-the-art for bulk-sequencing SV cellular-fraction estimation. Applied to longitudinal whole-genome sequencing from patients with mCRPC treated with bipolar androgen therapy, SVCFit reveals marked treatment-associated clonal reconfiguration, including contraction of highly rearranged subclones and expansion of resistant populations defined by distinct structural alterations. By enabling reconstruction of SV-defined clonal architecture from routine whole-genome sequencing, SVCFit expands the toolkit for studying tumor evolution under therapy in precision oncology. The COMBAT trial that provided these samples is registered on ClinicalTrials.gov (NCT03554317; first posted 13 June 2018).

## Introduction

Intratumor heterogeneity encompasses a broad spectrum of molecular alterations within a single tumor, including single nucleotide variants (SNVs), structural variants (SVs), copy number variants (CNVs), protein expression differences, and epigenetic modifications. Genomic instability and selective pressures can shape the emergence of diverse subpopulations, or subclones, that differ in their aggressiveness, metastatic potential, and resistance to therapy^1,2^. Reconstructing clonal architecture can reveal critical events that shape cellular phenotypes and underlying mechanisms^3^. Early research into cancer evolution focused primarily on aneuploidy^4^, and modern bulk-sequencing methods continue to leverage copy-number and allelic-imbalance signals to infer tumor purity, ploidy, and subclonal mixture proportions. The advent of Next Generation Sequencing (NGS) enabled scientists to investigate genetic alterations at the single-base level, using SNVs as markers for cancer progression^5^. More recently, paired-end sequencing has made the detection of structural variants more feasible and cost-effective, allowing their integration into studies of cancer evolution^6,7^. For both SNVs and SVs, estimating the cellular fraction, the proportion of cells that harbor a specific alteration is key to inferring their relative evolutionary ordering during tumor development.

Structural variants contribute significantly to genomic diversity and genome instability^8^. SVs come in various forms, such as insertions, deletions, tandem duplications, inversions, and translocations. Throughout, we use “duplication” to mean a tandem duplication unless otherwise specified. These large-scale changes can disrupt gene function by shifting or removing nucleotide sequences, which can lead to dysfunction in gene regulation or abnormalities in protein structure^9^. Because of their significance in shaping cancer genomes, SVs are critical to studying cancer genomics and metastasis, and they have direct relevance to therapeutic outcomes. Prostate cancer is increasingly recognized as a disease driven not only by small DNA sequence alterations but also by the presence and evolution of SVs. In primary prostate tumors, hallmark events such as the *TMPRSS2-ERG* fusion and losses of tumor suppressors like *PTEN* may arise through chromosomal rearrangements^10–13^. In metastatic castration-resistant prostate cancer (mCRPC), whole-genome sequencing has revealed that these structural alterations become even more pervasive, with complex events and noncoding rearrangements at loci such as *AR*, *MYC*, and *FOXA1* dominating the genomic landscape^14,15^. Under the selective pressure of androgen deprivation and modern *AR*-targeted therapies, clones carrying *AR* amplification, enhancer co-amplification, or other *AR* locus rearrangements are positively selected, while alternative trajectories involving SV-mediated loss of *TP53*/*RB1* and lineage plasticity give rise to *AR*-indifferent phenotypes. Understanding and monitoring this SV evolution is therefore important to predicting treatment response and prognosis, designing rational therapeutic combinations, and ultimately improving outcomes for men with metastatic prostate cancer.

Somatic alterations like SNVs and SVs in the human genome can be studied through bulk DNA sequencing, which is widely used due to its affordability. Incorporating somatic SVs detected from bulk sequencing into tumor evolution analysis can be done by first estimating variant allele frequency (VAF) at a specific locus. VAF is calculated as the ratio of altered reads to total reads at a genomic position, which works well for SNVs but does not transfer well to SVs: SNVs are detected at a single genomic position, while SVs are detected across multiple genomic positions and span multiple SV classes that each affect the genome differently, producing distinct patterns of altered and reference reads. While VAF-based approaches are widely used in clonal reconstruction, the process is noisy, due to confounding factors like aneuploidy and tumor impurity, both common in cancer samples^16,17^. A variant-centric strategy is to convert read support into cellular fraction, explicitly accounting for tumor purity and local copy-number state. While this method has been widely applied to SNVs^18–20^, to our knowledge, only one published method, SVclone^21^, estimates the cellular fraction of SVs from bulk sequencing data. Estimating the cellular fraction of a structural variant on a hemizygous chromosome calls for handling distinct from the diploid case, because a male sex chromosome carries no heterozygous germline SNPs to establish the local allele-specific copy number. This case matters in cancers of men, where the X chromosome carries recurrent prostate-cancer drivers, among them the androgen receptor, KDM6A and ZMYM3.

Despite their biological importance, SVs are typically excluded from tumor phylogenies inferred from bulk sequencing, largely due to challenges in estimating their cellular prevalence across diverse SV classes and copy-number contexts. Addressing this gap is critical for reconstructing tumor evolution under therapy. Here, we introduce SVCFit, a computational framework for estimating the cellular fraction of structural variants and use these estimates to reconstruct clonal evolutionary relationships from longitudinal whole-genome sequenced samples^22^.

## Results

### The SVCFit framework

The goal of this section is to estimate the SV cellular fraction (SVCF): the fraction of cells in a bulk tumor sample that carry a given structural variant. SVCF is defined as the number of carrier cells divided by the total number of cells sampled (Supplementary Note S1). Neither count is observable directly from bulk sequencing, but both are recovered from read counts through a single relationship,

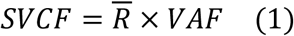

where VAF is the structural-variant allele frequency and R̅ is the average number of copies of the affected locus per cell, equal to 2 in a diploid region and to the mean number of copies of the locus per cell elsewhere; the hemizygous case is treated in “Estimation on hemizygous chromosomes” below. VAF measures how often the SV appears among sequenced alleles, and R̅ converts this allele-level frequency into a cell-level fraction. The conversion is needed because an SV resides on only a subset of a cell’s homologous copies: for a fully clonal heterozygous SV, every cell is a carrier (SVCF = 1) yet the variant is seen on one of two homologs (VAF ≈ 0.5), and the factor R̅ =2 recovers the true cell fraction. R̅ is fixed by the local copy number rather than estimated per cell; it equals 2 for diploid SV classes and 2(*r* − 1)/*r* for a tandem duplication of copy number *r* (Table 1; Supplementary Note S1).

**Table 1.** Closed-form SVCF for each SV class. *Breakpoints per cell* is the number of novel-junction breakpoints a single SV of the given class produces in a cell that carries it, a constant fixed by the SV class and, for tandem duplications, the local copy number *r*; it is the same in every carrier cell and is not estimated from data. R̅ is the average number of copies of the affected locus per cell across the sample, equal to the diploid value 2 for all classes except tandem duplications. On a hemizygous chromosome R̅ is the mean number of copies of the locus per cell, measured from read depth, and the breakpoint count for a tandem duplication becomes *r* − 1; see “Estimation on hemizygous chromosomes”. Every class reduces to the single observable form SVCF = R̅⋅VAF (Eq. 1); the per-class breakpoint counts are what make the derivations differ (full derivations, Supplementary Notes S1 and S7).

| SV class | Breakpoints per cell<br>(one SV of this class) | $\bar{R}$ (avg. copies of<br>affected locus per<br>cell) | Closed-form SVCF |
| --- | --- | --- | --- |
| Deletion | 1 | 2 | $\bar{R} \cdot VAF$ |
| Tandem duplication | $r - 2$ | $2(r - 1)/r$ | $\bar{R} \cdot VAF$ |
| Inversion | 2 | 2 | $\bar{R} \cdot VAF$ |
| Cut-paste<br>translocation | 3 | 2 | $\bar{R} \cdot VAF$ |
| Copy-paste<br>translocation | 2 | 2 | $\bar{R} \cdot VAF$ |
| Reciprocal<br>translocation | 4 | 2 | $\bar{R} \cdot VAF$ |

Applying the SNV-style cellular-fraction equation to SVs directly is insufficient, for two reasons that stem from how SVs appear in sequencing data. First, an SNV is detected at a single genomic position, where its variant and reference reads are counted together in one pileup. An SV is detected at several positions: SV-supporting reads arise at the breakpoint, where rearranged DNA is joined, while reference reads arise at the separate breakend location or locations, the original flanking sites (Fig. 1A-F). Computing VAF for an SV therefore means combining read counts gathered at different positions (defined in the next subsection), and the way they combine depends on the SV class. Second, the number of breakpoints per cell, and thus R̅, depends on the SV class as well (Fig. 1A-F). Estimating SVCF with the SNV formula, which assumes all supporting and reference reads come from a single position, systematically biases the result, particularly for tandem duplications and translocations. Throughout, breakpoint and breakend name the two read types an SV produces, SV-supporting reads at the novel junction and reference reads at the original flanking positions respectively, rather than the VCF/GATK convention in which a breakend denotes one end of a novel adjacency.

**Figure 1.**
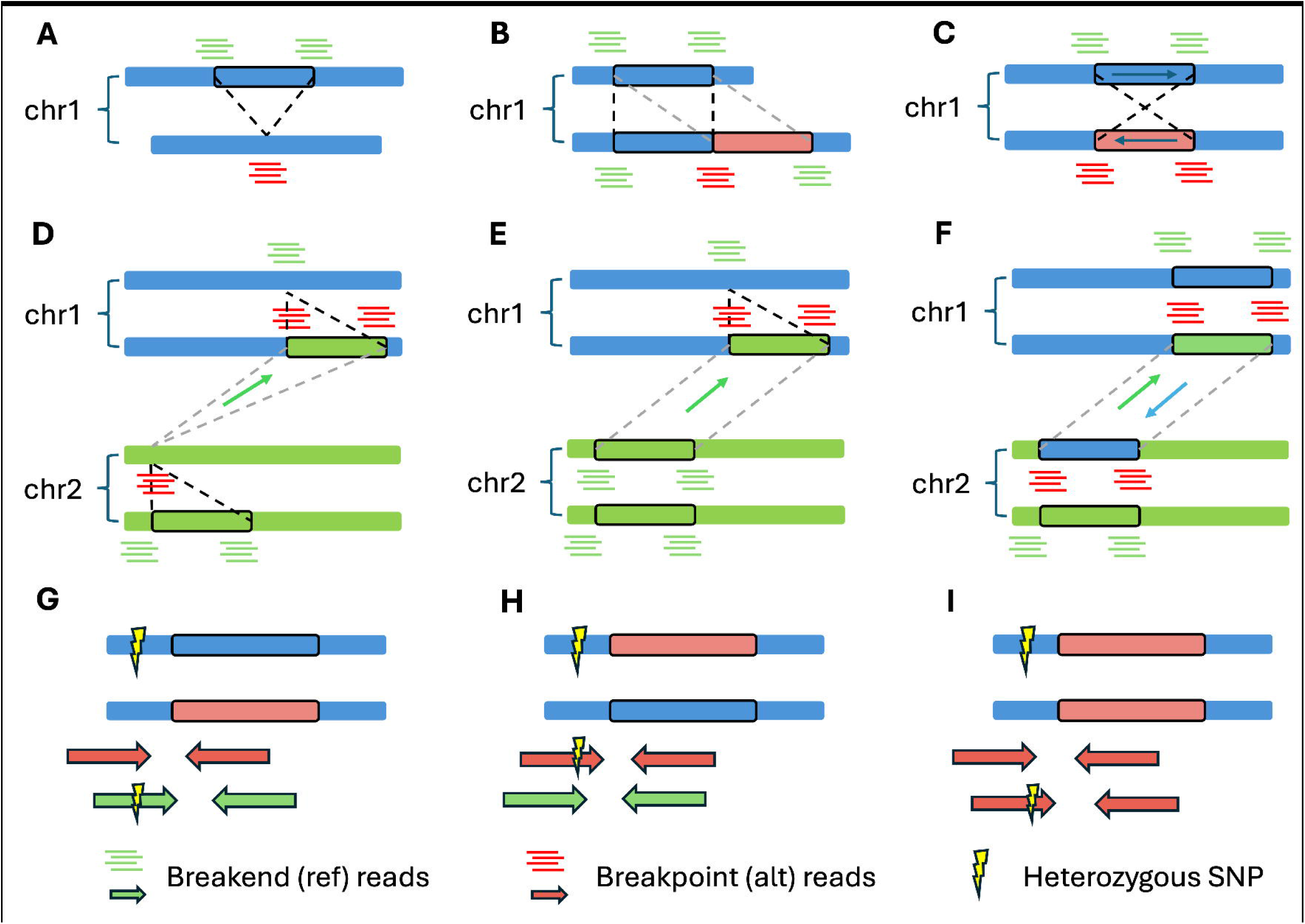
Read-evidence patterns of structural variant classes and haplotype-aware zygosity inference. (A-F) Schematics of the paired-end short-read evidence used by SVCFit for each supported structural variant (SV) class. Each chromosome is drawn as two bars for the two homologous haplotypes; black-outlined boxes mark the SV-affected segment and dashed lines link rearranged segments to their normal counterparts. Green pile-ups are breakend (reference) reads, red pile-ups are breakpoint (SV-supporting) reads. **(A)** Deletion. **(B)** Tandem duplication. **(C)** Inversion. **(D)** Cut-paste translocation: a chr1 segment is excised and inserted into chr2. **(E)** Copy-paste translocation: a chr1 segment is copied to chr2, leaving the source intact. **(F)** Reciprocal translocation: segments are exchanged between chr1 and chr2. **(G-I)** Haplotype-aware SV phasing and zygosity inference. Bars are the two haplotypes of one chromosome and the yellow lightning symbol marks a germline heterozygous SNP. Green arrows are breakend (reference) read pairs, red arrows breakpoint (SV-supporting) read pairs; a SNP symbol on an arrow means the captured allele is seen on that pair. **(G)** Heterozygous SV on the haplotype without the captured allele: the SNP appears only on breakend reads. **(H)** Heterozygous SV on the haplotype carrying it: the SNP appears on breakpoint but not breakend reads. **(I)** Homozygous SV: breakpoint reads arise from both haplotypes and both SNP alleles appear on them.

For each SV class SVCFit uses a closed-form estimator, an explicit formula that returns SVCF directly from the observed read counts without iterative fitting. Its two inputs are the counts of the SV-supporting and reference reads introduced above, denoted the breakpoint count (BPC) and breakend count (BEC), respectively. SVCFit takes the mean read depth at a breakpoint and at its breakends to be approximately equal, an assumption validated across simulated tumors (Supplementary Fig. S1), so these raw counts enter the estimate directly without a depth correction. Both are required: BPC measures how strongly the SV is supported, but a count on its own is not a fraction. BEC supplies the reference reads that form the denominator, so that the structural-variant allele frequency,

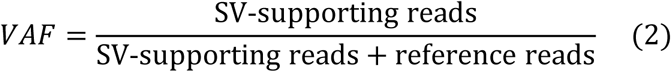

converts raw support into an allele frequency; R̅ then converts that frequency into the cell fraction through Eq. 1. In terms of these counts this is VAF = BPC/(BPC + BEC) for inversions and translocations and VAF = BPC/(BPC + ½·BEC) for deletions and tandem duplications, where the one-half corrects for reference reads counted on both flanks of the break interval (Table 1; Supplementary Note S1). Because the number of breakpoints per cell differs by SV class (Fig. 1A-F), the exact closed form differs accordingly; the per-class estimators are collected in Table 1.

SVs frequently arise within copy-number altered regions, where naive read-count ratios bias SVCF. SVCFit corrects for this by combining each SV’s zygosity and haplotype phase with the local allele copy ratio (Fig. 1G-I). Zygosity and phase are read from germline heterozygous SNPs near the breakpoint: when breakpoint-supporting reads carry only one heterozygous SNP allele, the SV is heterozygous and lies on the haplotype carrying that allele; presence of both alleles indicates a homozygous event.

The allele copy ratio (ACR), the copy number of the variant-bearing haplotype relative to the reference haplotype, is estimated from the allele frequencies of nearby heterozygous SNPs as *ACR* = *VAF*_ℎ*et*_/(1 − *VAF*_ℎ*et*_). ACR is continuous by construction: it equals 1 where the two haplotypes are balanced and moves away from 1 as one haplotype gains or loses copies. It is an intermediate quantity in the read-count adjustments below and is distinct from the integer copy number *r* that FACETS provides (Table 1). Because ACR is only a ratio, it cannot by itself say which haplotype changed: ACR > 1 can mean either the variant haplotype gained copies or the reference haplotype lost them. SVCFit resolves this directional ambiguity using the FACETS gain/loss classification.

Given the inferred zygosity, phase, and CNV class, SVCFit determines whether the overlapping CNV alters the breakpoint or breakend read count and adjusts the corresponding observable:

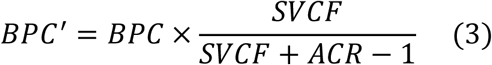

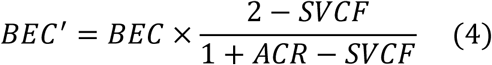

These adjustments are coupled through SVCF itself, so SVCFit applies an iterative procedure that we prove (Supplementary Note S2) converges to closed-form fixed points. When the breakpoint count is altered (a CNV in cis occurring after the SV), the fixed point is

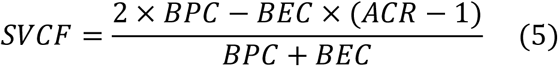

When the breakend count is altered (CNV preceding an in-cis SV, or CNV in trans), SVCF is the feasible root of the resulting quadratic,

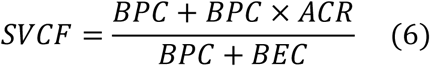

A property of Eq. 5 worth noting is that its sign is informative about the temporal order of an overlapping in-cis SV and CNV: under the algebraic constraint that an SV occurring after a CNV can affect at most the CNV-bearing alleles, the numerator is non-positive when the SV postdates the CNV and strictly positive when it predates (Supplementary Note S2.3; Supplementary Fig. S2). In practice, distinguishing these regimes from observed read counts requires sufficient breakpoint and breakend coverage to separate small numerator differences from sampling noise, so SVCFit uses the sign internally to choose between Eqs. 5 and 6 but reports it only as a directional indicator of relative SV-CNV ordering, not as a hard ordering call.

The allele copy ratio of Eq. 7 is estimated from the allele frequencies of nearby heterozygous germline SNPs, which do not exist on the X chromosome of a male subject. All COMBAT participants are male, so ACR is not estimable on chrX and the diploid conversion of Eq. 1 does not apply there. SVCFit therefore treats a declared hemizygous chromosome separately, replacing the constant 2 of Eq. 1 with the mean number of copies of the locus per cell, which we write cn and measure from read depth. Because the matched normal is also single-copy on chrX, the tumor-to-normal depth ratio at a segment gives cn directly once it is scaled by the sample’s autosomal ploidy. It is a continuous measurement and is not rounded to an integer copy number.

On a copy-neutral hemizygous locus cn = 1, a carrier cell has the structural variant on its only copy, and Eq. 1 reduces to SVCF = VAF. Where the locus is also copy-altered, counting alleles across the sample shows that the total number of alleles per cell equals cn whatever the order of the two events, so VAF is the number of variant alleles per cell divided by cn, and each ordering gives one closed form. When the copy-number change precedes the structural variant, each carrier cell holds the variant on one of its copies and SVCF = cn × VAF, which is Eq. 1 with R̅ cn. When the structural variant precedes the copy-number change, the amplified copies all carry it and SVCF = cn × VAF − (cn − 1). Both reduce to SVCF = VAF at cn = 1.

The two orderings are distinguished by the sign of the second expression, exactly as the sign of Eq. 5 distinguishes them on a diploid locus (Supplementary Note S2.3). If the copy-number change precedes the structural variant, the cells carrying the variant are a subset of those carrying the copy-number change, and that expression is then necessarily non-positive; if the structural variant arose first, it is the cellular fraction itself and is strictly positive. The ordering is therefore determined by the data, and no estimate of the copy-number change’s own cellular fraction is required.

A hemizygous tandem duplication is treated separately, because the duplication is itself the copy-number change and its novel junctions lie inside the region whose copy number it alters. A carrier chromosome bearing r tandem copies contributes r − 1 novel junctions, so cn = 1 + SVCF(r − 1) and SVCF = (cn − 1)/(r − 1). For this class VAF is a function of cn alone, so r cannot be determined from the locus and the cellular fraction cannot be separated from it. Since r ≥ 2, the value obtained at r = 2 is an upper bound, and SVCFit reports hemizygous tandem duplications as bounds rather than point estimates. A hemizygous deletion instead admits an internal check: it removes the only copy, so depth gives cn = 1 − SVCF while the junction reads give SVCF directly, and the two estimates are compared rather than combined.

Of the 132 structural variants called on chrX across the 22 subjects included in the phylogenetic analysis, 34 were resolved as preceding an overlapping copy-number change and 56 as following one, 15 lay on copy-neutral segments, 14 were deletions constituting their own copy-number change, and 13 hemizygous tandem duplications are reported as upper bounds. A cellular fraction was obtained for 123; the remaining 9 were excluded as infeasible.

The full procedure combines these components in a fixed order (Fig. 2). Each SV is first assigned to a class, which fixes its breakpoints per cell and R̅ (Table 1). SVCFit then counts the SV-supporting and reference reads and forms the variant allele frequency VAF from their ratio. In a diploid region the estimate is then complete: SVCF = R̅ × VAF (Eq. 1). When the SV overlaps a copy-number alteration, SVCFit adds copy-number context before applying Eq. 1. It phases the SV and infers its zygosity from nearby heterozygous SNPs, estimates the allele copy ratio ACR from those same SNPs, and takes the gain/loss direction and CNV class from FACETS. Together these determine whether the overlapping CNV has altered the count of SV-supporting or reference reads, which SVCFit corrects (Eqs. 3, 4); because each correction depends on SVCF itself, the estimate is refined by iteration to the closed-form fixed point (Eqs. 5, 6). The result is a cellular fraction for every SV; downstream clustering and phylogeny reconstruction (Fig. 2, Methods) assemble these into subclones and tumor-evolution trees.

**Figure 2.**
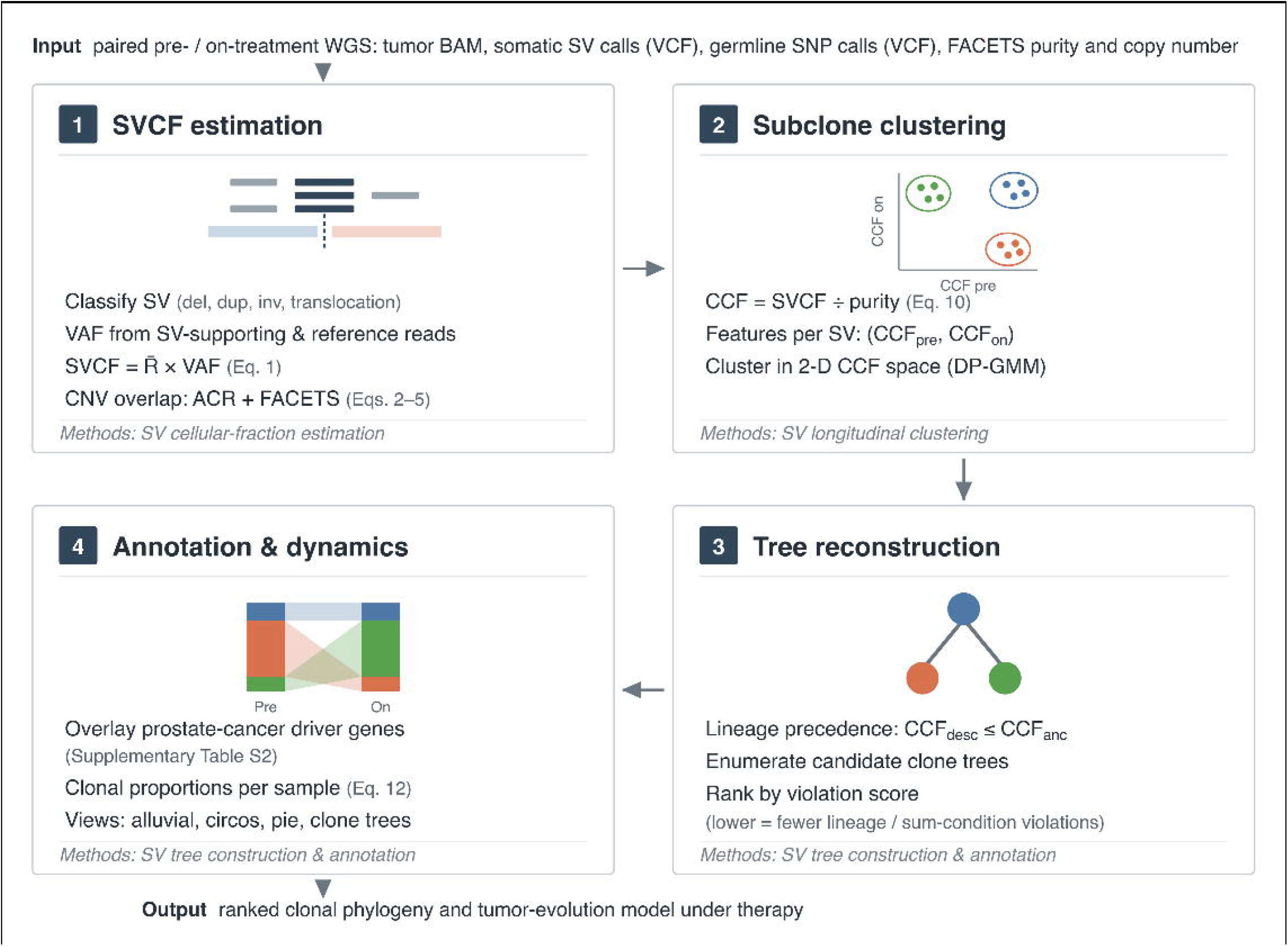
Overview of the SVCFit workflow. SVCFit reconstructs SV-based clonal phylogenies from paired pre-and on-treatment whole-genome sequencing. Inputs are the tumor BAM, somatic SV calls, germline heterozygous SNP calls for phasing, and allele-specific copy number and purity from FACETS^23^. The italic label in each panel names the Methods subsection specifying that step. **(1)** SVCF estimation. Each SV is typed, a variant allele frequency is formed from junction reads (dark) and flanking reference reads (gray), and converted to a per-SV cell fraction (Eq. 1), with a copy-number correction where the SV overlaps a CNV (Eqs. 3-6). **(2)** Subclone clustering. SVCF is converted to cancer cell fraction (Eq. 11) and SVs are clustered on their pre/on-treatment CCF pair with a Dirichlet-process Gaussian mixture model; each cluster is a candidate subclone. **(3)** Tree reconstruction. Candidate trees are enumerated under lineage precedence and ranked by a violation score; the top-ranked tree is retained. **(4)** Annotation and dynamics. Edges are annotated against prostate-cancer driver genes (Supplementary Table S2) and clonal proportions (Eq. 13) are displayed as alluvial, circos, pie and tree plots. Across panels 2-4, color denotes clone role: truncal (blue), declining (orange), emerging (green). BAT, bipolar androgen therapy; pre/on, pre-and on-treatment biopsies.

### SVCFit improves SV cellular fraction estimation compared to state-of-the-art methods

We benchmarked SVCFit against SVclone^21^, the current state-of-the-art for bulk-sequencing SV cellular fraction estimation, using synthetic tumor genomes generated by the haplotype-aware simulator VISOR^24^. We tested 75 scenarios spanning five SV-CNV overlap configurations, five SV tumor-purity levels (10%, 20%, 40%, 60%, and 80%), and three subclonal mixture settings, with replicate sampling and per-condition statistical comparisons described in Methods. In the four of the five configurations that include an overlapping copy-number event, the SVclone arm was supplied the ground-truth simulated cellular fraction to resolve per-SV multiplicity, following the SVclone authors’ evaluation convention, whereas SVCFit received no ground-truth information; the cellular-fraction comparison is therefore conservative for SVCFit wherever copy number is altered.

Accuracy was quantified as the signed error between true and estimated SVCFs (Eq. 12; positive values indicate underestimation, negative values indicate overestimation). Because each replicate averages thousands of simulated SVs, a replicate-level significance test is saturated, its p-values sitting at the resolution floor for nearly every condition and unable to distinguish a negligible difference from a large one; we therefore compare methods by mean signed error with a bootstrap 95% confidence interval and the paired effect size, and report exact and adjusted p-values in companion statistics tables rather than marking significance on the figures. Translocations refer to inter-chromosomal rearrangements unless otherwise specified. We report the comparison in stages: first an aggregate across SV classes and purities; then clonal deletions and tandem duplications in the baseline configuration and in the two configurations in which the SV arises before an overlapping copy-number change; next the subclonal conditions; and finally deletions in the two configurations in which the SV arises after a copy-number change.

Across the full five-purity range, SVCFit outperformed SVclone for inversions in 19 of 25 conditions and for translocations in 22 of 25 conditions (Supplementary Fig. S3). These two SV classes showed the largest number of conditions favoring SVCFit, although clonal translocations at 40% to 80% purity remained notable failure cases. For clonal deletions and tandem duplications, SVCFit had the lower absolute mean signed error throughout the baseline configuration; in the SV-before-CNV configurations the advantage was purity-dependent, holding at higher purity while SVclone was more accurate at low-to-mid purity, and the tandem-duplication advantage was restricted to ≥ 40% purity. Performance was comparable between methods for most subclonal conditions.

In clonal settings SVclone had the lower absolute mean signed error mainly for translocations, most consistently in the copy-number-altered configurations and at higher purity. It also held a smaller, low-to-mid-purity edge in the configuration where the SV precedes a cis duplication, for both clonal deletions and clonal inversions, an advantage that reversed by 80% purity. The translocation gap and these low-purity edges have different origins.

Clonal-translocation underestimation appears even in the baseline configuration, which has no overlapping copy-number event. SVCFit’s translocation cellular-fraction estimate increasingly underestimates the true value as purity rises, from near-zero mean signed error at 10% purity to recovering about two-thirds of the true value at 80%, a pattern not seen for deletions or duplications. We attribute this to translocations being the only SV class whose two breakends flank physically independent genomic loci, so, because SVCFit takes each breakend’s copy-number context from its maximum-overlap segment rather than averaging across the two, a translocation’s estimate can rest on the less reliable of two unrelated local contexts.

The low-purity edges, by contrast, are confined to the copy-number-altered configuration in which the SV precedes a cis duplication and to low-to-mid purity. That is one of the configurations in which the SVclone arm benefits from the ground-truth-assisted multiplicity step (Methods), so part of the apparent SVclone advantage there, for both the deletions and the inversions, may reflect that benchmark asymmetry rather than a genuine difference in accuracy.

For deletions in scenarios where a copy-number event on the same locus preceded the SV, the two methods were comparable overall, and for clonal deletions SVCFit had the lower absolute mean signed error in most conditions. This comparison is conservative for SVCFit: in these copy-number-altered configurations the SVclone comparison arm was provided the ground-truth simulated cellular fraction to disambiguate SV multiplicity (the SVclone authors’ evaluation convention; Methods), an assist SVCFit did not receive, so reaching parity, let alone an advantage, is achieved against a method handed the true cellular fraction. At 10% purity, performance differences narrowed for most SV types, consistent with the expected degradation of signal at extreme low tumor content. Fig. 3A shows three representative purities (10%, 40%, and 80%); the complete five-purity sweep is in Supplementary Fig. S3.

**Figure 3.**
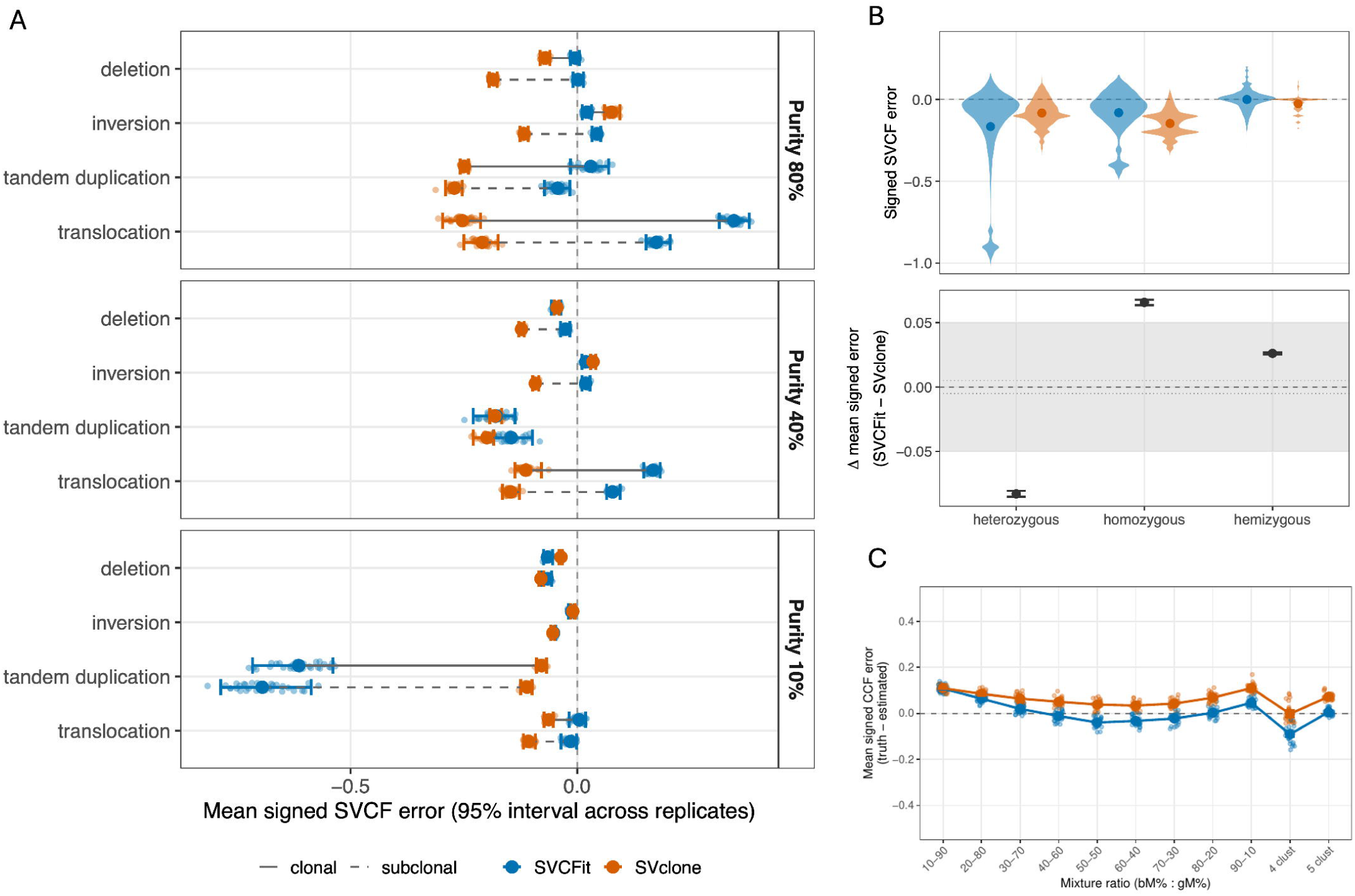
Benchmarking of SVCFit versus SVclone on simulated and prostate cancer mixture datasets. Throughout, SVCFit is blue and SVclone orange; clonal variants are solid lines and subclonal variants dashed. Error is truth minus estimate in every panel, so positive values indicate underestimation. Panels A and B report error in SVCF and panel C in CCF, the two being related by Eq. 11 (Methods); panel C uses CCF for comparability with the SVclone mixing benchmark, which reports CCF. **(A)** Mean signed SVCF error from VISOR^24^ simulations, pooling autosomal and hemizygous (chrX) structural variants, stratified by SV type (columns) and shown at three representative SV tumor purities (10%, 40%, 80%; rows). The complete five-purity sweep and per-condition breakdown are in Supplementary Fig. S3; per-cell effect sizes for the pooled comparison are in Supplementary Table S13. **(B)** Signed SVCF error at low tumor purity (10-20%) by zygosity, including hemizygous (chrX) loci. Top: per-SV signed-error violins with each method’s mean signed error overlaid; bottom: the paired between-method difference (SVCFit minus SVclone mean signed error) per zygosity with its bootstrap 95% confidence interval and a grey band marking the clone-assignment margin. Between-method effect sizes, confidence intervals, and adjusted p-values are in the companion statistics table (Supplementary Table S14). **(C)** Mean signed CCF error on the in-silico prostate cancer mixture benchmark of Cmero et al.^21^ Each x-axis position corresponds to one mixing condition: the nine three-cluster mixing ratios (bM:gM = 10:90 through 90:10) and the additional four-and five-cluster mixtures (“4 clust”, “5 clust”). Solid dots show per-replicate means; lines connect per-condition means; vertical bars indicate the 95% interval across replicates. Not all between-method differences are significant; per-condition effect sizes, confidence intervals, and adjusted p-values are in the companion statistics table (Supplementary Table S15).

We also analyzed the impact of zygosity on estimation precision at low tumor purity (10-20%; Fig. 3B). SVCFit achieved lower absolute mean signed error than SVclone for homozygous SVs, where both alleles carry the rearrangement and breakpoint read evidence is unambiguous. For heterozygous SVs, SVclone had the lower mean error, but this gap was driven almost entirely by low-purity heterozygous tandem duplications, which produced the long-error tail in Fig. 3B and the SV-type-stratified breakdown in Supplementary Fig. S3. Excluding these, SVCFit was more accurate for heterozygous inversions and translocations, while SVclone retained a modest advantage for heterozygous deletions.

This tail reflects SVCFit’s reliance on FACETS to genotype tandem duplications at low purity. When FACETS cannot confidently call the duplication’s own copy-number gain against the diluted low-purity background, SVCFit falls back to estimating that copy number from the same read-count-based quantity (R̅ × VAF) that forms the numerator of its cellular-fraction estimate, so the numerator and denominator cancel and the estimate is exactly 1 for every such event, regardless of the true value (affecting 99% of these events at 10% purity and 19% by 60%). Homozygous duplications, whose larger absolute copy-number change is more reliably detected, and deletions, which do not use this fallback, are unaffected.

Because these tandem duplications are copy-number-altered, the SVclone arm was supplied the ground-truth cellular fraction for them in the four configurations with an overlapping copy-number event, which fixes the very multiplicity that SVCFit must infer from FACETS. SVclone’s advantage on these events therefore reflects that assist rather than a genuine difference in low-purity tandem-duplication genotyping.

The comprehensive per-condition breakdown, stratified by SV-CNV overlap configuration, SV-CNV temporal ordering, zygosity, SV type, clonality, and the full five-purity range, is provided in Supplementary Fig. S3. Separate full-grid breakdowns by SV zygosity (Supplementary Fig. S4), by clonal versus subclonal SVs (Supplementary Figs. S5 and S6), and across the three subclonal-mixture settings (Supplementary Fig. S7) are also provided.

The benchmark above covers the autosomes. To evaluate the hemizygous estimators we generated a separate VISOR simulation carrying a hemizygous X chromosome, spanning 45 conditions with 30 replicates each, and compared SVCFit and SVclone on the same 4,935 ground-truth SVs. Uncertainty was quantified by a block bootstrap over replicates (B = 10,000), paired within each draw so that both methods are evaluated on the same replicate and the replicate-to-replicate variation common to both cancels. SVCFit detected 77.0% of simulated variants against 72.4% for SVclone (paired difference +4.6 percentage points, 95% CI 4.5 to 4.7), placed 66.6% of estimates within 0.05 of the true cellular fraction against 53.3% (+13.3, 12.8 to 13.7), and gave a mean absolute error of 0.0561 against 0.0888 (−0.033, −0.033 to −0.032). The margin is concentrated in deletions, where 82.6% of SVCFit estimates fell within 0.05 of truth against 53.3% for SVclone (+29.3, 28.8 to 29.9), the class the hemizygous forms are designed for. For inversions SVclone was the more accurate method, by 0.8 percentage points (53.4% against 52.7%; −0.8, −1.2 to −0.3), a difference whose paired interval excludes zero. As in the autosomal benchmark, SVclone was supplied the true tumor purity and SVCFit was not; both methods received the same variant calls and the same copy-number segmentation. Hemizygous tandem duplications are excluded from this comparison, because SVCFit reports them as upper bounds rather than as point estimates.

Next, we applied an in silico mixing benchmark developed by the SVclone authors, using two previously sequenced metastatic prostate tumor samples with known purities (49% and 46%) and shared ancestry^22^. Mixing the two source samples at nine pairwise ratios produced nine three-cluster mixtures (one source-A-only, one source-B-only, and one shared truncal cluster, with cellular fractions set by the mixing proportions); differential subsampling of odd and even chromosomes from each source generated additional four-cluster and five-cluster mixtures (Methods).

Across the nine three-cluster mixtures, SVCFit signed errors fluctuated near zero across mixing ratios (mean signed error approximately −0.04 to +0.11), with the largest magnitudes at the extreme 10:90 and 90:10 dilutions where one clone constitutes only 10% of the mixture. SVclone’s CCF values, computed with the analytical MapVaf2CcfPyClone convention used in its publication (Methods), showed a consistent positive offset across all nine three-cluster conditions (Fig. 3C). That convention resolves each SV’s multiplicity using the ground-truth cellular fraction, an assist SVCFit did not receive, so SVclone’s estimate is anchored to the true value. As in the VISOR benchmark, part of the offset therefore reflects that anchoring rather than a difference in subclonal-reconstruction accuracy.

For the four-and five-cluster mixtures, SVCFit retained low error in the five-cluster setting; the four-cluster mixture exhibited greater variance and was the only condition at which SVCFit showed worse performance than SVclone. This four-cluster exception reflects an upstream copy-number/purity-estimation instability specific to that simulated composition rather than a subclonal-reconstruction weakness: the FACETS-estimated tumor purity for this condition was both lower on average and about five-fold more variable across replicates than any other mixture, and per-replicate purity estimates correlated with SVCFit’s cellular-fraction error specifically in this condition (r = 0.65, p < 0.001) but not in the five-cluster condition (r = 0.13, p = 0.51).

SVCFit had the smaller absolute mean signed error in 7 of the 11 conditions (Benjamini-Hochberg-adjusted paired Wilcoxon, p < 0.05); the two methods did not differ significantly in three conditions (the 10:90, 50:50, and 60:40 mixtures, where SVCFit’s error was already near zero), and the four-cluster mixture favored SVclone, consistent with the copy-number and purity-estimation instability specific to that composition described above. Per-condition effect sizes, confidence intervals, and adjusted p-values are in the companion statistics table (Supplementary Table S15).

The mixture benchmark was also run with the hemizygous treatment, over 270 samples: nine mixture conditions at 30 replicates each. The four-and five-cluster mixtures are built from chromosome-split sources that cover the autosomes only and contain no X, so they are excluded by construction. Across 1,919 chrX variant rows, the ordering of an SV and an overlapping copy-number change was resolved for 753, with the copy-number change first in 456 and the structural variant first in 297, so both hemizygous forms carry substantial weight on these data. The principal limitation is one of resolution rather than of the estimator: 830 of the 1,165 chrX deletions could not be corroborated by depth, because 71.2% of them span less than a single 100 kb segmentation bin (median 22,141 bp), across which the depth difference is zero by construction. Their cellular fractions rest on junction reads alone, and in the simulation, where truth is known, uncorroborated deletions were the least accurate class (43.8% within 0.05 of truth, against 92.3% for those a depth estimate corroborates).

To quantify the practical consequences of these cellular-fraction errors, we scored SVCFit and SVclone head-to-head against the simulation truth on two discrete downstream decisions, pooling autosomal and chromosome X variants (Supplementary Fig. S12, Supplementary Table S12). Assigning each variant to the clone whose true cellular fraction is nearest, SVCFit misassigned 27.5% of variants against 45.4% for SVclone (paired difference −17.9 percentage points, 95% CI −18.1 to −17.7), a margin larger on chromosome X (−25.8 points) than on the autosomes (−8.2 points) and concentrated in the fragile subclonal decisions. On the ordering of subclones relative to the truncal clone, where an over-estimated subclonal fraction would imply an impossible ancestral placement, SVCFit produced 5.8% ordering violations against 18.3% for SVclone (−12.6 points, −13.9 to −11.2), worst at the tightest mixture. This complements the SVCFit-only phylogeny benchmark (Supplementary Figs. S8-S11).

### Longitudinal self-evaluation on paired pre/post-treatment simulations

To evaluate SVCFit’s downstream phylogenetic inferences against known ground truth, we generated paired pre-and post-treatment whole-genome simulations using VISOR^24^. Each simulated tumor carried a truncal clone (CCF = 1.0 at both timepoints) and two subclonal children whose CCFs cross under simulated therapeutic selection (one declines from CCF 0.83 to 0.17; one expands from 0.17 to 0.83). Sweeping SV tumor purity over {20%, 40%, 60%, 80%} (Methods), we evaluated three downstream capabilities per replicate: per-SV truncal/subclonal classification, per-clone CCF estimation, and rooted tree-topology recovery (Supplementary Note S3; Supplementary Fig. S8).

Truncal/subclonal classification and topology recovery were robust across the 20-80% purity range (Supplementary Fig. S8A,C). Per-SV balanced accuracy, the average of how often truly truncal SVs were correctly called truncal and how often truly subclonal SVs were correctly called subclonal, increased modestly from 0.79 at 20% purity to 0.89 at 80% purity, and was nearly identical at the pre-and post-treatment timepoints. This aggregate accuracy combines two components that trade off across the purity range rather than improving in parallel (Supplementary Fig. S8D): truncal recall (sensitivity) exceeds subclonal recall (specificity) at 20% purity (0.84-0.89 vs. 0.70-0.73), the two cross over by 60% purity (0.71-0.73 vs. 0.85-0.86), and both converge to high values by 80% purity, with subclonal recall modestly ahead (0.84-0.86 vs. 0.92-0.93). Rooted tree topology was correctly recovered in 83-88% of replicates (22/25 at 20%, 20/24 at 40%, 22/25 at 60%, 22/25 at 80%) with no monotonic dependence on purity. An illustrative recovered topology with paired pre/post centroid placement is shown in Supplementary Fig. S9.

Per-clone CCF point estimates show a systematic positive signed error (Supplementary Fig. S8B): the truncal cluster is underestimated by 15 to 26% across the purity range, and per-clone mean absolute error decreases modestly with purity (0.18 at 20% purity to 0.12 at 80%). This residual arises in the clustering step rather than in per-SV genotyping, and it does not affect the classification or topology results.

Two effects combine. The truncal CCF is bounded at 1.0, so any inferential noise can only push the inferred centroid below truth, producing a one-sided positive signed bias even under unbiased per-SV estimation. Larger in magnitude, the clustering step occasionally assigns individual truncal SVs to a subclonal cluster, and each such misassignment lowers the truncal centroid without changing the underlying per-SV evidence. This effect is strongest at intermediate purity (40 to 60%), where per-SV signal is adequate but the truncal and subclonal centroids are least separable, which is why the truncal bias is not monotonic in purity.

Per-SV genotyping by SVtyper^25^ shows no systematic undercount: for truncal SVs the ratio of raw SV cellular fraction to purity has a median near 1.0 at every purity, and any residual shortfall at high purity is small (at most 5.1%), consistent with the modest sensitivity limits of breakpoint-resolved SV genotyping reported in independent benchmarks^26^ but far too small to account for the cluster-level underestimate.

Per-SV read support and SV-type composition were tested directly as covariates of the residual error (Supplementary Figs. S10, S11); read support explained little of the variation, while subclonal CCF error grew with the number of breakend-defined translocations assigned to a cluster (Spearman rho of about 0.5), consistent with the harder placement of BND-defined events rather than a read-depth limitation. Topology recovery and truncal/subclonal classification, the inferences on which the longitudinal phylogenetic analysis depends, are unaffected by this magnitude bias.

### Longitudinal analysis of SV-defined clones in mCRPC subjects receiving bipolar androgen therapy

To demonstrate the clinical utility of SVCFit in resolving tumor evolution, we analyzed longitudinal whole-genome sequencing (WGS) data from 11 evaluable subjects with metastatic castration-resistant prostate cancer (mCRPC) treated with bipolar androgen therapy (BAT)^28,29^, a form of supraphysiologic testosterone treatment, on the COMBAT trial^30^. BAT is the cyclical use of high dose testosterone as treatment for CRPC, which is associated with a 30% response rate, but fairly rapid acquired resistance with a median progression-free survival of 6 months^31^.

Subjects on the COMBAT study were required to undergo paired sequential metastatic tumor biopsies at baseline (pre-treatment) and after 3 months of BAT treatment. SV-based phylogenies were reconstructed using SVCFit for each subject; for direct comparison, SNV-based phylogenies, with prostate-cancer driver annotations applied to clustered SNVs, were reconstructed from the same WGS data in parallel using PICTograph^19^. Per-subject phylogenies and a side-by-side SV-versus-SNV comparison across the cohort are presented in Supplementary Note S4, with the paired SV and SNV trees for all 11 subjects shown in Supplementary Figs. S14-1 to S14-11.

Two subjects, 10 and 31, are presented in detail below because they initially demonstrated a biochemical response (assessed by serum prostate-specific antigen (PSA) levels) to BAT, followed by a subsequent PSA rise at the time of the second biopsy, suggesting early development of acquired therapy resistance under selection pressures (Fig. 4). SVCFit-inferred SV cellular fractions, DP-GMM clustering, and modified Gabow-Myers tree reconstruction (Fig. 2; Methods, “SV Longitudinal Clustering and Dynamics” and “SV Tumor Evolution Tree Construction and Annotation”) were applied uniformly across all 11 evaluable subjects (Supplementary Note S4); we highlight Subjects 10 and 31 here to illustrate how this reconstruction informed our interpretation of acquired therapy resistance.

**Figure 4.**
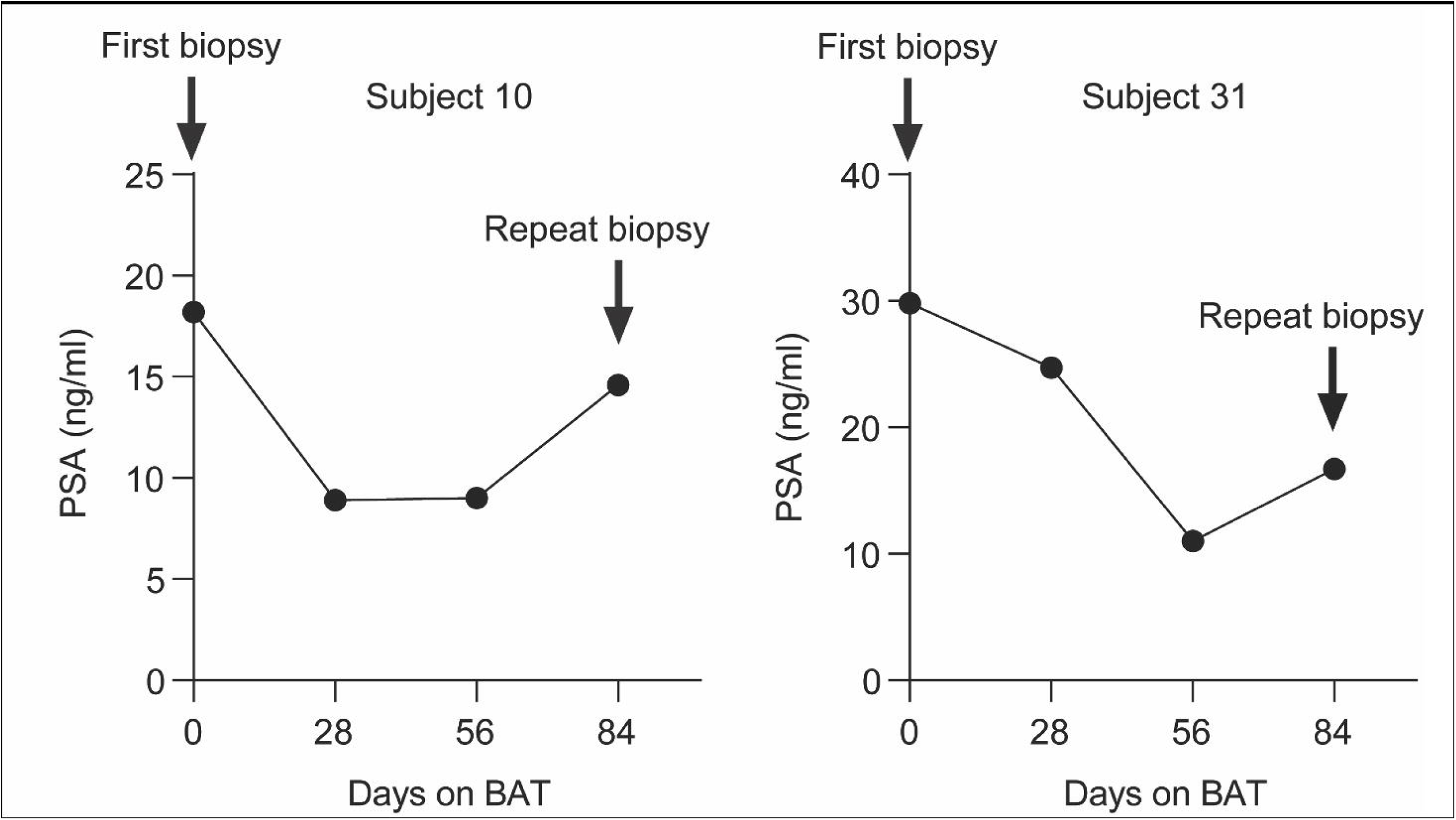
PSA dynamics during bipolar androgen therapy (BAT) and timing of biopsies. Two side-by-side panels, titled “Subject 10” and “Subject 31”, show serum prostate-specific antigen (PSA, ng/mL; linear scale) from the COMBAT trial clinical record plotted against days on BAT at days 0, 28, 56, and 84. Y-axis ranges differ between panels to accommodate per-subject PSA levels (Subject 10: 0-25 ng/mL; Subject 31: 0-40 ng/mL). Both subjects exhibit an initial decline in PSA followed by a rise by day 84 (week 12), consistent with acquired therapy resistance. Vertical arrows mark the time points of the pre-BAT first biopsy (day 0) and the on-BAT repeat biopsy (day 84).

In Subject 10, SVCFit resolved three distinct clonal populations, consistent with a branched evolutionary architecture (Fig. 5A; tree-display conventions and the breakpoint-versus-copy-number annotation rule are defined in the Fig. 5 legend and Methods). The truncal cluster of SVs, annotated to Clone 1 in Fig. 5A, was inherited by all tumor cells at both time points and contained canonical prostate cancer alterations, including *TP53* deletion^14,15^, *CHEK2* intragenic deletion, *MDM2* duplication^32,33^, and the *TMPRSS2-ERG* fusion^12,13^ (Although *TMPRSS2-ERG* was directly genotyped only in the pre-BAT WGS, the on-BAT BAM retained one supporting read and ERG-positive immunohistochemistry was confirmed in both biopsies. Of the SVs assigned to the truncal cluster, none were absent from the on-BAT sample after read-level rescue; mean breakpoint sequencing depth was 36× in the pre-BAT biopsy and 17× in the on-BAT biopsy. We therefore place the fusion on the trunk and attribute its absence from the on-BAT VCF to reduced tumor purity and lower effective coverage rather than subclonal loss; see Methods, “Rescue of key driver events”.) Following BAT, we observed a marked clonal reconfiguration, characterized by the disappearance of Clone 2 and the expansion of Clone 3 (Fig. 5B, 5C).

**Figure 5.**
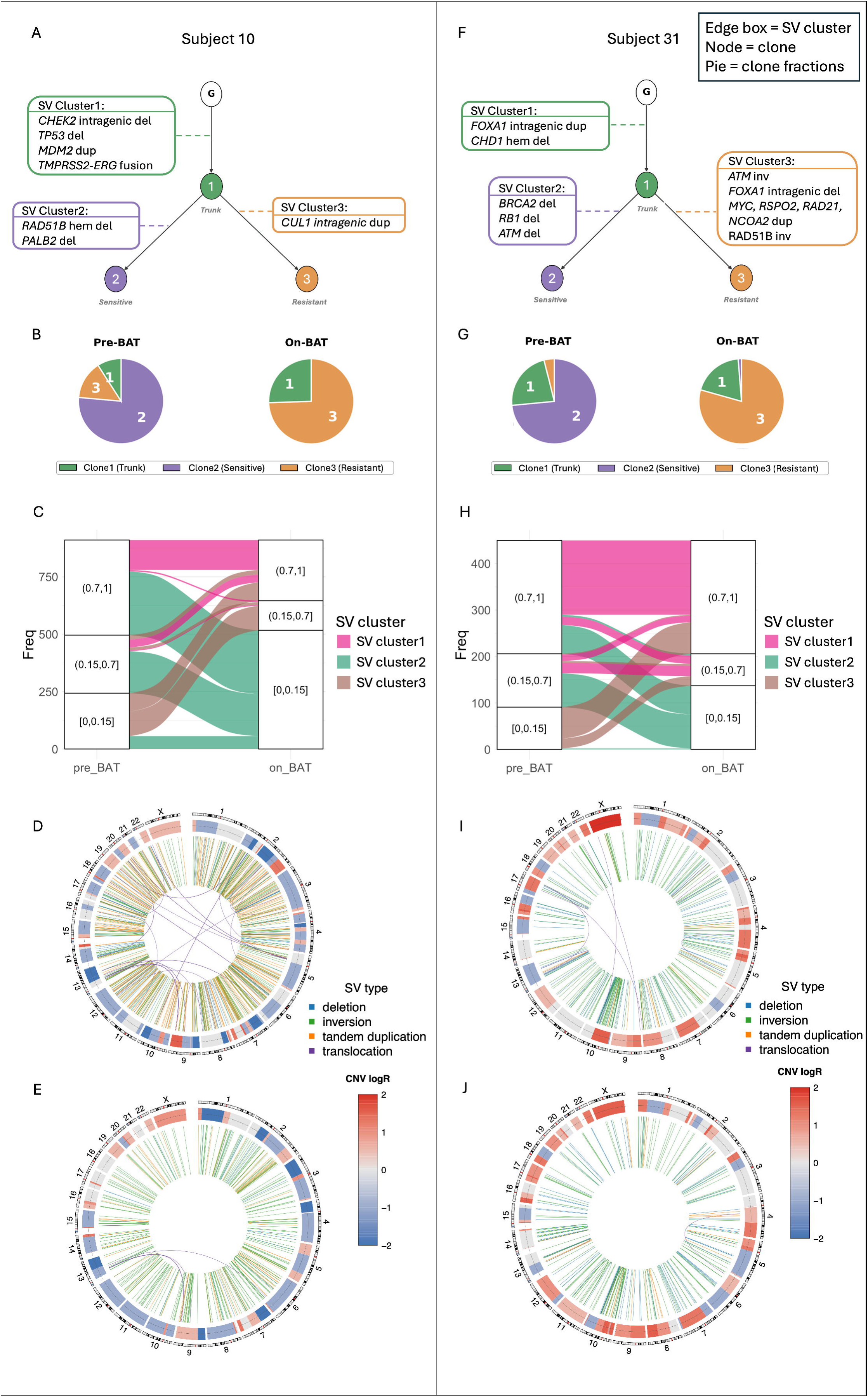
Longitudinal genomic landscape and tumor evolution in COMBAT mCRPC subjects treated with bipolar androgen therapy (BAT). Two subjects are shown side-by-side with the same five panel types in fixed order: phylogenetic tree, clone-proportion pie chart, alluvial plot, pre-BAT circos and on-BAT circos. Subject 10 occupies panels A-E and Subject 31 panels F-J. Reconstruction details, including SV inclusion criteria and clustering, are in Methods (“SV Longitudinal Clustering and Dynamics” and “SV Tumor Evolution Tree Construction and Annotation”). **(A, F)** SVCFit-inferred phylogenetic trees. Edge lengths indicate inferred ordering, not elapsed time. Only SVs overlapping the driver gene list of Hosseini et al.^37^ are annotated. **(B, G)** Pie charts of clone proportions (fraction of tumor cells per clone, not per-cluster prevalence) for the Pre-BAT and On-BAT biopsies. Slice colors match the tree nodes in panels A/F (Clone 1 = truncal; Clone 2 = sensitive; Clone 3 = resistant). **(C, H)** Alluvial plots tracing cluster-level SVCF between the pre-and on-BAT biopsies. SVCF is binned into (0, 0.15] (low-frequency or undetected), (0.15, 0.7] (subclonal) and (0.7, 1] (clonal or near-clonal). Stratum height is proportional to SV count and ribbons trace per-cluster shifts, colored as the clusters in panels A/F. SVs detected at only one timepoint are assigned SVCF = 0 at the other and fall in the lowest bin there. **(D, E, I, J)** Circos plots summarizing genome-wide structural variation per sample (D: Subject 10 pre-BAT; E: Subject 10 on-BAT; I: Subject 31 pre-BAT; J: Subject 31 on-BAT). The outer ring shows chromosomes 1-22 and X, and the inner ring somatic copy-number state as a blue-to-red log2 copy-ratio gradient (blue, loss; red, gain; scale beneath each circos as “CNV log2”). Curved interior links are individual SVs colored by type: blue (deletions), orange (tandem duplications), green (inversions), purple (translocations).

Clone 2 was notable for extensive genome-wide rearrangement: SV cluster 2 accounted for all 21 translocations detected in the pre-BAT biopsy and for 269 of its 297 duplications, and carried major structural alterations across multiple DNA-repair pathways, including hemizygous deletion of RAD51B and deletion of PALB2. In contrast, Clone 3 was defined by an 11 kb intragenic duplication within *CUL1* (chr7:148,755,369-148,766,429; hg38; Supplementary Fig. S16), the catalytic subunit of the SCF (SKP1-CUL1-F-box) E3 ubiquitin ligase complex. *CUL1* is a direct transcriptional target of c-MYC^34^. As the catalytic backbone of SCF^SKP2^, CUL1 mediates degradation of p27^Kip1^ and other cell-cycle regulators, with documented relevance to prostate cancer progression^35^. We treat this duplication as a candidate resistance-associated structural event, consistent with the broader observation that structural rearrangements in mCRPC can affect transcriptional programs of driver genes^14,15^; direct functional consequence of the *CUL1* intragenic duplication on *CUL1* expression will require clone-resolved transcriptional validation, which we discuss as a limitation. The expansion of Clone 3 under therapy illustrates how SVCFit can nominate candidate resistance-associated structural events that may be overlooked by SNV-centric analyses. The phylogeny reflects relative ordering of SV-defined clones constrained by pre-and on-treatment sampling, enabling direct comparison of clonal composition before and during therapy.

Longitudinal analysis on Subject 10 further revealed a remodeling of the genomic landscape following BAT. The pre-BAT (baseline) biopsy exhibited a hyper-rearranged phenotype indicative of pronounced genomic instability, with 21 translocations and 297 duplications (Fig. 5D). In contrast, the on-BAT sample showed a 10.5-fold reduction in translocations and a 5-fold reduction in duplications (2 translocations, 56 duplications; Fig. 5E; Supplementary Fig. S13A-C). To determine whether these differences could be explained by lower mean coverage in the on-BAT sample (36× pre-BAT, 17× on-BAT), we performed *in silico* downsampling of the pre-BAT DNA sequencing data to match the effective tumor depth of the on-BAT sample, repeated across 20 bootstrap replicates with distinct random seeds. Across replicates, the downsampled pre-BAT yielded a mean of 5.3 translocations (95% CI: 4.33-6.17) and 43 duplications (95% CI: 40.61-45.49). The translocation count remained higher in the downsampled pre-BAT than in the on-BAT sample (mean 5.3 vs. 2 observed), indicating a residual ∼2.6-fold difference not accounted for by coverage and consistent with a biological reduction in translocations driven by the elimination of the highly rearranged Clone 2. In contrast, the downsampled duplication count (mean 43; 95% CI: 40.61-45.49) fell below the on-BAT count (56), with the on-BAT value exceeding the bootstrap CI upper bound, indicating that the apparent reduction in duplications is fully attributable to lower sequencing depth in the on-BAT biopsy, with no evidence of net biological loss of duplication-bearing clones.

Remarkably, a similar pattern of branched evolution and clonal replacement was observed in Subject 31 (Fig. 5F). The truncal cluster of SVs in this tumor (Clone 1), inherited by all tumor cells at both time points, comprised CHD1 hemizygous deletion and a focal FOXA1 duplication (exon 2-txEnd). BAT treatment coincided with the divergence of two subclonal populations. Clone 2, which harbored *BRCA2*, *RB1*, and *ATM* deletions (a candidate compound HR-pathway defect^36^, with the caveats noted in Supplementary Table S2 *RB1* row), contracted following therapy (Fig. 5G, 5H). Conversely, Clone 3 expanded to become the dominant population and was defined by a characteristic set of structural alterations, including multiple ATM inversions predicted to be inactivating, FOXA1 intragenic deletion (exon 2-exon 2), duplications of NCOA2, MYC, RAD21 and RSPO2, and RAD51B inversion. In each *ATM* inversion, one breakpoint mapped within the *ATM* gene body (txStart-intron5), truncating the coding sequence.

Together, these observations demonstrate that mCRPC tumors can evolve through the selective expansion or contraction of SV-defined subclones during therapy. By estimating the cellular prevalence of these events, SVCFit enables inference of clonal evolutionary relationships, distinguishing early truncal SVs in Subject 31, including *CHD1* hemizygous deletion and focal *FOXA1* duplication, from later-arising subclonal alterations whose prevalence changed during BAT treatment. In Subject 31, Clone 2, which harbored *BRCA2*, *RB1*, and *ATM* deletions, contracted during treatment, while Clone 3, carrying predicted *ATM*-inactivating inversions and *NCOA2* duplication, expanded; the transcriptional implications of the Clone 3 alterations are considered in the Discussion.

Consistent with these findings, the on-BAT sample from Subject 31 exhibited a seven-fold reduction in translocations, all of which were confined to Clone 2, alongside a modest increase in duplications (pre-BAT: 7 translocations, 16 duplications; on-BAT: 1 translocation, 18 duplications; Fig. 5I, 5J; Supplementary Fig. S13D-F). As both samples had comparable tumor purity (∼0.6), no downsampling was required. Collectively, these cases suggest that BAT can selectively eliminate highly rearranged subclones, while leaving the dominant genomic architecture of resistant disease largely intact.

Across the broader COMBAT cohort, SV-based phylogenies resolved branched clonal architecture and assigned candidate sensitive-clone alterations in all 11 evaluable subjects and resistant-clone-specific alterations in 9 of 11, based on the prostate-cancer driver gene list of Hosseini et al.^37^ SNV-based phylogenies, even after annotating clustered SNVs against prostate-cancer driver genes, yielded actionable driver-gene information in only one subject (Subject 20, truncal BRCA2 frameshift together with the truncal AR ligand-binding-domain mutations p.Leu702His and p.Thr878Ala, and a SPOP Y83C subclone that fell from 97% of tumor cells pre-BAT to undetectable on-BAT), carried driver-gene SNVs that were truncal or otherwise non-distinguishing in three subjects (6, 7, 10), carried clone-distinguishing but non-actionable annotations in two subjects (12, 47), carried no annotated driver-gene SNVs in three subjects (1, 13, 49), and, in two subjects, implicated the same gene as a structural event: ZFHX3 in Subject 18, where both alterations fall on the BAT-sensitive clone, and NCOA2 in Subject 31, where the missense marks the declining clone and the duplication the expanding one.

Because SVCFit resolves structural variants on the X chromosome, the cohort analysis included 123 chrX events with estimated cellular fractions across the 22 subjects, among them variants spanning the AR, KDM6A and ZMYM3 loci; these were megabase-scale rearrangements containing the genes rather than focal alterations of them, and none introduced a subclone-defining event beyond those already resolved on the autosomes.

Candidate SV-driven resistance mechanisms identified across the cohort included an intragenic *CUL1* duplication (Subject 10), *FOXA1* structural remodeling (Subject 31), *MDM2 duplication*^32,33,38^ (Subject 20), AURKA duplication (Subject 6), *NCOR1* inversion (Subject 13), *BRAF duplication*^32,39^ with SETD2 deletion on the resistant clone (Subject 1), and ONECUT2 duplication with RB1 inversion (Subject 49), which together support an AR-independent survival program, while the BRCA2 deletion carried by the same clone does not itself confer resistance; RB1 loss is HR-supporting but not equivalent to BRCA1/2 loss^40^ (Supplementary Table S2 *RB1* row).

None of these candidate mechanisms was identified by the corresponding SNV trees. This does not by itself indicate that the loci are biologically SV-specific: an SNV-based reconstruction can place a gene on a tree only if that gene carries an informative, clustering-confident somatic point mutation, and most SNV-based phylogeny methods, including PICTograph, do not model copy-number loss, so a gene inactivated only by a structural deletion is intrinsically invisible to the SNV tree regardless of reconstruction quality. The SV-versus-SNV comparison therefore reflects, in part, what each data type and method can detect, not only differences in true clonal architecture. The reverse limitation also applies. In Subject 20 the AR ligand-binding-domain mutations p.Leu702His and p.Thr878Ala, both truncal, were missing from the SNV tree until they were recovered from the caller’s germline-risk-filtered output (Methods, “Rescue of key driver events”), so what an SNV tree can report is bounded by the upstream call set as well as by the reconstruction method.

Per-subject phylogenies, the side-by-side SV-versus-SNV comparison, and aggregate cohort summaries are presented in Supplementary Note S4, with the paired SV and SNV trees for all 11 subjects in Supplementary Figs. S14-1 to S14-11 and the pre-versus on-BAT SNV co-clustering for Subjects 13 and 31 in Supplementary Fig. S15. Literature support for the driver-status of each annotated gene, organized by pathway, with primary references, is compiled in Supplementary Table S2.

## Discussion

While clonal reconstruction in cancer genomics has historically relied on single nucleotide variants for clonal reconstruction^41^, this study demonstrates that genomic structural variants are not merely “co-passengers” but may shape clonal architecture and evolution^6,7,14,15^. By explicitly modeling the SV breakpoint patterns and the copy-number context, SVCFit provides a level of precision in cellular fraction estimation that has been difficult to achieve with bulk sequencing-based approaches. Across the simulated benchmark, SVCFit achieved lower cellular-fraction error than SVclone overall while operating without the ground-truth cellular fraction that SVclone’s evaluation convention supplies in copy-number-altered regions; the conditions where SVclone was more accurate coincide with that assist and so cannot be read as a genuine algorithmic advantage. Benchmarking against SVclone^21^ shows that failure to account for these structural features can bias clonality estimates, potentially misclassifying truncal events as subclonal or obscuring emerging resistant populations.

The application of SVCFit to the COMBAT trial patients offers a high-resolution view of how metastatic castration-resistant prostate cancer (mCRPC) navigates therapeutic bottlenecks. In Subject 31, *FOXA1* showed a truncal focal gene-body duplication spanning exon 2 through the transcript end (CN ∼3 pre-BAT; ∼5 on-BAT), while a dominant resistant subclone acquired a small intragenic deletion within exon 2. This suggests that therapeutic pressure during BAT may select for *FOXA1* locus remodeling with potential consequences for androgen receptor cistrome regulation.

Because this resistant subclone also carries an *NCOA2* duplication, and an NCOA2 duplication would conventionally be expected to enhance AR coactivation, the concurrent *FOXA1* alteration could redirect AR/coactivator activity toward an altered transcriptional program rather than simply amplifying the canonical AR-target repertoire associated with BAT sensitivity; these hypotheses require transcriptional and chromatin-state validation. Routine gene-level copy-number reporting would likely summarize *FOXA1* as a single dosage state, masking subclonal gene-body alterations that emerge in resistant lineages. Furthermore, the expansion of Clone 3 in Subject 10, characterized by an 11 kb intragenic duplication within *CUL1*, illustrates how SVCFit can pinpoint intragenic structural rearrangements at cell-cycle and ubiquitin-ligase loci, events that are difficult to detect with SNV-focused pipelines.

Interestingly, in both of these cases of early acquired resistance to BAT, we observed eradication of clones carrying multiple HR-pathway alterations: *BRCA2* deletion (Subject 31) or compound structural alterations spanning PALB2 and RAD51B (Subject 10). While homologous recombination repair deficiency (HRD) has been proposed to be a predictive biomarker of response to BAT^42,43^, we previously reported that it was not associated with durable response in the COMBAT study^44^. Application of SVCFit to these cases suggests that HRD may in fact be a valid predictive biomarker of response to BAT, but these responses can be limited by tumor HR status heterogeneity enabling outgrowth of resistant HR-proficient clones. Notably, subjects enrolled on the COMBAT trial had progressed on more prior therapies than those on previous trials of BAT^28,30,31,45, 46^and therefore may have had a higher degree of intra-and intertumoral heterogeneity and plasticity that enables acquired therapy resistance by clonal selection^3,47^.

We acknowledge the rising utility of single-cell sequencing in resolving tumor heterogeneity; however, technical limitations such as allelic dropout and high costs currently restrict its use in routine clinical practice^48,49^. SVCFit complements these approaches by recovering subclone-level resolution of SV-defined clonal architecture from standard bulk WGS, distinguishing truncal from subclonal events and reconstructing branched phylogenies in the cases shown here. This scalability makes SVCFit a viable candidate for integration into molecular tumor boards, where it might help clinicians distinguish stable truncal alterations from emerging resistant subclones.

SVCFit depends on the quality and characteristics of upstream SV calling. While the framework performs robustly for simple and moderately complex rearrangements when applied to consensus calls from established short-read SV callers, differences across callers in breakpoint-detection sensitivity, definitions of split-read and discordant-pair support, and internal quality filtering produce different per-SV reference-and alternate-supporting read counts and may therefore yield different downstream SVCFit estimates from the same underlying data; we mitigated this in the COMBAT analysis by using a Manta^50^ + Delly^51^ + GRIDSS^52^ consensus merged with SURVIVOR^27^ before genotyping, but a residual sensitivity to caller choice remains. Highly complex processes such as chromoplexy, chromothripsis, and breakage-fusion-bridge cycles generate multi-breakpoint architectures that are not yet explicitly represented in the current model, even when the breakpoints that make up these events are detectable with short-read sequencing. Future extensions will address this by incorporating structure-aware representations of complex SVs and by integrating SNVs, CNVs, and SVs within a unified evolutionary model, with long-read sequencing providing additional breakpoint resolution where available.

SVCFit estimates cellular fractions for structural variants on hemizygous chromosomes. On a male X chromosome the allele copy ratio cannot be read from heterozygous germline SNPs, so SVCFit substitutes the mean copy number per cell measured directly from read depth against the matched normal, which is itself single-copy there, and solves the resulting closed forms for each ordering of the structural variant and any overlapping copy-number change. This matters for cancers of men, where the androgen receptor and other X-linked genes are under strong selection: in metastatic castration-resistant prostate cancer the AR locus is a primary site of structural evolution, and quantifying its rearrangements alongside the rest of the genome lets these events enter the analysis on the same footing. Two limits bound the current implementation. Tandem duplications on a hemizygous chromosome are reported as upper bounds, because the number of tandem copies is not identifiable from a single locus. Short deletions can rest on read evidence alone, because a deletion smaller than the copy-number segmentation window has no depth contrast to corroborate it. Both are stated where the estimates are reported, and neither prevents the majority of chrX events from receiving a cellular fraction.

In conclusion, we demonstrate that SVCFit enables robust reconstruction of SV-defined clonal architecture from genomic data. Although shown here in metastatic prostate cancer, SVCFit is broadly applicable to other malignancies in which structural variation plays a central role. By enabling reconstruction of SV-defined clonal phylogenies from routine bulk sequencing, this framework supports more comprehensive analysis of tumor evolution and reduces the risk that clinically consequential genomic alterations are overlooked in precision oncology.

## Methods

### Closed-form SVCF in diploid regions

SVCFit estimates SVCF from breakpoint and breakend read counts obtained from whole-genome sequencing. For each SV class, SVCF is recovered from the observed breakpoint count (BPC) and breakend count (BEC) using the closed-form expressions summarized in Table 1 of Results, which follow from the SV-class-specific multiplicative identities derived by induction in Supplementary Notes S1 and S7. In diploid regions, R̅ =2 for all SV classes except tandem duplications, for which R̅ = 2(*r* − 1)/*r* where *r* is the local total copy number from FACETS.

### Allele-specific copy number from heterozygous SNPs

In non-diploid regions, SVCFit requires the local allele copy ratio (ACR) to apply the read-count adjustments in Eqs. 3-4. For each germline heterozygous SNP within an inferred copy-number segment, ACR is recovered from its variant allele frequency *VAF*_ℎ*et*_:

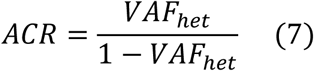

Eq. 7 introduces a directional ambiguity between alternate-allele amplification and reference-allele loss; SVCFit resolves this using the gain/loss classification from FACETS^23^. Per-segment ACR is taken as the median across the heterozygous SNPs in that segment. Full algebraic derivation is given in Supplementary Note S5.

### Haplotype-aware SV phasing and zygosity inference

Each SV is assigned to one of the two haplotypes using germline heterozygous SNPs that fall within 500 bp of an SV breakpoint. Tumor BAMs are filtered to breakpoint-supporting reads using samtools^53^ (v1.21;-f 1-F 2), and bcftools^53^mpileup is run at the heterozygous SNP positions over the filtered BAM. Presence of only one of the two heterozygous SNP alleles in the breakpoint pileup identifies a heterozygous SV on the haplotype carrying that allele; presence of both alleles identifies a homozygous SV. Overlapping CNVs are phased by the FACETS allele-specific call.

Which of the two non-diploid update rules applies depends on whether the CNV affects the SV-supporting or the reference reads, and that is fixed jointly by the cis/trans relationship and by the order of the two events: only a CNV in cis that postdates the SV affects the breakpoint reads, whereas a CNV in trans, or in cis but predating the SV, affects the breakend reads. These configurations therefore collapse to two observationally distinct cases, which SVCFit separates from the read counts rather than from the haplotype labels. The numerator of Eq. 5 is strictly positive only in the first case, so Eq. 5 is applied when it is positive and Eq. 6 otherwise; an overlapping deletion is always assigned to Eq. 6, since an SV predating a deletion of its own haplotype would leave no breakpoint reads to observe. Because this test uses only read counts, the rule is defined for every SV, including those with no informative heterozygous SNP. The haplotype assignment itself is retained for reporting; the quantities that enter the estimate are zygosity and the allele copy ratio.

Across the 11 evaluable COMBAT subjects, 61.3% of SVs (3,128 of 5,181) had at least one germline heterozygous SNP within 500 bp of a breakpoint and were assigned to a haplotype. The remaining 38.7% (1,963 of 5,181) could not be phased due to insufficient heterozygous SNP coverage within the 500 bp breakpoint window, a limitation inherent to short-read data at SV breakpoints, and take SVCFit’s default assignment rather than being dropped: each is assigned to one of the two haplotypes by convention and treated as heterozygous, so the halving applied to homozygous events is not applied to them. An overlapping CNV whose allele-specific state is uninformative receives the same default label. These labels distinguish the two haplotypes only; they carry no parent-of-origin information, which short-read tumor data without parental genotypes cannot establish.

### Iterative SVCF estimation in non-diploid regions

When an SV overlaps a CNV, the read-count adjustments in Eqs. 3-4 couple *BPC*^′^ and *BEC*^′^ to SVCF itself. SVCFit therefore applies a fixed-point iteration initialized at *SVCF*_0_ = 0.5. When the breakpoint count is altered (CNV in cis occurring after the SV), the update rule is *R* ⋅ *BPC* ⋅ *α_q_ SVCF_q_*

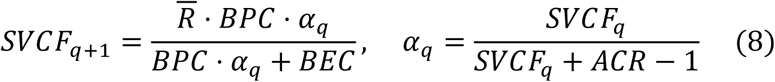

When the breakend count is altered (CNV preceding an in-cis SV, or CNV in trans), the update rule is

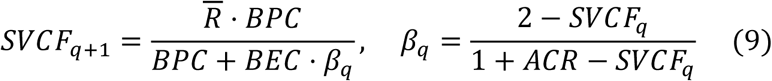

Iteration proceeds until |*SVCF_q_*_+1_ − *SVCF_q_*| < 10^−6^ or 100 iterations, whichever comes first. Both procedures converge to the closed-form fixed points given by Eqs. 5 and 6 of Results; convergence proofs and the formal derivation of the SV-CNV ordering criterion (sign of Eq. 5) appear in Supplementary Note S2.

### Sequencing-depth assumption

SVCFit assumes that average read depth at SV breakpoints is approximately equal to the average read depth at the corresponding breakends (Eq. S15, Supplementary Note S1). This was empirically validated in 1,000 simulated clonal tumors at 50× coverage spanning SV tumor purities from 10% to 100% (Supplementary Fig. S1; Pearson correlation ≈ 0; analytical justification in Supplementary Notes S6 and S8).

### Synthetic benchmark sets (VISOR)

VISOR was used to build three independent synthetic datasets: the 75-scenario autosomal accuracy benchmark described in this subsection, a separate hemizygous-chromosome accuracy benchmark on chromosomes 22 and X (described below), and a set of paired pre-and on-treatment genomes used to evaluate the downstream phylogenetic inferences, whose design is given under “Longitudinal self-evaluation simulations” below. For the accuracy benchmark we used the Haplotype-Aware Structural Variants Simulator VISOR^24^ (v1.1.3) to generate multi-clonal synthetic tumor genomes (with one matched normal). Somatic structural variants (SVs) with known structural-variant cellular fractions (SVCFs) and copy-number variants (CNVs) were spiked into tumor genomes to explicitly model SV-CNV overlaps in both cis and trans configurations. To optimize compute, this autosomal benchmark was restricted to chromosomes 1 and 2, spanning a range of SV/CNV overlap contexts, tumor purities, and subclonal compositions; the hemizygous benchmark instead used chromosomes 22 and X (below).

For each scenario, we specified a phylogeny with a germline root and a clonal tumor ancestor that bifurcates into two terminal subclones. SVs were simulated on chromosome 1, ordered from the p-to q-arm, and partitioned into three sets assigned to the truncal clone and two subclones (33/33/34 events, respectively). Chromosome 2 was used only as a partner for inter-chromosomal translocations. Restricting the simulation to these chromosomes simplified the controlled design of SV-CNV overlap patterns across tumor purities and subclonal mixtures.

Next, we simulated five SV-CNV configurations: (i) SV without CNV; (ii) SV preceding cis amplification; (iii) SV preceding trans amplification; (iv) SV following cis amplification; and (v) SV following trans deletion. Each configuration was crossed with five SV tumor-purity levels (10%, 20%, 40%, 60%, 80%; Eq. 10) and three subclonal mixture settings (10:90, 30:70, 50:50), yielding 75 scenarios at 50× mean coverage (one BAM per scenario). Each scenario was simulated as a single multi-clonal genome with 100 spiked-in SVs partitioned among the truncal clone and two terminal subclones (33 / 33 / 34 events, respectively), so that observations from SVs within the same simulated genome are not statistically independent; per-condition statistical comparisons between methods therefore treat each simulated genome as the unit of observation rather than each individual SV, and Supplementary Fig. S3 panels stratify the 75 scenarios by SV class for readability rather than presenting all panels for every SV class at every condition. For each spiked-in SV we retained start/end positions, SV type, and reference/alternate read counts in both tumor and normal BAMs, enabling direct comparison with SVCFit and SVclone^21^.

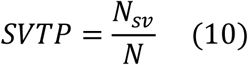

where *SVTP* is SV tumor purity, *N_sv_* is the number of cells containing any somatic SV, and *N* is the total number of cells in the sample.

### Tumor purity terminology

In this study, we distinguish between two related but distinct concepts of purity. *SV tumor purity (SVTP)* refers to the fraction of cells in a sample harboring somatic structural variants, the primary quantity modeled and estimated by SVCFit. In contrast, *tumor purity* (or *sample purity*) refers to the fraction of tumor cells in a sample, as estimated by copy-number-based methods such as FACETS^23^. Unless otherwise specified, references to “SV tumor purity” explicitly denote SVTP, whereas references to “tumor purity” denote conventional tumor cellularity.

### Prostate metastasis mixtures (benchmark set)

For direct comparison with SVclone, we generated eleven multi-clonal tumors by mixing BAM files from two single-clone metastatic prostate cancer samples. These samples were originally sequenced in a landmark study of subclonal structure in prostate cancer metastasis^22^ and were subsequently used as a key benchmark in the SVclone paper^21^. Raw tumor and matched-normal FASTQs were obtained from EGA (EGAD00001001343)^22^ and aligned to hs37d5 with BWA mem^54^ (v0.7.19) to generate a single-clone tumor BAM. Then, each single-clone tumor BAM was downsampled in 10% increments of clonal frequency and mixed to create BAMs of heterogeneous samples with known SVCFs and known clonal composition.

Nine mixtures yielded three-cluster tumors (cluster 1: SVs shared by both tumors; cluster 2: shared + private SVs from tumor A; cluster 3: shared + private SVs from tumor B). Four-and five-cluster mixtures were generated by subsampling odd and even chromosomes from each metastasis at different rates prior to mixing, as described in Cmero et al^21^. Specifically, the four-cluster mixture combined bM odd / bM even chromosomes at 20% / 60% with gM odd at 40%, yielding ground-truth CCFs of 0.2 (bM-odd), 0.6 (bM-even), 0.4 (gM-odd), and 1.0 (shared clonal); the five-cluster mixture combined bM odd / bM even at 80% / 60% with gM odd / gM even at 20% / 40%, yielding ground-truth CCFs of 0.8, 0.6, 0.2, 0.4, and 1.0 across the four sample-specific subclones plus the shared clonal cluster. Given that purity, mixture proportions, and per-SV clonality are all known quantities, and that SV tumor purity can be approximated as the purity, the true SVCF can be derived for each SV. For clonal SVs, true SVCF equals the mixture’s SV tumor purity (SVTP); for subclonal SVs, true SVCF equals the source tumor’s SVTP times its mixing proportion.

SVs were called with Manta^50^ (v1.6.0) and genotyped with SVtyper^25^ (v0.7.1). For direct comparison with SVclone, SVCF was converted to cancer cell fraction (CCF) via Eq. 11.

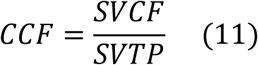

We regenerated the mixture BAM files using the same script described in the SVclone paper(https://github.com/mcmero/SVclone_Rmarkdown/tree/master/make_insilico_mixt ures.sh). We also contacted the first and senior authors to request the original mixture BAM files but did not receive a response.

mCRPC subject samples (COMBAT). The COMBAT trial, registered on ClinicalTrials.gov (NCT03554317; first posted 13 June 2018), tested bipolar androgen therapy (BAT; testosterone cypionate 400 mg intramuscularly every 28 days) for 3 months followed by BAT plus nivolumab in patients with metastatic castration-resistant prostate cancer (mCRPC) who had progressed on at least one prior androgen receptor pathway inhibitor. The trial was approved by the Johns Hopkins Medicine Institutional Review Board (IRB00164973). The study design and conduct complied with all relevant regulations regarding the use of human study participants and were conducted in accordance with the Declaration of Helsinki. Written informed consent was obtained from all participants prior to enrollment. Full eligibility criteria, response criteria, and clinical results are described in Markowski et al^30^. Participants underwent paired biopsies of a soft-tissue metastasis at screening (baseline) and after 3 months of BAT monotherapy (prior to nivolumab addition). Flash-frozen biopsies underwent laser-capture microdissection to enrich for cancer cells, followed by paired-end WGS on a NovaSeq 6000 S4 (2×150 bp; ∼110 Gb per sample) for each biopsy and matched PBMC normal. FASTQs were trimmed with trim_galore^55^ (v0.6.10), aligned to GRCh38 with BWA mem^54^ (v0.7.19), and processed with GATK4^56^ (MarkDuplicates, BaseRecalibrator, ApplyBQSR; v4.6.2.0).

### Somatic structural variant calling

For the VISOR accuracy benchmark and the prostate-mixture benchmark, somatic SVs were called with Manta^50^ (v1.6.0) using tumor-normal pairs with default parameters. For COMBAT, somatic SVs were called using Manta (v1.6.0), Delly^51^ (v1.5.0), and GRIDSS^52^ (v2.13.2) with default parameters, each provided with the tumor BAM and its matched normal. For SV genotyping we used SVtyper^25^ (v0.7.1), which was selected to decouple SV discovery from breakpoint-resolved read-evidence quantification.

Modern somatic SV pipelines achieve their best precision-recall by combining multiple discovery callers: Manta, Delly, and GRIDSS each exploit complementary evidence (assembly-based, paired-end + split-read, and breakend-assembly-with-rule-based-filtering, respectively), and consensus across two or more of these callers consistently outperforms any single tool, but the discovery callers report inconsistent and non-comparable read-support metrics, making their raw output unsuitable for VAF-based downstream analyses. SVtyper provides a uniform, per-breakpoint quantification using a Bayesian likelihood model over discordant paired-end and split-read evidence, the same evidence classes used by the discovery callers themselves, producing per-SV reference-supporting (RO), alternate-supporting (AO), and allele-balance (AB) values directly grounded in breakpoint geometry. These values are the per-SV reference/alternate counts required as input by Eq. 1 of the main text and the per-class identities in Supplementary Note S1. The Manta + Delly + GRIDSS → SURVIVOR^27^ → SVtyper workflow we adopt for the COMBAT cohort is an established pattern for cancer somatic SV analysis, used for example in the *BCL11B* enhancer-hijacking study in lineage-ambiguous stem-cell leukemia^57^. Alternative breakpoint genotypers, caller-specific built-in genotypers, or graph-based germline-population genotypers, are restricted to one caller’s VCF format or are optimized for population-scale germline rather than per-sample somatic VAF.

Because SVtyper expects a LUMPY-formatted VCF with INFO fields MATEID, END, CIPOS, CIEND, and SVTYPE, we manually appended CIPOS and CIEND to the VCF outputs from GRIDSS, Delly, and Manta to ensure compatibility. Finally, for COMBAT, all genotyped VCF files were harmonized using SURVIVOR^27^ (v1.0.7) to produce a consensus VCF prior to cellular fraction analysis.

### Rescue of key driver events

Using default parameters, SVtyper detected the canonical prostate cancer fusion *TMPRSS2-ERG*^12,13^ in the pre-BAT biopsy of COMBAT Subject 10 (tumor purity 68%; breakpoint coverage 95), but not in the on-BAT biopsy, which had substantially lower tumor purity and reduced read depth at the breakpoint (purity 36.6%; breakpoint coverage 31). Given that *TMPRSS2-ERG* is a well-established early event in prostate tumorigenesis and is rarely lost, we manually inspected the on-BAT BAM file and identified a single read supporting the *TMPRSS2-ERG* fusion (deletion at chr21:38,499,931-41,501,273). While this read alone is insufficient to confidently call the fusion, ERG protein expression was positive by immunohistochemistry in both the pre-and on-BAT biopsies, a pattern strongly suggestive of an underlying *ERG* rearrangement, most commonly *TMPRSS2-ERG*. Together with its robust detection by WGS in the pre-BAT sample, these orthogonal data support *TMPRSS2-ERG* as a clonal event, and we therefore placed it on the trunk of the inferred phylogeny. Similarly, TMPRSS2-ERG was also rescued for subjects 1, 6, 12, 13.

In subject 20, PTEN del and RAD51B inv were initially filtered out because it failed the 2 called criteria but both SVs were detected in pre-and on-BAT samples using delly and were recovered. In addition, PTEN loss is confirmed to be present at both pre-and on-BAT sample via IHC data.

The same principle was applied to two point mutations. In Subject 20, the AR ligand-binding-domain mutations p.Leu702His (chrX:67,711,621 T>A) and p.Thr878Ala (chrX:67,723,710 A>G) were called by MuTect2 but removed by its germline-risk filter, which fires on membership in dbSNP combined with a low normal-sample likelihood ratio; the matched PBMC normal was covered at only 6 and 13 reads at these two positions. Both are recorded as somatic prostate-cancer variants in ClinVar and COSMIC. Because AR lies on the X chromosome and these are male patients, a germline variant would be present at essentially the full allele fraction in the matched normal, whereas the normal carried 6 reference and 0 alternate reads at p.Leu702His and 13 reference and 0 alternate reads at p.Thr878Ala, excluding a germline origin.

In the tumor BAMs the supporting reads were at mapping quality 60, predominantly properly paired and balanced across strands, free of soft-clipping, with a median edit distance of 1 and mates mapping to the X chromosome. Median read depth across the AR gene body matched the X-chromosome background (13 versus 11 pre-BAT; 10 versus 9 on-BAT) against an autosomal median of 28 and 19, so the locus is single-copy with no focal amplification. We therefore recovered both mutations and genotyped them from the BAMs at both timepoints: p.Leu702His at 15 of 15 reads pre-BAT and 9 of 10 on-BAT, p.Thr878Ala at 6 of 6 pre-BAT and 2 of 3 on-BAT. PICTograph assigns both to the truncal cluster.

### Germline heterozygous SNP detection

Germline SNPs were identified from matched normal BAMs using GATK4^56^ HaplotypeCaller (v4.6.2.0; default parameters) and filtered to heterozygous sites using bcftools^53^ (v1.20). The downstream use of these heterozygous SNPs for SV phasing and ACR inference is described in the “Structural variant cellular fraction estimation” subsection above.

### Somatic copy-number variant detection

CNVs overlapping SVs were called with FACETS^23^ (v0.6.2). FACETS first generates a SNP pileup file using tumor and matched-normal BAMs and SNPs detected from GATK4^56^ HaplotypeCaller (v4.6.2.0; default parameters); then coverage and the germline B-allele frequencies (BAFs) were computed from the SNPs pileup to detect CNV. FACETS SNP pileups were generated with:-g-q15-Q20-P100-r25,0.

### Dependence on upstream SV calling

SVCFit consumes SV breakpoint coordinates and per-SV reference-and alternate-supporting read counts produced by upstream SV calling and breakpoint genotyping; it does not itself perform SV discovery. Different SV callers (Manta^50^, Delly^51^, GRIDSS^52^, and others) differ in breakpoint-detection sensitivity, in their definitions of split-read versus discordant-pair support, and in their internal quality filters. These differences propagate into the per-SV reference-and alternate-supporting read counts that feed Eq. 1 and the per-class identities of Supplementary Note S1, and can therefore influence downstream SVCF estimates. To mitigate caller-specific idiosyncrasies in the COMBAT analysis we used a multi-caller consensus (Manta, Delly, GRIDSS) merged with SURVIVOR^27^ before genotyping; for the VISOR and prostate-mixture benchmarks, SVs were called with Manta only to match the SVclone benchmark configuration. We discuss the implications of caller dependence further in the Discussion.

### SVCF comparison and benchmarking

SVCFit requires inputs including: SV positions and types, SV read counts (SVtyper), CNV data (FACETS), and germline heterozygous SNP pileups (GATK4 and bcftools). SVCF per SV was computed as defined in the Results (Eq. 1; Table 1) with non-diploid corrections from Eqs. 3-6.

SVclone (v1.1.2) was obtained from on March 10, 2024. SVclone was run with default parameters using as input the SV genomic coordinates, VISOR-specified tumor purity and ploidy from the simulations, and the tumor BAM file.

### Benchmarking on VISOR simulations

For each of the 75 VISOR simulation scenarios, structural variants (SVs) were identified using Manta and subsequently genotyped with SVtyper to obtain reference and alternate supporting read counts. Heterozygous single-nucleotide variants (SNVs) were detected using GATK4 HaplotypeCaller and further processed with bcftools. Copy-number variants (CNVs) were inferred with FACETS. Additionally, tumor purity and ploidy estimates from FACETS, along with the simulated mean sequencing depth (50×), were supplied as input to SVclone. For SVclone, per-SV cancer cell fractions in copy-number-altered regions were resolved using the ground-truth simulated cellular fraction to disambiguate SV multiplicity, following the SVclone authors’ evaluation convention; SVCFit’s estimates used no ground-truth information. This assist applies only to the four configurations with an overlapping copy-number event and should be considered when interpreting those comparisons. To quantify uncertainty in all performance metrics, we generated 30 independent bootstrap replicates of each scenario by resampling reads with different random seeds and ran the full SVCFit pipeline (Manta, SVtyper, GATK HaplotypeCaller, FACETS) and SVclone independently on each replicate. The ground-truth SVCF values derived from the simulation were compared to estimates produced by SVCFit and SVclone. Error was defined as:

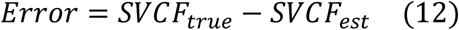

Between-method accuracy was summarized by each method’s mean signed error with a bootstrap 95% confidence interval across the 30 replicates, and by the paired effect size, the difference in absolute mean signed error (|SVCFit| minus |SVclone|), with its bootstrap 95% confidence interval. We also computed a two-sided paired Wilcoxon signed-rank test on the per-replicate errors with Benjamini-Hochberg correction; because each replicate averages thousands of SVs, this replicate-level test saturates near its 30-replicate floor for essentially every condition and is therefore uninformative about effect magnitude, so significance is not marked on the figures and the effect sizes, confidence intervals, and adjusted p-values are reported in companion statistics tables instead.

### Benchmarking on a hemizygous chromosome

The hemizygous estimators were evaluated on a separate VISOR simulation of a male genome, modeled with two haplotypes that both carry chromosome 22 and only one of which carries chromosome X, so that chromosome X is single-copy and chromosome 22 serves as a diploid copy-neutral control. Somatic deletions, inversions, and tandem duplications were spiked onto chromosome X under three SV-CNV configurations, the SV without an overlapping copy-number change and the SV arising before or after a copy-number amplification, each crossed with the same five tumor purities (10-80%) and three subclonal mixtures (10:90, 30:70, 50:50) as the autosomal benchmark, giving 45 conditions at 50× mean coverage and 30 replicates each (4,935 ground-truth SVs). Because FACETS is unusable on a hemizygous chromosome, copy number was obtained by DNAcopy circular binary segmentation of read depth, and both methods received the same variant calls and the same segmentation; SVclone was additionally supplied the true tumor purity, which SVCFit does not use. Hemizygous tandem duplications are reported by SVCFit as upper bounds and excluded from the error statistics. Confidence intervals were obtained by a block bootstrap over replicates (B = 10,000, percentile intervals), paired within each draw so that both methods are evaluated on the same replicate in every draw. These intervals resample tumor read noise only: every replicate shares one matched normal, so intervals on quantities that depend on the tumor-to-normal depth ratio are narrower than a full resampling would give.

### Benchmarking on prostate metastasis mixtures

SVs were called with Manta and genotyped with SVtyper on the mixture BAMs. SVs were filtered following Cmero et al.^21^: fewer than one supporting read in the matched normal, more than two supporting reads in at least one tumor sample, CCF capped at 1.0, and PASS calls only. SVCF estimates from SVCFit were converted to cancer cell fraction (CCF) using Eq. 11 (Methods), with tumor purity estimated by FACETS for each mixture. Error was defined as *Error* = *CCF_true_* − *CCF_est_* (positive values indicate underestimation), where the true CCF is computed from the known mixing proportions. To quantify uncertainty, we generated 30 independent replicates of each of the eleven mixture conditions by subsampling reads with different random seeds and ran the full SVCFit pipeline (Manta, SVtyper, GATK HaplotypeCaller, FACETS) independently on each replicate.

SVclone was run on the same eleven non-subsampled mixtures, but supplied with the same FACETS copy-number calls used by SVCFit rather than its default Battenberg copy number, so that both methods received the same FACETS copy-number and purity estimates; per-SV CCF was then computed analytically from the observed read counts, sample purity, and these copy-number genotypes using MapVaf2CcfPyClone, following the procedure that produced the SVclone publication’s Figure 2C. In this convention, SVclone resolves each SV’s multiplicity using the ground-truth simulated cellular fraction, an assist that SVCFit does not receive; this anchors its estimate to the true value and introduces a consistent positive offset in the computed CCF that is independent of SVclone’s subclonal-reconstruction quality, which we take into account when interpreting the SVclone comparison in this benchmark.

For each SVCFit replicate-SVclone pairing, mean signed CCF error was computed per mixture condition. Per-condition differences between methods were tested using a two-sided paired Wilcoxon signed-rank test on the 30 paired replicate mean errors, with Benjamini-Hochberg correction across the eleven mixture conditions. Unlike the VISOR benchmark, where each replicate averages thousands of simulated SVs and this test saturates for essentially every condition, the mixture errors are computed over the limited set of shared prostate SVs and several conditions carry a near-zero true difference between methods; the test is therefore not saturated here, and its adjusted p-values are informative (three of the eleven conditions are non-significant). To compare the two methods on the same variants, errors were computed on the intersection of SVs detected and truth-matched by both methods; results on each method’s full call set are also reported in Results.

### Longitudinal self-evaluation simulations

To evaluate downstream phylogenetic inferences, we generated paired pre-and post-treatment whole-genome simulations with VISOR^24^ following a single S1 scenario: a truncal clone (CCF = 1.0 at both timepoints) and two subclonal children whose CCFs cross under simulated therapeutic selection (one declining from CCF 0.83 → 0.17 and one expanding from 0.17 → 0.83). Each replicate carried 33 truncal SVs and 33 + 34 subclonal SVs on the two terminal subclones, drawn from a mixture of deletions, inversions, tandem duplications, and translocations, with reads simulated at 50× mean coverage and a matched normal at 50×. SV tumor purity was swept over {20%, 40%, 60%, 80%} with 25 paired pre/post replicates per purity (boots 0-4 × five experimental designs; n = 24 at 40% purity due to one failed to build tree).

The full SVCFit pipeline (BWA-MEM^54^ v0.7.19 alignment to GRCh38, Manta^50^ v1.6.0 SV calling, SVtyper^25^ v0.7.1 genotyping, GATK4^56^ v4.6.2.0 HaplotypeCaller germline SNP calling, FACETS^23^ v0.6.2 copy number inference) was run independently on each replicate at default SVCFit parameters with no per-scenario tuning (DP-GMM concentration α = 1).

For each replicate we computed three metrics: per-SV truncal/subclonal balanced accuracy (the mean of sensitivity and specificity) and F1 (truncal = positive class) at each timepoint, per-clone signed CCF error (true − inferred CCF, positive = underestimation), and exact rooted-topology recovery against the S1 ground-truth tree. Per-purity summaries report means, with bootstrap 95% confidence intervals for balanced accuracy, CCF error and topology recovery. Balanced accuracy, F1 and topology recovery use all 24-25 replicates per purity; CCF error uses the 20 to 22 replicates per purity that yielded clone CCFs. Full per-purity numerical results, the multi-covariate analysis of per-clone signed CCF residuals, and the operating-range conclusions are presented in Supplementary Note S3.

### SV Longitudinal Clustering and Dynamics

Structural variants (SVs) in the COMBAT dataset were clustered on a per-subject basis using Dirichlet Process Gaussian Mixture Models (DP-GMM). DP-GMM inference was performed using the scikit-learn library^58^ in R via the reticulate interface^59^. For clustering, all SVCF was converted to CCF using purity estimates from FACETS^23^ (Eq. 11). SVs detected at only one time point were assigned a CCF of 0 at the other time point. To avoid bias in DP-GMM clustering, we retained only the unique CCF pair observed across the two time points. Two features were then derived for clustering: the pre-BAT CCF and the change in CCF (ΔCCF = on-BAT CCF − pre-BAT CCF). DP-GMM models were fitted using a maximum of 10 components (kmax = 10), a Dirichlet process concentration parameter of 1 (weight_concentration_prior = 1), and a covariance regularization term of reg_covar = 1e−6, covariance_type = “full”, and init_params = “k-means++”. Both cluster-merging cutoffs are motivated as follows.

Three complementary considerations justify this floor. First, *algorithmic stability under 2D dimensionality*: SVs are clustered jointly in the two-dimensional space of (pre-BAT CCF, ΔCCF), so the mathematical minimum for a non-singular empirical covariance matrix is N = D + 1 = 3 non-collinear points; furthermore, a full 2×2 covariance matrix requires estimating three parameters (σ^2^_pre, σ^2^_Δ, σ_pre,Δ) rather than one, substantially increasing susceptibility to overfitting at low N. At 3-4 points, the DP-GMM Bayesian prior overwhelmingly dictates covariance matrix orientation, producing ill-conditioned estimates and triggering aggressive component pruning during Expectation-Maximization (EM); five points is the absolute bare minimum at which 2D covariance estimation is statistically feasible without numerical failure. Second, *biological signal vs. technical noise in multi-sample data*: SV calling carries inherent noise from sequencing artifacts, mapping anomalies, and read-depth fluctuations; in 2D clustering, spurious artifact calls at similar CCF across both timepoints can masquerade as subclonal clusters. A putative subclone supported by fewer than five SVs lacks the evidentiary footprint to be distinguished from multi-dimensional background noise. Third, *consistency with published practice*: minimum-variant-per-cluster thresholds of 5-10 are standard filtering heuristics across published tumor subclonal-reconstruction pipelines^60,61^; our floor of 5 in 2D is consistent with this convention, representing the conservative (lower) end of the established range. Clusters falling below five SVs were merged with their nearest neighbor by Euclidean distance.

The second cutoff merges clusters that are geometrically near-coincident. For each subject, SVCFit computes the pairwise Euclidean distance between cluster centres in the two-dimensional (pre-BAT CCF, on-BAT CCF) space and combines any two clusters whose centre-to-centre distance is below 0.2. This consolidates the near-duplicate splits that DP-GMM can produce when SV-type-dependent CCF noise fragments a single true subclone into two centres separated by only a small distance, while leaving genuine subclones, whose centres are separated well beyond 0.2 in this space, as distinct clusters. Both cutoffs were fixed prior to formal benchmarking and were not tuned per scenario. Alluvial plots were constructed using the R package ‘ggalluvial’^62^. Circos plots were constructed using the R package ‘circlize’^63^. Clonal trees and pie plots were created with SVCFit software.

### SV Tumor Evolution Tree Construction and Annotation

SV clusters were ordered onto the edges of an evolutionary tree. Candidate directed edges between clusters were first identified by enforcing lineage precedence, which required that descendant CCFs not exceed ancestral CCFs by more than a tolerance ε1. Valid edges were assembled into candidate trees using a modified Gabow-Myers algorithm^64^, following the pseudocode of Popic et al.^65^. Briefly, the original Gabow-Myers procedure enumerated all directed spanning trees of a directed graph rooted at the ancestral node, and the modification constrained enumeration to edges that satisfied the lineage-precedence filter. Tree topologies that violated the Sum Condition^66^, where the sum of descendant CCFs exceeded the ancestral CCF by more than ε2, were excluded. We used ε1=0.2 and ε2=0.2. The tree reconstruction approach did not assume clock-like evolution or edge lengths proportional to time.

The remaining trees were ranked using the SCHISM fitness function^67^, which assigned penalties for violations of lineage precedence and the Sum Condition. These penalties were combined as a weighted sum within the exponent of a negative exponential scoring function, such that trees with fewer or smaller violations received higher fitness scores. Equal weights were assigned to the two violation terms (weights = 1).

Per-sample clonal proportions were then inferred from the top-ranked tree, where the proportion of a parent clone was computed as its CCF minus the sum of the CCFs of its children:

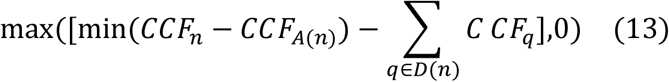

where *CCF_n_* is the CCF of the cluster associated with node n. *CCF_A_*_(*n*)_ is the CCF of the cluster directly upstream of node n. ∑*_q_*_∈*D*(*n*)_ *C CF_q_* is the sum of CCFs of clusters directly downstream of node n.

To annotate the edges of the trees, we first identified a set of prostate cancer driver genes and their hg38 coordinates from Hosseini et al.^37^ and web search. Custom R scripts were used to identify driver gene overlap with the coordinates of the SVs called in each subject, including nearby upstream regions. Literature support for the driver-status of each gene that appears on a tree edge in Fig. 5 or in Supplementary Note S4 is compiled in Supplementary Table S2, organized by pathway with primary references. When annotating tree edges, “deletions” and “duplications” refer to events defined from SV breakpoint evidence rather than read-depth copy-number calls. Hemizygous deletion is annotated only when additionally supported by copy-number evidence (CN=1), and intragenic duplication/deletion only when breakpoints fall within the gene body.

To control for differences in mean sequencing depth between the pre-BAT (36×) and on-BAT (17×) biopsies from Subject 10, we performed a down-sampling validation. The pre-BAT BAM file was subsampled using *samtools view-s* to reduce the total read count so that the effective tumor coverage matched that of the on-BAT sample. Structural variant calling was re-run on this down-sampled dataset using the identical pipeline. To quantify sampling variability, the downsampling was repeated across 20 bootstrap replicates using distinct random seeds; mean SV counts and 95% CIs across replicates are reported in the Results.

### SNV-based subclonal reconstruction

For a data-type-matched comparison with the SV-based phylogenies, we reconstructed SNV-based subclonal phylogenies for each COMBAT subject using PICTograph^19^ (Zheng et al.; v1.3.2). The somatic single-nucleotide variants were the COMBAT trial’s own published calls (GATK MuTect2; Markowski et al.^30^), accepted as reported without re-calling. The alternate and reference read counts for each SNV at the pre-and on-BAT timepoints were taken directly from that output and provided to PICTograph as input. Two variants were added beyond the trial’s PASS calls, both in Subject 20: the AR ligand-binding-domain mutations p.Leu702His and p.Thr878Ala, which the caller’s germline-risk filter had removed. Their reference and alternate read counts were taken from the aligned BAMs at both timepoints. The rationale and the supporting read-level evidence are given under “Rescue of key driver events”. PICTograph performs multi-sample joint inference across both timepoints in a single reconstruction, matching the multi-sample design of the SVCFit analysis. PICTograph and SVCFit share the same modified Gabow-Myers tree-construction and tree-scoring approach, so the SV-and SNV-based phylogenies differ only in the input variant class and its clustering. Clustered SNVs were annotated against the same prostate-cancer driver gene list used for the SV clusters (Supplementary Table S2). The paired per-subject SNV-and SV-based phylogenies are shown in Supplementary Note S4.

### IHC analysis

ERG IHC was performed on an automated staining platform (Ventana/Roche) using a validated IHC assay as described previously^68^ using formalin-fixed, paraffin-embedded tumor sections were that were taken adjacent to the fresh frozen tissue used for the whole genome sequencing.

## Statistical analysis

Unless stated otherwise, analyses used the two-sided paired Wilcoxon signed-rank test (α = 0.05). All p-values are corrected for multiple testing using the Benjamini-Hochberg method, with significance assessed against BH-adjusted p-values. Medians and interquartile ranges were reported where appropriate. The function wilcox_test from the R package rstatix (v0.7.2)^69^ was used to apply the statistical test.

## Data availability

All open access data is accessible through Mendeley (10.17632/2nhhdjx225.6)

## Code Availability

https://github.com/KarchinLab/SVCFit/ (released under the MIT license)

## Acknowledgements

This work was supported by The Prostate Cancer Foundation, the Lustgarten Foundation (Y.L. L.D.W., R.K.), CDMRP/DOD grant W81XWH-21-PCARP-IDA (Y.L. L.D.W., R.K.), and NIH-NCI grants U01CA271273 (J.L., L.D.W., R.K.), P30CA006973 (S.Y, A.M.D.M) and P50CA272391 (S.Y., A.M.D.M), and Break Through Cancer, Cambridge, MA (to R.K., L.D.W., J.L.). E.S.A. is partially supported by NCI grant P30 CA077598 and DOD grant W81XWH-22-2-0025. M.C.M is supported by DOD grant W81XWH-19-1-0692.

Special thanks to Dr. Winston Timp and Dr. Benjamin Langmead for their support of this project in its early days.

## Author Contributions

Y.L. and R.K. conceived the study. Y.L. and R.K. developed the methodology. Y.L. implemented the software and performed the computational analyses. J.L. and Y.Y. contributed to data analysis and figure generation. M.C.M., E.S.A., A.M.D.M., S.Y., L.D.W., and L.A.S. provided clinical interpretation, biological insight, and domain expertise. L.A.S. coordinated access to clinical data and resources. Y.L., L.A.S., and R.K. wrote the initial manuscript. All authors contributed to manuscript review and editing. R.K. supervised the study.

