## Supplementary Material for "Longitudinal structural variant phylogenies define tumor evolution under therapeutic selection pressure in metastatic prostate cancer"

#### Supplementary Information

Yunzhou Liu, Jiaying Lai, Yi Yang, Mark C. Markowski, Emmanuel S. Antonarakis, Angelo M. De Marzo, Srinivasan Yegnasubramanian, Laura D. Wood, Laura Sena*, Rachel Karchin†

#### Table of Contents

- Supplementary Note S1: Structural variant cellular fraction (SVCF) definition, full derivations
- Supplementary Note S2: Algorithm convergence and SV-CNV ordering, convergence proofs
- Supplementary Note S3: Longitudinal self-evaluation on paired pre/post-treatment simulations
- Supplementary Note S4: Cohort-wide SV-based vs SNV-based clonal phylogenies in COMBAT
- Supplementary Note S5: SVCF estimation in non-diploid genomic regions, full derivations
- Supplementary Note S6: Robustness of SVCF estimation to breakpoint read depth
- Supplementary Note S7: Mathematical properties of SVCF under tandem duplications and deletions, induction proofs
- Supplementary Note S8: SV breakpoint and breakend read depth across tumor purities
- Supplementary Figures S1-S16 (Figure S14 comprises the eleven per-subject panels S14-1 to S14-11)
- Supplementary Tables: S1 (bioinformatics tools, versions and parameters); S2 (driver-gene citation table); S3-1 and S3-2 (per-purity longitudinal self-evaluation results, Note S3); S4-1 (per-subject SV-tree vs SNV-tree summaries, Note S4); S7-1 (tandem-duplication variables, Note S7)

#### Where in the main text

Each Supplementary Note backs up specific content in the main text. This table maps each Note to its corresponding main-text section and equations.

| Note | Topic | Main-text location | Main-text eq(s) |
|---|---|---|---|
| S1 | SVCF definition; per-class derivations; multiplicative identities | Results, “The SVCFit framework”; Methods, “Closed-form SVCF in diploid regions” | Eq. 1; Table 1 |
| S2 | Convergence proofs for the two iterative update rules (S2.1, S2.2); SV-CNV ordering criterion proof (S2.3) | Methods, “Iterative SVCF estimation in non-diploid regions”; closing paragraph of Results, “Estimation in non-diploid genomic regions” | Eqs. 5, 6, 8, 9 |
| S3 | Longitudinal self-evaluation: paired pre/post-treatment VISOR[1] simulations across SV tumor purities 20-80%; per-SV truncal/subclonal classification, per-clone CCF estimation, and rooted tree-topology recovery; multi-covariate analysis of per-clone signed CCF residuals | Results, “Longitudinal self-evaluation on paired pre/post-treatment simulations” | (no main-text equation) |
| S4 | Cohort-wide SV-based vs SNV-based clonal phylogenies for the 11 evaluable COMBAT subjects; per-subject summaries for the 9 non-featured subjects; aggregate SV-vs-SNV comparison and recurrent resistance/sensitivity-determinant tables | Results, “Longitudinal analysis of SV-defined clones in mCRPC subjects receiving bipolar androgen therapy” (cohort-framing paragraph and cohort-summary paragraph) | (no main-text equation) |
| S5 | ACR inference; non-diploid read-count adjustments; closed-form fixed points; SV/CNV ordering | Results, “Estimation in non-diploid genomic regions”; Methods, “Allele-specific copy number from heterozygous | Eqs. 3, 4, 5, 6, 7 |

| Note | Topic | Main-text location | Main-text eq(s) |
|---|---|---|---|
| | | SNPs", "Iterative SVCF estimation in non-diploid regions" | |
| S6 | Empirical validation that breakpoint and breakend read depths are approximately equal across purity (1,000 VISOR[1] simulations) | Methods, "Sequencing-depth assumption"; Supplementary Fig. S1 | (no main-text equation) |
| S7 | Tandem-duplication and deletion multiplicative-identity induction proofs; derivation of $\overline{R} = 2(r-1)/r$ | Methods, "Closed-form SVCF in diploid regions" (Table 1 entry for tandem duplications) | Table 1 |
| S8 | Analytical justification of breakpoint vs breakend read-depth equality across purities | Methods, "Sequencing-depth assumption" | (no main-text equation) |

| Supplementary figure | Subject | Referenced from |
|---|---|---|
| Fig. S1 | Read depth stability across SV tumor purities | Methods, "Sequencing-depth assumption"; Note S6 |
| Fig. S3 | Per-condition benchmarking error (SVCFit vs SVclone) across SV-CNV overlap configurations × SV tumor purity | Results, "SVCFit improves SV cellular fraction estimation…" paragraph |
| Fig. S13 | Genome-wide SVs in pre- and on-BAT samples for Subjects 10 and 31 | Results, COMBAT downsampling discussion |
| Fig. S2 | SV-CNV ordering schematic | Note S2.3; closing paragraph of Results, "Estimation in non-diploid genomic regions" |
| Fig. S8 | Operating range of SVCFit's downstream phylogenetic inferences across the 20-80% purity gradient (per-SV balanced accuracy; per-clone | Results, "Longitudinal self-evaluation…"; Note S3 |

| Supplementary figure | Subject | Referenced from |
|---|---|---|
| | signed CCF error; rooted-topology recovery) | |
| Fig. S9 | Illustrative recovered topology and paired pre/post centroid placement at 60% purity | Results, “Longitudinal self-evaluation…”; Note S3 |
| Fig. S10 | Per-clone signed CCF error vs. number of SVs assigned to cluster, stratified by clone identity | Results, “Longitudinal self-evaluation…”; Note S3 |
| Fig. S11 | Multi-covariate Spearman ρ heatmap (4 covariates vs per-clone signed CCF error, stratified by clone × SV type); N SVs × BND cells localize the BND- mappability contribution | Results, “Longitudinal self-evaluation…”; Note S3 |

#### Supplementary Note S1: Structural variant cellular fraction (SVCF) definition

**Cross-reference to main text.** The summary relationship $SVCF = \overline{R} \times VAF$ appears as Eq. 1 of the main text, and the per-class closed forms summarized below (Eqs. S26-S53) are tabulated in Table 1 of the main text. The VAF definition of Eq S9 appears as Eq. 2 of the main text, with BPRC and BERC written there as SV-supporting and reference reads. This Note retains the full step-by-step derivation.

This note formally defines the structural variant cellular fraction (SVCF), which quantifies the fraction of cells in a bulk tumor sample that harbor a given structural variant. We derive SVCF from observable quantities in whole-genome sequencing data, including breakpoint read counts, breakend read counts, and average read depth, and show how these quantities relate across different classes of structural variants.

Structural variant cellular fraction (SVCF) quantifies the fraction of cells in a sample containing an SV. It can be expressed as a product of two observable variables from bulk DNA sequencing data: the average number of break intervals ($\overline{R}$) and variant allele frequency (VAF),

$$SVCF = \overline{R} \times VAF \quad (S1)$$

Here, SV breakpoints are genomic sites of somatic alterations producing juxtaposition of distant sequences in the normal genome, and they result in sequence reads absent from the normal genome (SV supporting reads). SV breakends are genomic sites of somatic alterations that mark the start and end of an SV, and they result in sequence reads that are also present in the normal genome (reference reads). Break intervals are the regions between pairs of breakpoints, breakends, or a combination of both (main-text Fig. 1A-F). In the following derivations, variables with uppercase letters refer to values for the total number of cells in a sample, and variables with lowercase letters refer to values for a single cell. SVCF is the number of cells containing a structural variant ($SVC$) divided by the total number of cells ($N$) sampled from bulk sequencing of a tumor.

$$SVCF = \frac{SVC}{N} \quad (S2)$$

The number of breakpoints in a structural variant depends on the types of structural variants. Deletions in a single cell have one breakpoint (main-text Fig. 1A-F). Therefore, in a sample of $N$ cells, the total count of breakpoints BPC for each deletion is the same as the number of cells that harbor that deletion. Therefore,

$$SVCF_{del} = \frac{BPC}{N \times 1} \quad (S3)$$

Tandem duplications may have more than one breakpoint in each cell, and the number of breakpoints depends on the number of tandem copies (and resulting break intervals).

When the number of break intervals $r$ in each cell is $r = 3$, a tandem duplication has 1 breakpoint, and when $r = 4$, it has 2 breakpoints (main-text Fig. 1A-F). In general, tandem duplications have $r - 2$ breakpoints in a single cell. Therefore, for tandem duplications,

$$SVCF_{dup} = \frac{BPC}{N \times (r-2)} \quad (S4)$$

Because inversions always have two breakpoints (main-text Fig. 1A-F), for inversions,

$$SVCF_{inv} = \frac{BPC}{N \times 2} \quad (S5)$$

There are three types of translocations, including copy-paste, cut-paste, and reciprocal. Each type has a different number of breakpoints (main-text Fig. 1A-F), and for translocations:

$$SVCF_{cut-trans} = \frac{BPC}{N \times 3} \quad (S6)$$

$$SVCF_{copy-trans} = \frac{BPC}{N \times 2} \quad (S7)$$

$$SVCF_{reci-trans} = \frac{BPC}{N \times 4} \quad (S8)$$

The definition of variant allele frequency (VAF) is the ratio of reads supporting an alternative allele and the sum of reference and alternative reads. For SVs, the equation is

$$VAF = \frac{BPRC}{BPRC + BERC} \quad (S9)$$

where $BPRC$ (breakpoint read count) is the total number of reads at breakpoints in $N$ cells, and $BERC$ (breakend read count) is the total number of reads at breakends in $N$ cells. Both $BERC$ and $BPRC$ can be expressed in terms of the breakend count ($BEC$) or breakpoint count ($BPC$), along with the averaged read depth ($\overline{RD}$) at the corresponding breakend or breakpoint genomic location across all cells in the sample. Therefore, Eq S9 can be rewritten as,

$$VAF = \frac{BPC \times \overline{RD}}{(BPC + BEC) \times \overline{\overline{RD}}} \quad (S10)$$

Given:

$$\overline{RD} = \frac{\sum_{i=1}^{N} r\, d_i}{N} \quad (S11)$$

For all SV types, $\overline{RD}$ can be separated into the average read depth at a breakpoint $\overline{RD}_{bp}$ and average read depth at left and right breakends $\overline{RD}_{bel}$, $\overline{RD}_{ber}$ (main-text Fig. 1A-F),

$$VAF = \frac{BPC \times \overline{RD}_{bp}}{BEC_l \times \overline{RD}_{bel} + BEC_r \times \overline{RD}_{ber} + BPC \times \overline{RD}_{bp}} \quad (S12)$$

$$= \frac{BPC}{BEC_l \times \frac{\overline{RD}_{bel}}{\overline{RD}_{bp}} + BEC_r \times \frac{\overline{RD}_{ber}}{\overline{RD}_{bp}} + BPC} \quad (S13)$$

Because

$$\frac{\overline{RD}_{bel}}{\overline{RD}_{bp}} \approx \frac{\overline{RD}_{ber}}{\overline{RD}_{bp}} \approx 1 \quad \text{(Supplementary Note S6)} \quad (S14)$$

Therefore,

$$VAF = \frac{BPC}{BEC_l + BEC_r + BPC} \quad (S15)$$

$$= \frac{BPC}{BEC + BPC} \quad (S16)$$

Eq S9 is adjusted for deletions and tandem duplications, as these variants result in a doubling of breakend read counts, which flank the break interval (main-text Fig. 1A-F). These additional reads do not represent the true SVCF, so to obtain the correct VAF, the $BERC$ must be divided by two.

$$VAF_{del} = VAF_{dup} = \frac{BPRC}{BPRC + (0.5 \times BERC)} \quad (S17)$$

Because $BPRC = BPC \times \overline{RD}$ and $BERC = BEC \times \overline{RD}$ and Eq S14,

$$VAF_{del} = VAF_{dup} = \frac{BPC}{BPC + (0.5 \times BEC)} \quad (S18)$$

For inversions, cut-paste translocations, and reciprocal translocations, breakend read counts and breakpoint read counts are at one-to-one ratio (main-text Fig. 1A-F) therefore,

$$VAF_{inv} = VAF_{cut-trans} = VAF_{reci-trans} = \frac{BPC}{BPC + BEC} \quad (S19)$$

Because the original genome segment is preserved in copy-paste translocations, they are treated as if the SV is overlapped with a duplication event (Supplementary Note S5). Here, we treat copy-paste translocations similarly to other translocations,

$$VAF_{copy-trans} = \frac{BPC}{BPC + BEC} \quad (S20)$$

In practice, directly calculating Eq S3, S4 and S5 is not possible, because they depend on unobservable variables $N$, $BEC$ and $BPC$. However, by the multiplicative identity property, for deletions and tandem duplications (Supplementary Note S7):

$$\frac{R}{\frac{BEC}{2}+BPC}=1 \quad (S21)$$

for inversions:

$$\frac{R}{\frac{BEC}{2}+\frac{BPC}{2}}=1 \quad (S22)$$

for cut-paste translocations:

$$\frac{R}{\frac{BEC}{3}+\frac{BPC}{3}}=1 \quad (S23)$$

for copy-paste translocations:

$$\frac{R}{\frac{BEC}{2}+\frac{BPC}{2}}=1 \quad (S24)$$

and for reciprocal translocations:

$$\frac{0.5R}{\frac{BEC}{4}+\frac{BPC}{4}}=1 \quad (S25)$$

These multiplicative identity properties can be used to estimate SVCF for inversions, tandem duplications and deletions from observable variables VAF, $\overline{R}$, and $r$.

$$SVCF_{del}=1\times\frac{BPC}{N} \quad (S26)$$

$$=\frac{R}{0.5\times BEC+BPC}\times\frac{BPC}{N} \quad (S27)$$

$$=\frac{R}{N}\times\frac{BPC}{0.5\times BEC+BPC} \quad (S28)$$

$$=\overline{R}\times VAF \quad (S29)$$

$$SVCF_{dup}=1\times\frac{BPC}{N\times(r-2)} \quad (S30)$$

$$=\frac{R}{0.5\times BEC+BPC}\times\frac{BPC}{N\times(r-2)} \quad (S31)$$

$$= \frac{R}{N \times (r-2)} \times \frac{BPC}{0.5 \times BEC + BPC} \quad (S32)$$

$$= \frac{\overline{R} \times VAF}{r-2} \quad (S33)$$

$$SVCF_{inv} = 1 \times \frac{BPC}{N \times 2} \quad (S34)$$

$$= \frac{R}{0.5 \times BEC + 0.5 \times BPC} \times \frac{BPC}{N \times 2} \quad (S35)$$

$$= \frac{R}{N} \times \frac{BPC}{2 \times (0.5 \times BEC + 0.5 \times BPC)} \quad (S36)$$

$$= \frac{R}{N} \times \frac{BPC}{BEC + BPC} \quad (S37)$$

$$= \overline{R} \times VAF \quad (S38)$$

$$SVCF_{cut-trans} = 1 \times \frac{BPC}{N \times 3} \quad (S39)$$

$$= \frac{R}{\frac{BEC}{3} + \frac{BPC}{3}} \times \frac{BPC}{N \times 3} \quad (S40)$$

$$= \frac{R}{N} \times \frac{BPC}{3 \times (\frac{BEC}{3} + \frac{BPC}{3})} \quad (S41)$$

$$= \frac{R}{N} \times \frac{BPC}{BEC + BPC} \quad (S42)$$

$$= \overline{R} \times VAF \quad (S43)$$

$$SVCF_{copy-trans} = 1 \times \frac{BPC}{N \times 2} \quad (S44)$$

$$= \frac{R}{\frac{BEC}{2} + \frac{BPC}{2}} \times \frac{BPC}{N \times 2} \quad (S45)$$

$$= \frac{R}{N} \times \frac{BPC}{2 \times (\frac{BEC}{2} + \frac{BPC}{2})} \quad (S46)$$

$$= \frac{R}{N} \times \frac{BPC}{BEC + BPC} \quad (S47)$$

$$= \overline{R} \times VAF \quad (S48)$$

$$SVCF_{reci-trans} = 1 \times \frac{BPC}{N \times 4} \quad (S49)$$

$$= \frac{0.5R}{\frac{BEC}{4} + \frac{BPC}{4}} \times \frac{BPC}{N \times 4} \quad (S50)$$

$$= \frac{0.5R}{N} \times \frac{BPC}{4 \times (\frac{BEC}{4} + \frac{BPC}{4})} \quad (S51)$$

$$= \frac{0.5R}{N} \times \frac{BPC}{BEC + BPC} \quad (S52)$$

$$= \overline{R} \times VAF \quad (S53)$$

In the absence of an overlapping duplication, inversions, cut-paste translocations, and reciprocal translocations are assumed to be diploid and $\overline{R} = 2$ (main-text Fig. 1A-F) and for deletions $\overline{R} = 2$ (Supplementary Note S7). FACETS[2] total copy number output (the sum of the major and minor copy number) is used as $r$ for tandem duplication. Copy-paste translocations are always in non-diploid region and additional modification to Eq S20 is required as described in Supplementary note S2. After additional modification, $\overline{R} = 2$ will also apply to copy-paste translocation. In scenarios where deletion overlapped with deletions or duplication overlapped with duplications, we used the uncorrected output to avoid over-correcting.

#### Supplementary Note S2: Algorithm convergence and SV-CNV ordering

> **Cross-reference to main text.** The closed-form fixed points proved here appear as Eqs. 5 and 6 of the main text. The two iterative update rules whose convergence is proved in Sections 2.1 and 2.2 (with auxiliary variables $\alpha_q$ and $\beta_q$) appear as Eqs. 8 and 9 of the main-text Methods ("Iterative SVCF estimation in non-diploid regions" subsection). The sign criterion proved in Section 2.3 is referenced in the closing paragraph of the "Estimation in non-diploid genomic regions" Results subsection (with a coverage caveat) and illustrated by Supplementary Fig. S2 below.

SVCFit estimates SVCF in regions affected by overlapping SV and CNV events using an iterative optimization procedure. This note demonstrates that the algorithm converges to stable closed-form solutions and shows how the inferred solutions encode the relative ordering of SV and CNV events. These convergence results concern diploid loci, where each update rule is iterated to its closed-form fixed point. On a hemizygous chromosome the corresponding estimators (Methods, "Estimation on hemizygous chromosomes") are exact closed forms that require no iteration, and so fall outside the scope of this note.

##### 2.1: Convergence for CNV occurs after an in cis SV

We show that the iterative update rule used by SVCFit converges to a unique, stable fixed point when a copy-number alteration (CNV) occurs after an in cis structural variant (SV). Here, we used inversion as an example (Eq S37).

Proof.

Let $SVCF_q \in (0,1]$ denote the estimated SV cellular fraction at iteration $q$, initialized arbitrarily (we use $SVCF_0 = 0.5$). The update rule is defined as:

$$SVCF_{q+1} = \frac{\overline{R} \times (BPC \times \alpha_q)}{BPC \times \alpha_q + BEC}, \quad \text{where} \quad \alpha_q = \frac{SVCF_q}{SVCF_q + ACR - 1} \quad \text{and} \quad R = 2$$

Substituting $\alpha_q$ and simplifying yields the recurrence:

$$SVCF_{q+1} = \frac{2 \times BPC \times SVCF_q}{(BPC + BEC)SVCF_q + BEC \times (ACR - 1)}$$

Assume the sequence converges to a limit $L \in (0,1]$. Taking the limit as $q \to \infty$ and solving for the fixed point gives:

$$L = \frac{2 \times BPC - BEC \times (ACR - 1)}{BPC + BEC}$$

This fixed point corresponds to the closed-form SVCF solution for the CNV occurs after an in cis SV case (main-text Eq. 5). Therefore, the iterative update converges to a stable solution consistent with the analytic formulation. ■

#### 2.2: Convergence for CNV precedes an in cis SV or is in trans with an SV

We next consider the case in which a copy-number alteration (CNV) occurs before an in cis structural variant (SV), or affects a different haplotype (in trans). Under these conditions, SVCFit applies an alternative iterative update rule. We show that this update also converges to a stable closed-form solution. Here, we used inversion as an example (Eq S37).

Proof.

Let $SVCF_q \in (0,1]$ denote the SV cellular fraction estimate at iteration $q$, initialized arbitrarily. The update rule is:

$$SVCF_{q+1} = \frac{\overline{R} \times BPC}{BPC + BEC \times \beta_q}, \quad \text{where} \quad \beta_q = \frac{2 - SVCF_q}{1 + ACR - SVCF_q}, \quad \text{and} \quad R = 2$$

Substituting $\beta_q$ into the update rule and simplifying yields:

$$SVCF_{q+1} = \frac{2 \times BPC \times (1 + ACR) - 2 \times BPC \times SVCF_q}{BPC + BPC \times ACR + 2BEC - (BPC + BEC) \times SVCF_q}$$

Assuming convergence to a limit $L \in (0,1]$, taking the limit as $q \to \infty$ yields a quadratic equation with two fixed points. One solution corresponds to a constant value outside the biologically meaningful range and is discarded. The remaining solution is:

$$L = \frac{BPC + BPC \times ACR}{BPC + BEC}$$

This expression corresponds to the closed-form SVCF estimate for CNV precedes an in cis SV or is in trans with an SV (main-text Eq. 6, Eq. S69). Thus, the iterative procedure converges to a stable and biologically valid solution in this case. ■

#### 2.3: Inferring relative SV-CNV ordering

The closed-form solutions derived above encode information about the relative ordering of overlapping structural variants (SVs) and copy-number alterations (CNVs). Here, we show that the SVCF solution for the SV occurring before CNV case is non-positive when the SV occurs after the CNV, enabling inference of relative event ordering. This result is the formal basis for the SV-CNV ordering criterion stated in the closing paragraph of the main-text Results "Estimation in non-diploid genomic regions" subsection (with a coverage caveat) and illustrated in Supplementary Fig. S2 below.

Proof.

Consider a tumor sample containing $C$ total cells, of which $D$ cells harbor a CNV that increases copy number by $k \geq 1$, and $N$ cells remain diploid. The total allele count in the sample is:

$$A_{tot} = D(2 + k) + 2N$$

Suppose an SV affects $m$ alleles in the sample. Because the SV occurs after the CNV, it can only affect alleles present in CNV-bearing cells, implying $m \leq D$. The breakpoint and breakend read counts are therefore:

$$BPC = 2m$$

$$BEC = 2(A_{tot} - m) = 2(D(2 + k) + 2N - m)$$

The allele-specific copy number is:

$$ACR = \frac{kD}{C} + 1$$

Substituting these expressions into the fixed-point solution for the SV occurring before CNV case (main-text Eq. 5) yields:

$$L = \frac{2 \times BPC - BEC \times (ACR - 1)}{BPC + BEC}$$

Because all denominator terms are strictly positive, the sign of $L$ is determined by the numerator. Under the constraint $m \leq D$, the numerator is non-positive, implying $L \leq 0$. Therefore, a non-positive solution indicates that the SV occurred after the CNV. Conversely, when the SV occurs before the CNV, the corresponding solution remains positive. Thus, the sign of the inferred SVCF solution provides a principled criterion for inferring the relative ordering of in cis SV and CNV events. ■

### Supplementary Note S3: Longitudinal self-evaluation on paired pre/post-treatment simulations

**Cross-reference to main text.** This Note backs the Results subsection "Longitudinal self-evaluation on paired pre/post-treatment simulations" and the Methods subsection "Longitudinal self-evaluation simulations" under "SVCF comparison and benchmarking", together with Supplementary Figs. S8, S9, S10, S11. The simulation design and pipeline summary are reproduced below for self-contained reference; the unique content of this Note is the per-purity numerical tables (S3-1, S3-2), the multi-covariate analysis of per-clone signed CCF residuals (S3.5), and the operating-range conclusions (S3.6).

#### S3.1 Simulation design

Paired pre- and post-treatment whole-genome simulations were generated with VISOR[1] following a single scenario (S1): a truncal clone with CCF = 1.0 at both timepoints plus two subclonal children that diverge under simulated therapeutic selection; one declines (pre-treatment CCF = 0.83 → post = 0.17) and one expands (pre = 0.17 → post = 0.83). Each paired pre/post replicate carried 33 truncal SVs and 33 and 34 subclonal SVs on the two terminal subclones, drawn from a mixture of deletions, inversions, tandem duplications, and translocations. SV tumor purity was swept over {20%, 40%, 60%, 80%}, with the five SV-CNV overlap configurations and bootstraps 0-4 yielding 25 paired pre/post replicates per purity, except at 40% purity, where one replicate failed to build a tree (n = 24). Sequencing reads were simulated at 50× mean genome-wide coverage with a matched normal at 50× per replicate.

#### S3.2 SV calling, CCF estimation, and tree reconstruction

Reads were aligned to GRCh38 with BWA-MEM[23] (v0.7.19) and SVs were called with Manta[24] (v1.6.0) followed by SVtyper[25] (v0.7.1) for breakpoint-resolved genotyping. Allele-specific copy number was estimated with FACETS (v0.6.2). CCFs were estimated for each SV using SVCFit at default parameters (no per-scenario tuning; DP-GMM concentration $\alpha$ = 1, kmax = 10, covariance_type = "full", init_params = "k-means++"). SVs were clustered jointly in the two-dimensional (pre-treatment CCF, $\Delta$CCF = post-treatment CCF − pre-treatment CCF) space using a Dirichlet-process Gaussian mixture model. Post-clustering, components containing fewer than five SVs were merged with their nearest neighbor by Euclidean distance; the five-SV floor reflects the minimum number of observations required for stable 2D covariance estimation ($N = D + 1 = 3$ non-collinear points is the mathematical minimum; 5 provides a conservative practical floor against both numerical instability and false-positive artifact clusters, consistent with standard practice in published tumor subclonal-reconstruction pipelines[26,27]; see Methods "SV Longitudinal Clustering and Dynamics" for the full three-part justification). Pre- and post-treatment clusters were matched into clones, and the rooted clone phylogeny was reconstructed using SVCFit's modified Gabow-Myers tree builder.

#### S3.3 Evaluation metrics

For each replicate we computed:

1. **Per-SV truncal vs. subclonal classification.** Each SV was labeled truncal (true CCF = 1.0 at both timepoints) or subclonal (otherwise) in ground truth and as inferred. We report **balanced accuracy** and **F1** (truncal = positive class), evaluated separately at pre- and post-treatment.
2. **Per-clone CCF estimation.** For each true clone (Trunk, Subclone 1, Subclone 2), we computed the **signed error** ($CCF_{true} - CCF_{est}$) between the true CCF and the inferred centroid of the matched cluster, at each timepoint. Signed error is reported in preference to absolute error because the direction of error is itself diagnostic.
3. **Tree topology recovery.** A replicate scored 1 if the inferred rooted topology was an exact match to the S1 ground-truth tree (root → trunk → two children) and 0 otherwise.

Per-purity summaries report means, with bootstrap 95% confidence intervals for balanced accuracy, CCF error and topology recovery. Balanced accuracy, F1 and topology recovery use all 24-25 replicates per purity; CCF error uses the 20 to 22 replicates per purity that yielded clone CCFs.

#### S3.4 Operating-range results

**Truncal vs. subclonal classification was robust across the 20-80% purity range** (Supplementary Fig. S8A). Balanced accuracy increased modestly with purity, from 0.79 at 20% purity (pre 0.784 [95% CI 0.749-0.820]; post 0.797 [0.756-0.837]) to 0.89 at 80% purity (pre 0.882 [0.867-0.897]; post 0.896 [0.882-0.910]), and was nearly identical at the pre- and post-treatment timepoints. Per-purity classification, CCF MAE, and topology results are summarized in Table S4. Even at the lowest purity tested, the classifier recovered the truncal/subclonal partition with balanced accuracy ≥ 0.78, indicating that the relative ordering of CCFs, rather than absolute values, drives this assay and is preserved even when absolute CCF resolution is degraded.

**Per-clone CCF point estimates show a systematic positive signed error** (Supplementary Fig. S8B). Across replicates the truncal cluster is underestimated by 15-26% across the purity range (inferred centroid 0.74-0.85 against true 1.00), and per-clone mean absolute error decreases modestly with purity (0.18 at 20% to 0.12 at 80%). The pattern is consistent across clones: high-CCF subclones (true CCF ~0.83) carry comparable underestimation to the trunk, while low-CCF subclones (true CCF ~0.17) are estimated with near zero error at 40% purity and above. At 20% purity these low-CCF clusters collapse toward zero (true 0.150 and 0.200 inferred at 0.027 and 0.034). Per-clone inferred-vs-true CCFs at each purity are summarized in Table S3.

**Table S3. Per-clone inferred-vs-true CCFs at each purity, computed over the replicates that yielded clone CCFs (22 at 20% purity, 19 to 20 at 40%, 20 to 21 at 60%, 22 at 80%, against 25, 24, 25 and 25 replicates run at each purity).** Subclones are labeled Sub1 (declining clone) and Sub2 (expanding clone). The trunk and the high-CCF clone at each timepoint carry comparable underestimation; low-CCF subclones (true CCF ~0.17) are accurately estimated at 40% purity and above, but collapse toward zero at 20% purity.

| Purity | Clone | True CCF (pre) | Inferred CCF (pre, mean ± SD) | True CCF (post) | Inferred CCF (post, mean ± SD) |
|---|---|---|---|---|---|
| 20% | Trunk | 1.000 | 0.795 ± 0.054 | 1.000 | 0.803 ± 0.050 |
| 20% | Sub1 | 0.800 | 0.634 ± 0.026 | 0.150 | 0.027 ± 0.026 |
| 20% | Sub2 | 0.200 | 0.034 ± 0.039 | 0.850 | 0.654 ± 0.041 |
| 40% | Trunk | 1.000 | 0.778 ± 0.136 | 1.000 | 0.741 ± 0.149 |
| 40% | Sub1 | 0.825 | 0.645 ± 0.097 | 0.150 | 0.117 ± 0.156 |
| 40% | Sub2 | 0.175 | 0.143 ± 0.060 | 0.850 | 0.674 ± 0.067 |
| 60% | Trunk | 1.000 | 0.778 ± 0.210 | 1.000 | 0.769 ± 0.203 |
| 60% | Sub1 | 0.833 | 0.671 ± 0.107 | 0.167 | 0.190 ± 0.100 |
| 60% | Sub2 | 0.167 | 0.182 ± 0.100 | 0.833 | 0.687 ± 0.105 |
| 80% | Trunk | 1.000 | 0.846 ± 0.050 | 1.000 | 0.828 ± 0.054 |
| 80% | Sub1 | 0.825 | 0.649 ± 0.091 | 0.163 | 0.163 ± 0.041 |
| 80% | Sub2 | 0.175 | 0.185 ± 0.046 | 0.838 | 0.696 ± 0.131 |

**Table S4.** Per-purity summary of the three downstream metrics (per-SV balanced accuracy and F1, per-clone CCF mean absolute error, rooted-topology recovery), split by timepoint. F1 uses truncal as the positive class. Balanced accuracy, CCF MAE and topology recovery carry bootstrap 95% confidence intervals; F1 is a mean without an interval. Topology recovery is a per-purity quantity and is not split by timepoint. Balanced accuracy, F1 and topology use all replicates (25, 24, 25 and 25 at 20%, 40%, 60% and 80% purity); CCF MAE is computed per replicate as the mean over the three clones and uses those replicates that yielded clone CCFs (22, 20, 21 and 22).

| Purity | Timepoint | Balanced accuracy [95% CI] | F1 | CCF MAE [95% CI] | Topology recovery (n/N; [95% CI]) |
|---|---|---|---|---|---|
| 20% | Pre | 0.784 [0.749-0.820] | 0.772 | 0.179 [0.166-0.193] | 22/25 = 88.0% [70.0%-95.8%] |

| Purity | Timepoint | Balanced accuracy [95% CI] | F1 | CCF MAE [95% CI] | Topology recovery (n/N; [95% CI]) |
|---|---|---|---|---|---|
| | Post | 0.797 [0.756-0.837] | 0.771 | 0.172 [0.158-0.185] | |
| 40% | Pre | 0.798 [0.734-0.862] | 0.769 | 0.151 [0.128-0.173] | 20/24 = 83.3% [64.1%-93.3%] |
| | Post | 0.824 [0.769-0.878] | 0.789 | 0.183 [0.151-0.216] | |
| 60% | Pre | 0.784 [0.708-0.860] | 0.755 | 0.143 [0.105-0.181] | 22/25 = 88.0% [70.0%-95.8%] |
| | Post | 0.792 [0.709-0.875] | 0.769 | 0.146 [0.106-0.186] | |
| 80% | Pre | 0.882 [0.867-0.897] | 0.849 | 0.121 [0.109-0.133] | 22/25 = 88.0% [70.0%-95.8%] |
| | Post | 0.896 [0.882-0.910] | 0.865 | 0.117 [0.093-0.140] | |

**Rooted tree topology was correctly recovered in 83-88% of replicates across the purity range** (Supplementary Fig. S8C), with no monotonic dependence on purity (22/25 = 88% at 20%, 20/24 = 83% at 40%, 22/25 = 88% at 60%, 22/25 = 88% at 80%; Table S4). The mild improvement with purity reflects the fact that the binary trunk-vs-subclone partition that determines topology is preserved in most replicates, even though absolute CCF estimates degrade at low purity. An illustrative successfully recovered topology at 60% purity is shown in Supplementary Fig. S9: the inferred rooted topology (panel B) matches ground truth (panel A) with the trunk and two diverging subclones correctly recovered; inferred clone-CCF centroids fall close to their true coordinates in paired (pre, post) space (panel C); and the per-SV confusion matrices at both timepoints show the truncal/subclonal assignment is dominantly correct (panel D). Even in this "what success looks like" replicate, the trunk centroid is inferred at 0.81/0.86 against true 1.00 and the post-treatment dominant subclone at 0.65 against true 0.83, the same direction of bias visible in Supplementary Fig. S8B.

#### S3.5 Mechanism of the per-clone CCF residual

Trunk underestimation at every purity level reflects two properties of the bounded estimator and the clustering step rather than a SVCFit failure mode:

1. **CCF = 1.0 ceiling effect.** The truncal CCF is bounded above at 1.0, so any inferential noise can only push the inferred centroid below truth, producing a one-sided positive signed bias even under unbiased estimation. This effect is amplified at low purity, where the centroid noise is largest.

2. **Sampling noise dominates the aggregate cluster-size effect, with a residual BND-specific contribution.** At the aggregate (un-stratified) level, per-clone signed CCF error decreases as the per-pair number of SVs assigned to the cluster grows (Sub1 ρ = −0.51 [−0.68, −0.30]; Sub2 ρ = −0.40 [−0.58, −0.18]; trunk ρ = −0.01 [−0.25, +0.22], no monotone relationship; Supplementary Fig. S10), consistent with the expected sampling-noise effect that more SVs per cluster support a tighter centroid estimate.

3. The truncal panel of Fig. S10 carries a higher linear-fit $R^2$ (0.274) than the rank correlation alone would suggest because a single high-leverage 60% purity bootstrap replicate, in which the DP-GMM split the truncal SVs into multiple components, produced a small truncal cluster with an unusually large CCF residual; Spearman ρ remains the robust summary statistic for the per-cluster size effect.

4. SV-type stratification (Supplementary Fig. S11) reveals a BND-specific contribution that the aggregate signal masks: within breakend-type SVs, cluster size and signed CCF error correlate **positively** in both subclones (ρ ≈ +0.49 in Subclone 1, +0.56 in Subclone 2), mean SV segment length correlates **negatively** (ρ ≈ −0.51 and −0.52, respectively), and mean SVtyper AO is negative for BND across all clones (trunk ρ ≈ −0.33). Short BNDs carry lower mappability and noisier alternate-allele counts; a cluster dominated by many short BNDs carries noisier per-SV AO and a larger cluster-level CCF residual, even when the aggregate cluster-size signal is dominated by sampling-noise gains in non-BND SV types. The per-clone signed CCF error vs. number of SVs assigned to the cluster is shown as Supplementary Fig. S10; the SV-type stratification of cluster-size correlations is shown as Supplementary Fig. S11.

To confirm that these mechanisms, rather than read-support depth, drive the residual, we tested per-SV alternate-allele read support (mean SVtyper AO), cluster size (number of SVs assigned to the cluster), mean SV segment length, and ACR mismatch fraction as covariates of per-clone signed CCF error (Spearman ρ with bootstrap 95% CI; mixed-effects regression `lmer(signed_error ~ log10(covariate) × clone + (1 | replicate))` with likelihood-ratio test on the interaction). Results were stratified by SV type (DEL, DUP, INV, INS, TRA/BND) to localize any class-specific contribution.

The dominant cluster-level signal is **SV-type composition**, not read-support depth. Per-SV AO, if the primary driver, should show the strongest correlation with subclone residuals; instead it explains little of the subclone residual (Sub1 ρ = −0.20 [−0.34, −0.03]; Sub2 ρ = −0.15 [−0.31, +0.03], CI includes zero) and carries its strongest correlation with the trunk (ρ = −0.35 [95% CI −0.48, −0.19]), opposite to the prediction of a coverage-limited subclone regime.

To test whether a per-SV AO undercount contributes to the trunk bias, we computed mean `raw_svcf / p` across non-DUP trunk SVs in diploid background segments, where the formula reduces to $raw\_svcf = 2 \times AO/(AO + RO)$ and the theoretical expectation is $raw\_svcf/p = 1$. The empirical ratio had a median near the theoretical expectation of 1.0 at every purity (median 1.019 at 20%, 0.957 at 40%, 0.995 at 60%, 0.993 at 80%; n = 270-281 SVs per purity).

The pooled mean is inflated at low purity (1.198 at 20%, 1.118 at 40%, converging to 0.984 at 60% and 0.949 at 80%) by a heavy right tail of SVs with above-expected AO, a variance effect from Manta selection bias and stochastic noise at low read counts rather than a systematic bias. There is thus no meaningful systematic per-SV AO undercount; any residual shortfall at high purity is small (at most 5.1%), consistent with the modest sensitivity limits of breakpoint-resolved SV genotyping reported in independent benchmarks[28], but far too small to account for the 15 to 26% cluster-level trunk CCF underestimate, which is instead a property of the DP-GMM clustering step.

The non-monotonic purity pattern (largest bias at 40-60%, smallest at 80%) is therefore a product of DP-GMM cluster instability at low purity interacting with the ceiling effect, rather than a consequence of FACETS copy-number-estimation variability; the simulation pipeline uses ground-truth purity throughout (passed via `pur_path` in the longitudinal driver), so FACETS' purity estimate is bypassed.

#### S3.6 Operating-range summary

Taken together, these results define a clear operating range for SVCFit's downstream phylogenetic inferences. At ≥40% tumor purity, SVCFit recovers the rooted topology in ≥80% of replicates and estimates subclonal CCFs with diminishing residual error as purity increases. The systematic underestimation of truncal CCF is a characterizable property of the bounded estimator and the clustering step rather than a failure mode, and is straightforwardly accounted for in downstream interpretation. 20% purity remains a soft lower bound: classification and topology are preserved in most replicates, but per-clone CCF estimates carry substantial residual error. We document this as the method's data-driven operating-range boundary rather than a tunable parameter choice.

#### Supplementary Note S4: Cohort-wide SV-based vs SNV-based clonal phylogenies in COMBAT

**Cross-reference to main text.** Subjects 10 and 31 are detailed in the main-text Results "Longitudinal analysis of SV-defined clones in mCRPC subjects receiving bipolar androgen therapy" subsection (Fig. 5). This Note presents the corresponding SV- and SNV-based phylogeny analysis for the 9 additional subjects with paired pre/on-BAT WGS suitable for analysis, aggregates the SV-vs-SNV comparison across all 11 analyzed subjects, and shows the paired SV and SNV phylogeny diagrams for all 11 subjects (S4.4).

##### S4.1 Methods overview

SV-based phylogenies were reconstructed using SVCFit as described in Methods. SNV-based phylogenies were reconstructed in parallel from somatic single-nucleotide variants called from the same WGS data using the released, peer-reviewed PICTograph package[6], with prostate-cancer driver annotations applied to clustered SNVs. PICTograph and SVCFit share the same modified Gabow-Myers tree-construction and tree-scoring approach, so the SV- and SNV-based phylogenies differ only in the input variant class and its clustering. Per-subject summaries below tabulate, for each tree type, topology, trunk annotations, sensitive- and resistant-clone annotations, clonal dynamics across the pre-/on-BAT pair, and a short clinical-insight verdict comparing the two tree types. SNV driver mutations are given as the gene symbol with the amino-acid change (for example, *SPOP* Y83C). Clones and clusters are numbered in the order they appear on the tree: the trunk is 1, the declining clone is 2 and is drawn on the left, and the emerging clone is 3 and is drawn on the right. Roles are assigned from the inferred cluster cellular fractions rather than from the raw cluster index, which PICTograph and the SVCFit clustering both assign arbitrarily per run. Pair labels are paired-biopsy identifiers; Subject numbers are the trial identifiers used elsewhere in this manuscript. Featured Subjects 10 and 31 are not retabulated here; their detailed analysis appears in the main text and Fig. 5.

The cohort comprises 11 of the 12 paired-biopsy subjects on the COMBAT trial. **Pair 9 was excluded from the SVCFit analysis** because FACETS could not estimate tumor purity from the on-BAT biopsy (returning the diagnostic "Not enough information to estimate purity"), most likely owing to too few informative germline heterozygous SNPs in the matched normal/tumor pileup. Without a reliable purity estimate, the SVCF → CCF conversion (Eq. 11) and the haplotype-aware ACR inference required for non-diploid SVCF estimation cannot be applied, so we excluded Pair 9 from both the SV-tree and the side-by-side SV-vs-SNV cohort analysis rather than report results from a partially specified pipeline.

One note on interpretation. For the three subjects whose SNV tree is a linear chain (Subjects 7, 20, 49; root → cluster 1 → cluster 2), cluster 2 is a descendant of cluster 1, so cluster 1's mutations are inherited by every cluster-2 cell and remain present throughout treatment; what changes is the fraction of cells carrying cluster 2's additional private mutations. We therefore describe these as genotype shifts within a single lineage rather than as replacement of one clone by another.

#### S4.2 Per-subject summaries (9 non-featured subjects)

**Table S5. Per-subject SV-tree vs SNV-tree case summaries across all 11 COMBAT subjects (Subjects 10 and 31 are also featured in the main text).** Each subject is presented as a side-by-side comparison of the SV-based and SNV-based phylogenies, including topology, trunk and branch annotations, clonal dynamics across the pre-/on-BAT pair, and a clinical-insight comparison of the two tree types. The SNV-tree columns reflect the released PICTograph reconstruction.

##### *Subject 1 (Pair 1): Acquired Resistance*

| | SV tree | SNV tree |
|---|---|---|
| **Topology** | Linear: G → Clone 1 (resistant) → Clone 2 (sensitive) (2 clones; no separate trunk clone) | Linear: G → Clone 1 (resistant) → Clone 2 (sensitive); 2 clones (no separate trunk clone) |
| **Trunk annotations** | Not applicable, linear topology | None |
| **Sensitive clone** | Clone 2: TP53 inversion (intron1-intron4), SETD2 del | No driver-gene SNVs |
| **Resistant clone** | Clone 1: TMPRSS2-ERG fusion (IHC-confirmed ERG expression at both timepoints), SETD2 deletion, and BRAF duplication (recognized as a driver alteration in prostate cancer7,8); dominates on-BAT | Not resolved as a separate clone |
| **Clonal dynamics** | Clone 2 (Sensitive) eliminated; Clone 1 (Resistant) becomes dominant on-BAT | Genotype shift within a single lineage: the clone-1 private genotype expands to dominance on-BAT (ancestral clone-2 genotype retained throughout) |
| **Clinical insight** | The resistant clone (Clone 1) carrying BRAF duplication7,8 pre-exists therapy and expands markedly on-BAT. *BRAF duplication activates the* | Resolves two clones and the clonal shift but carries no driver-gene SNVs, so it cannot attribute the shift to specific driver annotations |

| | SV tree | SNV tree |
|---|---|---|
| | *RAS/RAF/MAPK pathway*, providing AR-independent proliferative signaling that can sustain growth across the testosterone cycling imposed by BAT. The sensitive clone (Clone 2) carrying *TP53* inv (intron1-intron4) and a second SETD2 deletion event (biallelic with the truncal hit) is eliminated; the functional consequences of the specific intron1-intron4 *TP53* inversion and biallelic SETD2 deletion for BAT sensitivity in this clone are of uncertain mechanistic significance beyond their known roles in epigenetic dysregulation and impaired chromatin-coupled DNA repair. The linear topology places resistance on a clone that pre-exists therapy rather than on a separately acquired branch | |

PSA dropped sharply from ~28 to ~2 ng/dL over 56 days, then rose slightly at day 84; Ki-67 ~30 → ~40%; tumor volume stable.

##### *Subject 6 (Pair 2): Stable Disease*

| | SV tree | SNV tree |
|---|---|---|
| **Topology** | Branched: trunk (Clone 1) → Clone 2 (sensitive), Clone 3 (resistant) | Branched: trunk (Clone 1) → Clone 2 (sensitive), Clone 3 (resistant); 3 clones |
| **Trunk annotations** | *ATM del, PTEN del (intron5-exon9)*, *TMPRSS2-ERG* fusion (IHC-confirmed ERG expression at both timepoints) | *KMT2C* E534fs (frameshift), truncal |

| | SV tree | SNV tree |
|---|---|---|
| **Sensitive clone** | Clone 2: ARID1A del, ARID2 inversion | No branch-specific driver SNVs |
| **Resistant clone** | Clone 3: NKX3-1 del, AURKA dup | No branch-specific driver SNVs |
| **Clonal dynamics** | Near-complete swap: Clone 2 eliminated, Clone 3 expands | Dominant child (pre-BAT) nearly eliminated; second child expands from absent to dominant on-BAT |
| **Clinical insight** | *AURKA duplication on the resistant clone is a candidate* actionable and mechanistically rationalized target. Aurora Kinase A reconfigures AR pre-mRNA splicing to upregulate AR-V7 transcripts9 and phosphorylates SPOP, which normally ubiquitinates and degrades AR, causing SPOP degradation and stabilizing both full-length AR and AR-V79. Because AR-V7 is constitutively active and ligand-independent, the resistant clone can sustain AR-driven transcription regardless of the testosterone cycling imposed by BAT, decoupling it from the therapy's mechanism of action10. Truncal *TMPRSS2-ERG* fusion (IHC-confirmed ERG expression at both timepoints) establishes the AR-axis founder event shared by all clones. Truncal ATM deletion (DNA damage-response kinase) and PTEN | The truncal *KMT2C* frameshift identifies a meaningful epigenetic trunk event, but it sits on the trunk and does not distinguish the two branches, so the SNV tree does not inform clonal dynamics |

| | SV tree | SNV tree |
|---|---|---|
| | deletion (PI3K/AKT hyperactivation) create a pan-tumor background of impaired DSB sensing and constitutive pro-survival signaling, but are insufficient alone to prevent BAT lethality in the sensitive clone. The resistant clone's NKX3-1 deletion further de-differentiates it away from AR dependence | |

PSA rose ~148 → ~190 (day 56) then dropped to ~167 (day 84); Ki-67 ~40 → ~20%; tumor volume stable.

##### *Subject 7 (Pair 3): Response*

| | SV tree | SNV tree |
|---|---|---|
| **Topology** | Branched: trunk (Clone 1) → Clone 2 (sensitive), Clone 3 (resistant) | Linear: G → Clone 1 (sensitive) → Clone 2 (resistant); 2 clones |
| **Trunk annotations** | *BRCA2 del, RB1 del, MYCN dup, ERF del, MSH2 del, MSH6 del, ARID1A inv (7 events, heavily rearranged trunk)* | *ETV1* R213C, truncal (present throughout) |
| **Sensitive clone** | Clone 2: ARID1A del, MSH2 inv, MSH6 inv, RAD51B inv (intron7-intron7) | No driver SNVs private to the descendant clone |
| **Resistant clone** | Clone 3: no branch-specific annotations | Not resolved as a separate clone (linear) |
| **Clonal dynamics** | Clone 2 eliminated; Clone 3 expands | Genotype shift within a single lineage: the descendant (clone 2) private genotype rises modestly (pre ~0.36 → post ~0.51); the *ETV1* truncal mutation is retained throughout |
| **Clinical insight** | The trunk carries 7 SV events shared by all clones, | The SNV tree carries only a truncal *ETV1* mutation that |

| | SV tree | SNV tree |
|---|---|---|
| | including BRCA2 deletion, RB1 deletion7,11-13, MYCN duplication, ERF deletion, ARID1A inversion, and bilateral MMR deletion (MSH2, MSH6), making the founder genotype itself heavily HR- and MMR-deficient. The eliminated sensitive clone (Clone 2) acquires additional ARID1A deletion, further MSH2/MSH6 inversions, and a *RAD51B* inversion (intron7-intron7) layered on top of this truncal HRD/MMR background, progressive compound HRR and MMR disruption that renders this lineage acutely vulnerable to BAT-induced DNA double-strand breaks11. The expanding resistant clone (Clone 3) carries no branch-specific events; its BAT resistance is therefore driven by the shared truncal genotype rather than a private resistance mechanism | does not distinguish the two clones, so it is uninformative for the sensitive/resistant split; its dynamics are also much milder than the SV-based branched model |

PSA dropped ~250 → ~120 ng/dL; Ki-67 ~25 → ~45%; modest tumor-volume reduction.

*Subject 10 (Pair 4): Acquired Resistance*

| | SV tree | SNV tree |
|---|---|---|
| **Topology** | Branched: trunk (Clone 1) → Clone 2 (sensitive), Clone 3 (resistant); 3 clones | Branched: trunk (Clone 1) → Clone 2 (sensitive), Clone 3 (resistant); 3 clones |
| **Trunk annotations** | TP53 deletion, CHEK2 intragenic deletion, MDM2 duplication, TMPRSS2-ERG fusion (rescued: robust in | ZFHX3 missense (p.Glu1356Gln), AKT1 E17K; truncal, non-distinguishing |

| | SV tree | SNV tree |
|---|---|---|
| | pre-BAT WGS, single supporting on-BAT read, ERG-positive IHC at both timepoints) | |
| **Sensitive clone** | Clone 2: the most-rearranged cluster (21 translocations, 269 duplications), RAD51B hemizygous deletion, PALB2 deletion | Declining clone: no driver-gene SNVs |
| **Resistant clone** | Clone 3: 11 kb intragenic CUL1 duplication (chr7:148,755,369-148,766,429; hg38) | Emerging clone: no driver-gene SNVs |
| **Clonal dynamics** | Clone 2 disappears, Clone 3 expands on-BAT; whole-sample SV burden falls from 21 translocations / 297 duplications (pre-BAT) to 2 / 56 (on-BAT), a 10.5-fold and 5-fold reduction. In silico downsampling to matched depth shows the translocation drop is partly biological (residual ~2.6-fold beyond coverage, from elimination of the hyper-rearranged Clone 2) while the duplication drop is fully coverage-attributable | Declining leaf falls from cellular fraction ~0.60 to ~0.003; emerging leaf rises ~0.40 to ~0.70, recovering the same clonal-replacement timing as the SV tree |
| **Clinical insight** | SVCFit nominates the intragenic CUL1 duplication as a candidate resistance-associated structural event that SNV-centric analysis would miss (Supplementary Fig. S16). CUL1 is the catalytic backbone of the SCF(SKP2) E3 ubiquitin-ligase complex and a direct c-MYC transcriptional target, so subclonal | The SNV tree resolves the same clonal replacement but carries only one truncal, non-distinguishing driver (ZFHX3) and recovers neither the CUL1 resistance mechanism nor the sensitive-clone HR-pathway deletions. |

| | SV tree | SNV tree |
|---|---|---|
| | amplification at this cell-cycle/ubiquitin-ligase locus is a plausible resistance mechanism, pending clone-resolved transcriptional validation. The eliminated Clone 2 combines extensive genome-wide rearrangement with HR-pathway deletions (RAD51B, PALB2), consistent with BAT selectively eradicating a highly rearranged, HR-deficient subclone while leaving the dominant resistant architecture intact. | |

PSA dropped ~18 → ~9 (nadir, days 28–56), then rose to ~15 by day 84; Ki-67 ~26 → ~55%; tumor volume decreased ~15%.

*Subject 12 (Pair 5): Response (strongest clinical response in the cohort)*

| | SV tree | SNV tree |
|---|---|---|
| **Topology** | Branched: trunk (Clone 1) → Clone 2 (sensitive), Clone 3 (resistant) | Branched: trunk (Clone 1) → Clone 2 (sensitive), Clone 3 (resistant); 3 clones |
| **Trunk annotations** | *PTEN del, TMPRSS2-ERG fusion (intron2-txEnd*; IHC-confirmed ERG expression in both timepoints) | *TP53* P177T (truncal) |
| **Sensitive clone** | Clone 2: CDKN2A del, CDKN2B del | *CTNNB1* S45P |
| **Resistant clone** | Clone 3: *SETD2* inv | No branch-specific driver SNVs |
| **Clonal dynamics** | Clone 2 nearly eliminated; Clone 3 expands from near-zero | Child carrying *CTNNB1* S45P nearly eliminated; second child expands to dominance on-BAT |
| **Clinical insight** | Despite excellent response, the resistant clone (Clone | *CTNNB1* S45P on the sensitive clone contributes |

| | SV tree | SNV tree |
|---|---|---|
| | 3) expanded to dominant; its branch-specific resistance mechanism consists only of *SETD2* inv. *SETD2* encodes the sole H3K36 trimethyltransferase in mammals; its loss paradoxically promotes tolerance to replication stress by slowing fork progression and altering origin firing, allowing cells to survive despite BAT-induced cycling damage rather than collapsing into fork catastrophe. *PTEN deletion and TMPRSS2-ERG fusion are both confirmed truncal* (the latter by IHC showing ERG expression in pre- and on-BAT biopsies), shared by all clones. The sensitive clone (Clone 2) carrying CDKN2A deletion and CDKN2B deletion was nearly eliminated. *CDKN2A/B* encode p16 (CDK4/6 inhibitor), p14ARF (MDM2 inhibitor stabilizing p53), and p15 (CDK4/6 inhibitor); their combined loss abolishes G1/S checkpoints and further destabilizes p53, preventing cells from pausing at G1 to repair BAT-induced DSBs; instead, they race into S-phase with unrepaired damage and enter mitotic catastrophe. The convergence of CDKN2B deletion (SV) and *CTNNB1* | to BAT sensitivity through β-catenin-AR coactivation under supraphysiologic androgen, rather than conferring resistance as it would under AR-targeted therapy |

| | SV tree | SNV tree |
|---|---|---|
| | S45P (SNV) on the same sensitive clone is consistent with compound Wnt/cell-cycle disruption contributing to BAT sensitivity. Although Wnt activation is generally associated with resistance to AR-targeted therapies, the mechanism is distinct under BAT: β-catenin (stabilized by the S45P mutation) acts as a direct AR coactivator through physical interaction with the AR ligand-binding domain, potentiating AR-driven transcription. Under supraphysiologic testosterone, this β-catenin-AR coactivation amplifies the AR transcriptional burden and attendant Topoisomerase IIβ-mediated DSB generation (the very mechanism by which BAT kills cells), compounding the mitotic catastrophe already imposed by loss of G1/S checkpoints (*CDKN2A/B*). | |

PSA dropped ~100 → ~40; Ki-67 ~40 → ~5%; tumor volume decreased ~50%.

##### *Subject 13 (Pair 6): Response*

| | SV tree | SNV tree |
|---|---|---|
| **Topology** | Branched: trunk (Clone 1) → Clone 2 (sensitive), Clone 3 (resistant) | Branched: trunk (Clone 1) → Clone 2 (sensitive), Clone 3 (resistant); 3 clones |
| **Trunk annotations** | *MGA del, RAD51C inv, TMPRSS2-ERG fusion* (IHC- | No driver-gene SNVs |

| | SV tree | SNV tree |
|---|---|---|
| | confirmed ERG expression at both timepoints) | |
| **Sensitive clone** | Clone 2: CDK12 del, RB1 inv, NCOA2 dup, CDK4 dup, MDM2 dup, KRAS dup, RSPO2 dup, MGA inv, ZFHX3 del (9 branch-specific SV events) | Declining clone: no driver-gene SNVs |
| **Resistant clone** | Clone 3: *NCOR1* inv (intron22-intron22) | Emerging clone: no driver-gene SNVs |
| **Clonal dynamics** | Clone 2 nearly eliminated; Clone 3 expands | Three clones: a subclone at low frequency pre-BAT (cellular fraction ≤4%) sweeps to the majority on-BAT while the pre-dominant subclone collapses, with a stable truncal population, corroborating the SV-based clonal replacement |
| **Clinical insight** | The most striking SV-tree case in the cohort: massive clonal replacement with the sensitive clone (Clone 2) carrying 9 branch-specific events: *CDK12 deletion, RB1 inversion, NCOA2 duplication, CDK4 duplication, MDM2 duplication, KRAS duplication, RSPO2 duplication, MGA inversion, and ZFHX3 deletion*, making it the most heavily annotated subclone in the cohort. Truncal *TMPRSS2-ERG* fusion (IHC-confirmed ERG expression at both timepoints) is shared by all clones. The truncal *RAD51C* inv disrupts the BCDX2/CX3 HRR complexes in all cells, creating a pan-tumor HRD | The SNV tree resolves three clones and independently corroborates the timing of the SV-defined clonal replacement, but carries no driver-gene annotations, so it contributes no gene-level mechanism |

| | SV tree | SNV tree |
|---|---|---|
| | background. The near-elimination of Clone 2 despite co-occurring *MDM2*, *KRAS*, *CDK4*, and NCOA2 duplications suggests that, in this clone, cumulative DNA-repair and cell-cycle vulnerability dominated over pro-survival or AR-coactivating signals. Specifically, *CDK12 deletion superimposed on truncal RAD51C disruption*, together with *RB1* inv, may have produced a DSB-repair and checkpoint-control defect that rendered Clone 2 highly susceptible to BAT-induced DNA damage. Thus, the response of Clone 2 emphasizes that the selective fitness of an SV-defined subclone reflects the integrated genotype rather than the expected effect of any single alteration.<br>*NCOR1* inv (intron22-intron22) on the resistant clone is the sole branch-specific annotated event; however, as both breakpoints fall within intron 22, direct protein-coding disruption cannot be inferred from SV breakpoint evidence alone. *NCOR1* is a well-established AR co-repressor whose expression declines with prostate cancer progression, and | |

| | SV tree | SNV tree |
|---|---|---|
| | loss of NCOR1 function is found in ~16% of metastatic prostate tumors, resulting in enhanced AR signaling14; this intronic inversion is a candidate resistance-associated event at the *NCOR1* locus, but whether it affects *NCOR1* expression or splicing requires transcriptomic validation. | |

PSA dropped ~370 → ~30; Ki-67 ~35 → ~5%; tumor volume decreased ~50%.

*Subject 18 (Pair 7): Primary Resistance*

| | SV tree | SNV tree |
|---|---|---|
| **Topology** | Branched: trunk (Clone 1) → Clone 2 (sensitive), Clone 3 (resistant) | Branched: trunk (Clone 1) → Clone 2 (sensitive), Clone 3 (resistant); 3 clones |
| **Trunk annotations** | *PALB2 inv, PTEN inv, NKX3-1 inv, SETD2 del* | No driver-gene SNVs |
| **Sensitive clone** | Clone 2: *ZFHX3* inv (intron5-intron5) | Declining clone: *ZFHX3* V777del (in-frame deletion) |
| **Resistant clone** | Clone 3: no branch-specific annotations | Emerging clone: no driver SNVs |
| **Clonal dynamics** | Clone 2 eliminated; Clone 3 expands from undetectable | Declining clone (pre-dominant) eliminated; emerging clone expands from near-zero to dominant on-BAT |
| **Clinical insight** | Resistance appears driven by the heavily damaged truncal genotype (*PALB2*, *PTEN*, *NKX3-1*, *SETD2*) rather than by branch-specific events; the sole sensitive-clone annotation (*ZFHX3* inv) is of uncertain functional significance | The SNV tree independently places a *ZFHX3* in-frame deletion on the same eliminated clone that the SV tree marks with a *ZFHX3* inversion, an independent convergence of both variant classes on *ZFHX3* in the BAT-sensitive clone |

PSA rose continuously ~52 → ~95; Ki-67 ~55 → ~25%; tumor volume increased ~65% (worst clinical outcome in cohort).

*Subject 20 (Pair 8): Stable Disease*

| | SV tree | SNV tree |
|---|---|---|
| **Topology** | Branched: trunk (Clone 1) → Clone 2 (sensitive), Clone 3 (resistant) | Linear: G → Clone 1 (resistant) → Clone 2 (sensitive); 2 clones |
| **Trunk annotations** | *MLH1 inv, SETD2 inv, PTEN del, RAD51B inv* | Truncal (8 driver SNVs, present throughout): AR L702H, AR T878A, BRCA2 I605fs, *CHD1* S173fs, *KMT2C* Y1362, PIK3R1* N602fs, *PALB2* S382R, *NCOR2* A1824T |
| **Sensitive clone** | Clone 2: TMPRSS2 del (intron1-txEnd), MLH1 del | Descendant subclone: XPO1 V421A, SPOP Y83C (falls from ~97% of tumor cells pre-BAT to undetectable on-BAT) |
| **Resistant clone** | Clone 3: MDM2 dup | Not resolved as a separate clone (linear) |
| **Clonal dynamics** | Clone 2 eliminated; Clone 3 expands | The truncal genotype, which includes BRCA2 fs and both AR ligand-binding-domain mutations, is present throughout; the SPOP Y83C-marked descendant subclone falls from ~97% of tumor cells pre-BAT to undetectable on-BAT, leaving the trunk-only population, ~3% of tumor cells pre-BAT, as the entire tumor on-BAT |
| **Clinical insight** | *PTEN deletion and RAD51B inversion are truncal events* (recovered upon filter correction, both SVs were absent from pre-BAT biopsy 83694 owing to single-caller DELLY-only detection and | This is the one subject whose SNV phylogeny carries information not available from the SV phylogeny. **Truncal BRCA2 fs is a directly actionable PARP-inhibitor indication** |

| | SV tree | SNV tree |
|---|---|---|
| | initially appeared branch-specific). Truncal *RAD51B* inv disrupts the BCDX2 HRR complex (RAD51B-RAD51C-RAD51D-XRCC2) required for stabilizing the RAD51 presynaptic filament; biallelic alteration of *RAD51B* has been confirmed to produce a measurable HRD phenotype15. *MDM2 duplication (a canonical p53-pathway oncogenic event*7,16 that is amplified in ~25% of CRPC17) uniquely defines the resistant clone (Clone 3) as the branch-specific BAT-resistance mechanism: MDM2 ubiquitinates and degrades p53, blocking the apoptotic response to BAT-induced DSBs even on the HRD truncal background; MDM2 additionally stabilizes AR by releasing p53-mediated transcriptional repression of AR target genes, providing a dual survival advantage. *TMPRSS2 deletion and MLH1 deletion on the sensitive clone* (Clone 2), together with truncal *MLH1* inv, create complete biallelic MMR deficiency, a cell that cannot repair BAT-induced DSBs by HRR (*RAD51B* inv) nor correct replication errors by MMR (*MLH1* | **that is present in all cells, including the resistant lineage. The AR ligand-binding-domain mutations L702H and T878A are truncal as well, so every tumor cell carries a receptor altered at the ligand-binding pocket; both are canonical mutations selected by AR-targeted therapy, which this subject had received before enrollment, and both were recovered from filtered calls rather than reported by the caller (Methods, "Rescue of key driver events"). CHD1 fs adds further driver-gene context. SPOP Y83C marks the descendant subclone that BAT eliminates, a sharper localization of the SPOP event than a purely truncal call** |

| | SV tree | SNV tree |
|---|---|---|
| | biallelic) is in extreme genomic distress under testosterone cycling. Truncal MMRd also makes this subject a potential immune checkpoint blockade candidate (TMB-high phenotype) | |

PSA rose continuously ~50 → ~225; Ki-67 ~35 → ~20%; tumor volume slightly increased. *Subject 20 is the single subject in the cohort where the SNV phylogeny carried information not available from the SV phylogeny.*

##### *Subject 31 (Pair 10): Acquired Resistance*

| | SV tree | SNV tree |
|---|---|---|
| **Topology** | Branched: trunk (Clone 1) → Clone 2 (sensitive), Clone 3 (resistant); 3 clones | Branched: trunk (Clone 1) → Clone 2 (sensitive), Clone 3 (resistant); 3 clones |
| **Trunk annotations** | CHD1 hemizygous deletion, focal FOXA1 duplication (exon 2-txEnd; CN ~3 pre-BAT, ~5 on-BAT) | KMT2D nonsense (p.Glu1682Ter), TBX3 frameshift (p.Asn673LysfsTer37) |
| **Sensitive clone** | Clone 2: BRCA2, RB1, ATM deletions (candidate compound HR-pathway defect; RB1 caveat per Supplementary Table S2 RB1 row) | Declining clone: NCOA2 missense (p.Ile988Leu) |
| **Resistant clone** | Clone 3: multiple inactivating ATM inversions (one breakpoint within the ATM gene body, txStart-intron5, truncating the coding sequence), FOXA1 intragenic deletion (exon 2-exon 2), NCOA2, MYC, RAD21 and RSPO2 duplications, RAD51B inversion | Emerging clone: no driver-gene SNVs |
| **Clonal dynamics** | Clone 2 contracts, Clone 3 expands to dominance on-BAT; translocations drop | Declining leaf falls from cellular fraction ~0.82 to ~0.06; emerging leaf rises |

| | SV tree | SNV tree |
|---|---|---|
| | seven-fold (pre-BAT 7 → on-BAT 1), all confined to Clone 2, with a modest duplication increase (16 → 18). Both samples have comparable tumor purity (~0.6), so no downsampling was required | ~0.002 to ~0.83, recovering the same clonal-replacement timing as the SV tree |
| **Clinical insight** | FOXA1 undergoes structural remodeling under BAT: a truncal focal gene-body amplification opposed by a resistant-subclone intragenic deletion, co-occurring with an NCOA2 duplication. SVCFit nominates this FOXA1/NCOA2 remodeling as a candidate resistance mechanism that could redirect AR-cistrome/coactivator activity toward an altered transcriptional program, hypotheses requiring transcriptional and chromatin-state validation; routine gene-level copy-number reporting would summarize FOXA1 as a single dosage state and mask the subclonal gene-body alteration. The contracting Clone 2's BRCA2/RB1/ATM deletions mark a BAT-sensitive, HR-vulnerable population, again consistent with elimination of a highly rearranged HR-deficient subclone. | The SNV tree recovers the same replacement timing and also implicates NCOA2, carrying an NCOA2 missense on the eliminated clone opposite the SV NCOA2 duplication on the expanding clone, but cannot supply the FOXA1 remodeling mechanism. |

PSA dropped ~30 → ~11 (day 56), then rose to ~17 by day 84; Ki-67 ~60 → ~42%; tumor volume decreased ~13%.

##### *Subject 47 (Pair 11): Stable Disease*

| | SV tree | SNV tree |
|---|---|---|
| **Topology** | Branched: trunk (Clone 1) → Clone 2 (sensitive), Clone 3 (resistant) | Branched: trunk (Clone 1, resistant) → Clone 2 (sensitive), Clone 3 (borderline resistant); 3 clones |
| **Trunk annotations** | *PALB2 del, ERF inv* | No driver-gene SNVs |
| **Sensitive clone** | Clone 2: BRCA2 inv, CHD1 inv, PALB2 del, RB1 inv | No driver SNVs on the eliminated child |
| **Resistant clone** | Clone 3: PALB2 del | *KMT2C* Y1963fs (frameshift), on the roughly-flat child |
| **Clonal dynamics** | Clone 2 eliminated; Clone 3 expands from near-zero | One child (pre-dominant) is eliminated; the *KMT2C*-carrying child stays roughly flat; the trunk's own residual becomes the largest distinguishable population on-BAT, so the SNV tree does not cleanly map to a sensitive/resistant split |
| **Clinical insight** | The sensitive clone (Clone 2) carries compound HRR deficiency across 4 DNA-repair genes, *BRCA2 inversion, CHD1 inversion, PALB2 deletion, and RB1 inversion, layered on top of the truncal PALB2 deletion* and *ERF* inv, making it among the most HRR-deficient subclones in the cohort; its elimination is consistent with BAT-induced synthetic lethality in a heavily HRR-compromised | The sole annotation (subclonal *KMT2C* Y1963fs) is not directly actionable, and its subclone is not the population that gains share under BAT, so the SNV tree does not cleanly resolve the sensitive/resistant split |

| | SV tree | SNV tree |
|---|---|---|
| | background11. *CHD1* loss specifically impairs nucleosome eviction at DSB sites required for HR repair factor recruitment, forcing cells toward error-prone NHEJ; CHD1-deficient prostate cancer cells are hypersensitive to DSB-inducing treatments and PARP inhibition in vitro, in vivo, and in patient-derived organoid models18,19. *PALB2* bridges BRCA1 and BRCA2 during DSB repair; with truncal PALB2 deletion (first hit) plus clone-private PALB2 deletion (second hit) plus *BRCA2* inv, this clone's HRR is disrupted at three interdependent nodes of the same repair pathway. Paradoxically, the resistant clone (Clone 3) also achieves biallelic PALB2 deletion (via its own private second hit) yet survives, suggesting either incomplete phenocopying of a functional null state or a compensating epigenetic mechanism not captured by SV analysis | |

PSA rose ~5 → ~17; Ki-67 ~25 → ~70%; tumor volume stable.

##### *Subject 49 (Pair 12): Acquired Resistance*

| | SV tree | SNV tree |
|---|---|---|
| **Topology** | Branched: trunk (Clone 1) → Clone 2 (sensitive), Clone 3 (resistant) | Linear: G → Clone 1 (sensitive) → Clone 2 (resistant); 2 clones |

| | SV tree | SNV tree |
|---|---|---|
| **Trunk annotations** | None | No driver-gene SNVs assigned to the ancestral clone |
| **Sensitive clone** | Clone 2: BRCA2 del, CHD1 del, FOXA1 inv, NCOA2 dup, ONECUT2 dup, APC del, RAD51B inv, SETD2 inv | Not resolved as a separate clone (linear) |
| **Resistant clone** | Clone 3: BRCA2 del (biallelic), RB1 inv, ONECUT2 dup | Descendant (emerging) subclone: no driver-gene SNVs |
| **Clonal dynamics** | Clone 2 nearly eliminated; Clone 3 expands | Genotype shift within a single lineage: the descendant (clone 2) private genotype sweeps from ~1.5% pre-BAT to ~88% on-BAT |
| **Clinical insight** | The trunk carries no annotated driver events; clonal evolutionary history is entirely branch-specific. The sensitive clone (Clone 2) accumulates eight branch-specific events: triple HRR deficiency (BRCA2 deletion, CHD1 deletion, RAD51B inversion), hyper-AR-dependent chromatin remodeling (FOXA1 inversion, NCOA2 duplication), plus ONECUT2 duplication, APC deletion, and SETD2 inversion. The triple HRR deficiency renders Clone 2 acutely vulnerable to BAT-induced DSBs, *CHD1* loss impairs nucleosome eviction at DSBs required for HR-factor access, compounding the *BRCA2* | The SNV phylogeny carries no driver annotations and is uninformative for driver-level clonal dynamics |

| | SV tree | SNV tree |
|---|---|---|
| | and *RAD51B* defects, while FOXA1 inversion and NCOA2 duplication create a hyper-AR-dependent state maximally vulnerable to BAT's testosterone cycling. Clone 2 is thus eliminated from both directions simultaneously. The resistant clone (Clone 3) carries BRCA2 deletion and ONECUT2 duplication as its own branch-specific events, together with *RB1* inv. *ONECUT2 duplication*20,21 drives AR-independent transcription via direct activation of the glucocorticoid receptor (*GR/NR3C1*) and the neuroendocrine splicing factor *SRRM4*, completely decoupling proliferation from androgen levels; *RB1* inv further enables lineage plasticity facilitating the neuroendocrine switch. Despite carrying BRCA2 deletion, the resistant clone is insensitive to BAT's DSB mechanism because it has exited the AR-dependent state entirely, ONECUT2-mediated AR-bypass overrides HRD-mediated sensitivity. | |

PSA dropped ~125 → ~65 (day 56), then rose to ~110 (day 84); Ki-67 ~20 → ~5%; tumor volume slightly increased.

#### S4.3 Aggregate summary across the 11-subject cohort

*SNV-tree informativeness*

| Category | Count | Subjects |
|---|---|---|
| No driver-gene SNVs detected at all | 3/11 | 1, 13, 49 |
| Driver-gene SNVs present but truncal / non-distinguishing only | 3/11 | 6, 7, 10 |
| Driver-gene SNVs distinguish clones, non-actionable | 2/11 | 12, 47 |
| Same gene altered in both trees | 2/11 | 18 (ZFHX3, same clone), 31 (NCOA2, opposing clones) |
| Genuinely clinically informative | 1/11 | 20 (truncal BRCA2 fs and truncal AR L702H/T878A; SPOP Y83C marks a subclone that BAT eliminates, falling from ~97% of tumor cells to undetectable) |

*SV-tree informativeness*

| Category | Count | Subjects |
|---|---|---|
| Clonal architecture resolved with sensitive- and/or resistant-clone driver-gene annotations | 11/11 | All |

*Recurrent SV-driven candidate resistance mechanisms*

| Mechanism | Subject(s) |
|---|---|
| Intragenic *CUL1* duplication | 10 |
| *FOXA1* structural remodeling (opposing dup vs. del) | 31 |
| *MDM2 duplication* | 20 |
| *AURKA duplication* | 6 |
| *NCOR1* inv (intron22-intron22; 1 intronic, functional significance uncertain) | 13 |
| *BRAF duplication + SETD2 deletion* | 1 |
| *ONECUT2 duplication + RB1 inv (AR-independent/NE survival)* | 49 |

| Mechanism | Subject(s) |
|---|---|
| Truncal-genotype-driven (no branch-specific mechanism) | 7, 18 |

Most of these SV-defined mechanisms were not detectable from the corresponding SNV trees. The exceptions are *ZFHX3* (Subject 18) and *NCOA2* (Subject 31), each independently altered by both variant classes: in Subject 18 a *ZFHX3* in-frame deletion (SNV) coincides with the *ZFHX3* inversion (SV) on the same BAT-sensitive clone, and in Subject 31 a *NCOA2* missense (SNV) and a *NCOA2* duplication (SV) implicate the same gene through different alterations in opposing subclones.

*Recurrent SV-defined sensitivity determinants*

| Determinant | Subject(s) |
|---|---|
| Heavy SV rearrangement burden on sensitive clone (≥9 driver-gene SVs) | 10, 13 |
| HR / DNA-repair deficiency on sensitive clone | 7, 10, 31, 47, 49 |
| Compound DNA-repair deficiency (≥4 events on sensitive clone) | 7, 47, 49 |
| Chromatin remodeling disruption (ARID1A del, CHD1 inv) | 7, 47 |
| MMR deficiency (*MSH2*/*MSH6* inv, *MLH1* inv) | 7, 20 |

*Cross-cohort biological themes*

1. **BAT preferentially eliminates heavily rearranged, DNA-repair-deficient subclones.** Visible only on SV trees and consistent with BAT's mechanism of action through supraphysiologic-androgen-induced DNA damage. BAT induces DNA double-strand breaks through AR-mediated co-recruitment of Topoisomerase IIβ at transcription sites and through disruption of AR-dependent DNA replication licensing; cells with multi-gene HRR deficiency cannot repair these breaks and undergo mitotic catastrophe. Across the cohort, sensitive clones eliminated by BAT consistently accumulate HRR gene disruptions (*BRCA2*, *PALB2*, *RAD51B*, *CHD1*, *ATM*, *RAD51C*, *CDK12*), in many cases compounding truncal HRD with additional branch-specific hits, while resistant clones acquire AR-bypass or apoptosis-evasion mechanisms (AURKA, MDM2, ONECUT2 and BRAF duplications) or carry candidate resistance-associated alterations of uncertain mechanism (*NCOR1* inv intron22-intron22, Subject 13) that are largely invisible to SNV-based phylogenetics.
2. **HRD is a predisposing factor for BAT sensitivity but is not deterministic, the depth of HRD disruption and the presence of compensating resistance mechanisms determine outcome.** Sensitive clones with branch-specific or compounding HRR disruption are eliminated in Subjects 7, 10, 31, 47, and 49, consistent with BAT-induced synthetic lethality in HRD backgrounds. *CHD1* loss

specifically impairs nucleosome eviction at DSBs required for HR factor access, sensitizing cells to DSB-inducing therapy even in the absence of canonical *BRCA1/2* alterations18,19. Conversely, in Subject 49, the resistant clone carries biallelic BRCA2 deletion yet expands, ONECUT2 duplication20,21 provides a complete AR-bypass mechanism through neuroendocrine reprogramming that overrides HRD-mediated sensitivity. The implication is that a single AR-bypass event can neutralize even severe HRD vulnerability. Patterns derived from individual subjects should be interpreted with appropriate caution.

3. **Candidate resistance mechanisms are diverse, predominantly SV-defined, and mechanistically distinct.** They span: (i) MAPK pathway activation (BRAF duplication, Subject 1); (ii) AR splice-variant stabilization via kinase-mediated splicing/proteasome reprogramming (AURKA duplication → AR-V7, Subject 69,10); (iii) candidate *NCOR1* locus alteration (*NCOR1* inv intron22-intron22, Subject 13; both breakpoints intronic, coding disruption not confirmed; functional significance requires transcriptomic validation14); (iv) p53-pathway bypass (MDM2 duplication, Subject 20); (v) AR-independent neuroendocrine reprogramming (ONECUT2 duplication, Subject 4920,21); and (vi) epigenetic replication-stress tolerance (SETD2 inv/del, Subjects 1, 12). None of these SV-defined events was detectable from the corresponding SNV trees. On the SNV side, Subject 20 additionally carries the truncal AR ligand-binding-domain mutations L702H and T878A, recovered from filtered calls; being truncal they mark the founder genotype rather than a BAT-resistant branch.
4. **Recurrent *KMT2C* loss-of-function across the SNV trees** (Subjects 6, 20, 47) may define an epigenetic subtype, but does not require phylogenetic analysis to detect.

#### S4.4 Paired SV and SNV phylogeny diagrams (all 11 subjects)

For visual comparison, the SVCFit-inferred SV phylogeny and the PICTograph SNV phylogeny for each of the 11 subjects are shown side by side below (SV phylogeny above, matched SNV phylogeny below). SV trees annotate prostate-cancer driver structural events on each clone's edge with clone-proportion pie charts for the pre- and on-BAT biopsies; SNV trees annotate driver point mutations, with amino-acid change, on the incoming edge of the cluster carrying them (germline root, G). Structural events are abbreviated Inv (inversion), Dup (duplication or copy-number gain), Del (deletion or copy-number loss), fusion (gene fusion) and hemi (hemizygous); exact breakpoint coordinates are given in the per-subject tables above. In both trees clones are numbered in tree order (trunk 1; declining clone 2, drawn left; emerging clone 3, drawn right), but the clones themselves are inferred independently by each method and a given number does not denote the same set of cells in the SV and SNV trees. Subjects 10 and 31 are also shown in main-text Fig. 5 (SV tree only). The per-subject paired SV and SNV phylogenies are shown in Supplementary Figures S14-1 through S14-11, and the pre- versus on-BAT SNV co-clustering for Subjects 13 and 31 in Figure S15.

#### Supplementary Note S5: SVCF estimation in non-diploid genomic regions

**Cross-reference to main text.** The non-diploid adjustment expressions $BPC' = BPC \times SVCF/(SVCF + ACR - 1)$ and $BEC' = BEC \times (2 - SVCF)/(1 + ACR - SVCF)$ appear as Eqs. 3 and 4 of the main text. The ACR inference relation $ACR = VAF_{het}/(1 - VAF_{het})$ derived in this Note (Eq. S60) appears as Eq. 7 of the main-text Methods. The closed-form fixed points (Eqs. S67 and S69 below) appear as Eqs. 5 and 6 of Results. The SV-CNV ordering criterion (sign of Eq. 5, main text) is proved formally in Supplementary Note S2.3 and illustrated in Supplementary Fig. S2. The operational phasing pipeline (filtering tumor BAMs to breakpoint-supporting reads with samtools and running bcftools mpileup over heterozygous SNP positions) is described in the "Haplotype-aware SV phasing and zygosity inference" paragraph of the main-text Methods.

Structural variants frequently occur in genomic regions affected by copy-number alterations. This note describes how SVCFit estimates SVCF in non-diploid regions by integrating allele-specific copy number, structural variant zygosity, and phasing information to account for overlapping CNV and SV events.

In tumors, structural variants (SVs) are not confined to diploid regions. They often overlap with focal or large copy number alterations. To accurately estimate SVCF in these cases, the breakpoint read counts (BPRC) and breakend read counts (BERC) must be appropriately adjusted. This adjustment process requires inference of SV zygosity, SV and CNV phasing, as well as determination of the relative timing or ordering between the SV and the CNV events. SVCFit implements this by integrating data from an external copy number caller FACETS[2] and heterozygous germline single nucleotide polymorphism (SNP) calls from GATK4[3].

**SV zygosity and phasing.** The mechanics of SV zygosity inference and haplotype assignment from breakpoint-supporting reads at proximal heterozygous SNPs, and the SNP detection pipeline (GATK4[3] HaplotypeCaller, bcftools[4] heterozygous filter, samtools[4] breakpoint-read filter, bcftools mpileup), are described in the "Haplotype-aware SV phasing and zygosity inference" and "Germline heterozygous SNP detection" subsections of the main-text Methods. The illustrative schematic appears as main-text Fig. 1G-I.

**CNV phasing, ACR derivation.** CNV phasing was inferred through allele copy ratio (ACR) analysis, with ACR values estimated from heterozygous-SNP read counts. The closed-form result $ACR = VAF_{het}/(1 - VAF_{het})$ appears as main-text Eq. 7; the full algebraic derivation is given below.

$$VAF_{het} = \frac{alt}{ref + alt} = \frac{ACR \cdot cell}{ACR \cdot cell + 1 \cdot cell} = \frac{ACR}{ACR + 1} \quad (S54)$$

This equation can be rewritten into:

$$VAF_{het} = \frac{ACR}{ACR + 1} \quad (S55)$$

$$VAF_{het} \times (ACR + 1) = ACR \quad (S56)$$

$$VAF_{het} \times ACR + VAF_{het} = ACR \quad (S57)$$

$$VAF_{het} = ACR - VAF_{het} \times ACR \quad (S58)$$

$$VAF_{het} = ACR \times (1 - VAF_{het}) \quad (S59)$$

$$\frac{VAF_{het}}{1 - VAF_{het}} = ACR \quad (S60)$$

If we plug in the expression of VAF,

$$\frac{VAF_{het}}{1 - VAF_{het}} = ACR \quad (S61)$$

$$\frac{\frac{alt}{alt + ref}}{1 - \frac{alt}{alt + ref}} = ACR \quad (S62)$$

$$\frac{\frac{alt}{alt + ref}}{\frac{ref}{alt + ref}} = ACR \quad (S63)$$

$$\frac{alt}{ref} = ACR \quad (S64)$$

Because the calculation relies on the variant allele frequency (VAF) of heterozygous SNPs, the allele copy ratio (ACR) inherently tracks the alternate allele. For events affecting the reference allele, the ACR represents the reciprocal of the observed ratio (Eq S64). This creates directional ambiguity: as shown in Eq S64, an ACR $\geq 1$ indicates either alternate allele amplification or reference allele loss, while an ACR $\leq 1$ signifies reference amplification or alternate loss. To resolve this ambiguity and distinguish between gain and loss events, we integrated CNV classifications from FACETS[2].

**SVCF estimation:** Based on the inferred zygosity and phasing of an SV and the phasing of overlapping CNVs, SVCFit determines whether the CNV changes the total breakpoint read count or the total breakend read count of the SV. If the breakpoint read count has changed, it is adjusted according to Eq S65 (main-text Eq. 3),

$$BPRC' = BPRC \times \frac{SVCF}{SVCF + ACR - 1} \quad (S65)$$

If the breakend read count has changed, it is adjusted according to Eq S66 (main-text Eq. 4),

$$BERC' = BERC \times \frac{2 - SVCF}{1 + ACR - SVCF} \quad (S66)$$

SVCFit estimates SVCF using Eq S65 and Eq S66 with iterative optimization. For changes in both breakpoint and breakend read counts, the optimization converges to stable solutions (S67, S69), as shown in Supplementary Note S2.

When breakpoint read count changes (main-text Eq. 5),

$$SVCF = \frac{2 \times BPC - BEC \times (ACR - 1)}{BPC + BEC} \quad (S67)$$

When breakend read count changes,

$$\left(BPC + \frac{BEC}{2}\right) SVCF^2 - (3BPC + BPC \times ACR + BEC)SVCF + 2BPC(1 + ACR) = 0 \quad (S68)$$

Eq S68 is a quadratic equation with one feasible solution,

$$SVCF = \frac{BPC + BPC \times ACR}{BPC + BEC} \quad \text{(Supplementary Note S2)} \quad (S69)$$

**SV/CNV ordering:** When an SV and CNV overlap, the SVCF calculation depends on the relative timing of the two events. If the CNV occurred before the SV, the breakpoint read count is the same as in a diploid region, while the breakend read count changes. In this case, SVCF can be calculated with Eq S69. Conversely, if the SV occurred before the CNV, the breakend read count is the same as in a diploid region, while the breakpoint read count changes. Then SVCF can be calculated with Eq S67. Importantly, the solution to Eq S67 will be negative if the CNV occurred before the SV. So the order of an overlapping SV and CNV can be inferred from the sign of Eq S67 (sign of main-text Eq. 5; Supplementary Note S2; Supplementary Fig. S2). In practice this distinction depends on having sufficient breakpoint and breakend coverage to separate small numerator differences from sampling noise.

**Estimation on hemizygous chromosomes.** The derivations above read the allele copy ratio (ACR) from the allele frequencies of heterozygous germline SNPs near the structural variant (main-text Eq. 7). A hemizygous chromosome has none: the X chromosome of a male subject carries a single allele, so no heterozygous SNP exists and the ACR is undefined. On such a chromosome SVCFit replaces the ACR-based adjustment with $\bar{R}$, the mean number of copies of the affected locus per cell, measured directly from read depth relative to the matched normal, which is itself single-copy there. $\bar{R}$ is the same quantity that equals 2 in a diploid region and enters the general conversion of main-text Eq. 1; on a hemizygous chromosome it is estimated from depth rather than fixed by ploidy (main-text "Estimation on hemizygous chromosomes"). Writing VAF for the structural-variant allele frequency (main-text Eq. 2), the cellular fraction takes one of three closed forms, according to whether a copy-number change overlaps the SV and, if so, its order relative to the SV.

**Copy-neutral.** When no copy-number change overlaps the SV, each cell carries a single copy of the locus, $\bar{R} = 1$, and the cellular fraction is the allele frequency itself, SVCF = VAF.

**Copy-number change before the SV.** When a copy-number change precedes the SV, the SV lies on a locus already present at $\bar{R}$ copies per cell and its supporting reads are diluted in that proportion, giving SVCF = $\bar{R}$ × VAF.

**SV before the copy-number change.** When the SV precedes the copy-number change, the rearranged copy is itself amplified and the count of SV-supporting reads is inflated by the copies gained; balancing the read counts gives SVCF = $\bar{R}$ × VAF − ($\bar{R}$ − 1). The sign of this expression also resolves the order: only one ordering yields a cellular fraction in [0, 1], which is how SVCFit assigns the SV-CNV order on a hemizygous chromosome when it is not otherwise known.

**Two limits bound the hemizygous estimator.** A tandem duplication is not point-identifiable from a single hemizygous locus, because the number of tandem copies r cannot be separated from the cellular fraction; SVCFit reports ($\bar{R}$ − 1)/(r − 1) evaluated at r = 2, an upper bound on SVCF rather than a point estimate. A deletion shorter than the copy-number segmentation window has no depth contrast to corroborate it and is checked against the read evidence directly. Both limits are stated where the corresponding estimates are reported.

#### Supplementary Note S6: Robustness of SVCF estimation to breakpoint read depth

The definition of SVCF assumes that average read depth at structural variant breakpoints and breakends is approximately equal. Here, we evaluate the robustness of SVCF estimation to deviations from this assumption using simulated tumor samples across a wide range of SV cellular fractions and sequencing depths.

The SVCFit framework is built around a definition of variant allele frequency (VAF) that is a ratio of breakpoint read counts and the sum of breakpoint and breakend read counts for an SV, summed over $N$ cells from a sample (Eq S9). This ratio can be converted into a ratio of breakpoint counts and the sum of breakpoint and breakend counts, with the observation that the average read depth at an SV's breakpoints and breakends are approximately equal (Eq S14).

To validate this assumption, we performed simulations using VISOR[1], generating 1,000 replicates of a clonal tumor sample (six SVs per replicate) at 50X sequencing coverage. SV tumor purity (fraction of cells carrying the SV) was randomly assigned between 10% and 100%. We computed the maximum read depth at every SV breakpoint (including two flanking base pairs) using samtools[4] and then plotted total read depth against tumor purity and overlaid a Pearson correlation trendline using `geom_smooth(method = "glm")` from ggplot2[5] (v3.5.1).

Theoretically, increasing tumor purity shifts the cell population, increasing the fraction of cells containing SVs (yielding breakpoint reads) while decreasing the fraction of normal cells (yielding breakend reads). Consequently, if the total read depth remains constant despite this shift, the read generation rates for both types must be equivalent. Fig. S1 shows our simulation confirmed that total read depth remained stable across varying purities (Pearson correlation). This result supports the assumption that average read depth is approximately equal at the breakpoint and breakend positions despite SV tumor purities (Supplementary Note S8).

#### Supplementary Note S7: Mathematical properties of SVCF under tandem duplications and deletions

This note establishes key mathematical properties of the SVCF formulation for tandem duplications and deletions. We derive multiplicative identities and relationships between breakpoint counts, breakend counts, and copy number that enable SVCF estimation from observable sequencing data. The closed-form $\overline{R} = 2(r-1)/r$ formula for tandem duplications listed in Table 1 of the main text is derived in Section S7.3 below.

##### S7.1: Tandem duplication multiplicative identity

**Table S6. Variables for tandem duplications.** $N_{SV}$ is the number of cells with a tandem duplication; SVCF is cellular fraction for a tandem duplication; $N$ is the number of cells in a bulk sequenced sample; $R_{gain}$ is the number of break intervals copied by a tandem duplication; R is the total number of break intervals; BEC is breakend counts; BPC is breakpoint counts.

| $N_{SV}$ | SV CF | N | $R_{gain}$ | R | BEC | BPC |
|---|---|---|---|---|---|---|
| 1 | P | 1/P | g | 2/P + g | 4/P | g |
| 2 | P | 2/P | g | 4/P + 2g | 8/P | 2g |
| … | … | … | … | … | … | … |
| K | P | K/P | g | 2K/P + Kg | 4K/P | Kg |
| K+1 | P | (K+1)/P | g | 2(K+1)/P + (K+1)g | 4(K+1)/P | (K+1)g |

Proof.

Given $P \in [0,1], g \in \mathbb{N}, N_{SV} = 1$. Then,

$$\frac{R}{\frac{BEC}{2} + BPC} = \frac{\frac{2}{P} + g}{\frac{4}{2P} + g} = \frac{\frac{2}{P} + g}{\frac{2}{P} + g} = 1$$

Assume $\frac{R}{\frac{BEC}{2}+BPC} = 1$ is true given $N_{SV} = K$. Then,

Given $P \in [0,1], g \in \mathbb{N}, N_{SV} = K + 1$,

$$\frac{R}{\frac{BEC}{2} + BPC} = \frac{\frac{2(K+1)}{P} + (K+1)g}{\frac{4(K+1)}{2P} + (K+1)g} = \frac{\frac{2(K+1)}{P} + (K+1)g}{\frac{2(K+1)}{P} + (K+1)g} = 1$$

By induction, $\frac{R}{\frac{BEC}{2}+BPC} = 1 \mid P \in [0,1], g \in \mathbb{N}$ applies to all $N_{SV}$ for tandem duplications. ■

#### S7.2: Deletion multiplicative identity

The multiplicative identity for deletions follows directly from the tandem duplication case (S7.1) by fixing the number of break intervals and breakpoint counts (Table S6). A full induction table is therefore omitted for brevity, as no additional assumptions are introduced.

By induction, $\frac{R}{\frac{BEC}{2}+BPC} = 1 \mid P \in [0,1]$ and $\overline{R} = 2$ applies to all $N_{SV}$ for deletions.

#### S7.3: $\overline{R}$ formulation for tandem duplication

> **Cross-reference to main text.** Table 1 of the main text lists $\overline{R} = 2(r-1)/r$ for tandem duplications. This $r$-dependent formula is derived from the induction below.

The expression for R in tandem duplications follows directly from the multiplicative identity derived above (S7.1) by rearrangement of terms. We therefore omit the full induction table, as no additional assumptions are introduced.

By induction, $\frac{\overline{R}-2}{\overline{R}} = VAF \mid P \in [0,1], T \in \mathbb{N}$ applies to all $R_{gain}$ for tandem duplications.

#### Supplementary Note S8: SV breakpoint and breakend read depth across tumor purities

This note provides an analytical justification for the observation that average read depth at structural variant breakpoints and breakends remains approximately constant across tumor purities. This property supports the use of breakpoint- and breakend-based read counts in SVCF estimation.

To support this assumption, we build on our result from Supplementary Note S6, which showed that total read depth at the start and end of six SVs remained approximately constant across varying SV tumor purities in simulated tumor replicates. Without loss of generality, we equate the total read depth of an SV in a sample with $N$ cells and tumor purity $SV_{TP1}$ to that in a sample with the same number of cells and a different SV tumor purity $SV_{TP2}$. On both sides, total read depth is expressed in terms of the average read depth at the breakpoints and breakends of an SV, and the read depth at the corresponding genomic positions in cells lacking the SV. Simplifying both sides shows that the average read depth at SV breakpoints and breakends is approximately equal, regardless of SV tumor purity.

$$(N \times SV_{TP1} \times \overline{RD}_{bp}) + (N \times SV_{TP1} \times \overline{RD}_{be}) + (N \times (1 - SV_{TP1}) \times \overline{RD}_{be})$$

$$\approx (N \times SV_{TP2} \times \overline{RD}_{bp}) + (N \times SV_{TP2} \times \overline{RD}_{be}) + (N \times (1 - SV_{TP2}) \times \overline{RD}_{be})$$

$$(N \times SV_{TP1} \times \overline{RD}_{bp}) + (2N - N \times SV_{TP1}) \times \overline{RD}_{be}$$

$$\approx (N \times SV_{TP2} \times \overline{RD}_{bp}) + (2N - N \times SV_{TP2}) \times \overline{RD}_{be}$$

$$(N \times SV_{TP1} \times \overline{RD}_{bp}) - (N \times SV_{TP2} \times \overline{RD}_{bp})$$

$$\approx (2N - N \times SV_{TP2}) \times \overline{RD}_{be} - (2N - N \times SV_{TP1}) \times \overline{RD}_{be}$$

$$\overline{RD}_{be} \times N(SV_{TP1} - SV_{TP2}) \approx \overline{RD}_{bp} \times N(SV_{TP1} - SV_{TP2})$$

$$\overline{RD}_{be} \approx \overline{RD}_{bp}$$

**Variable Definitions** - $N$: Total number of cells in a sample. - $SV_{TP1}$: SV tumor purity in sample 1. - $SV_{TP2}$: SV tumor purity in sample 2. - $\overline{RD}_{bp}$: Average read depth at a breakpoint position of an SV. - $\overline{RD}_{be}$: Average read depth at a breakend position of an SV.

### Supplementary Figures

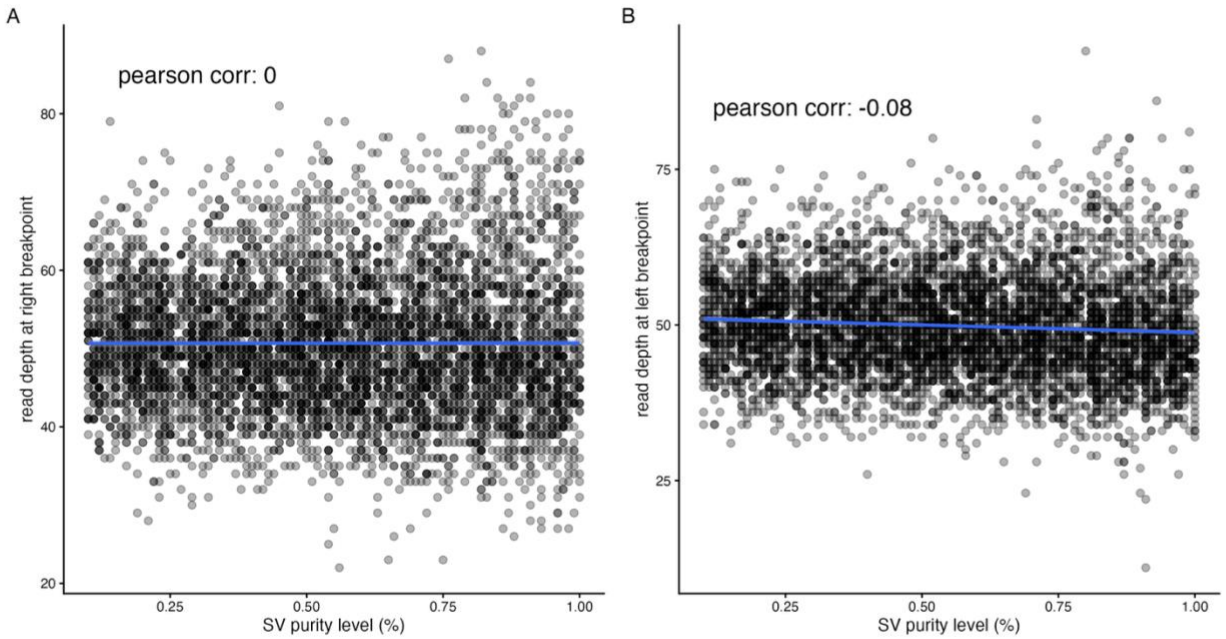


**Figure S1. Stability of read depth across SV tumor purities.** Total read depth at structural variant (SV) breakpoints and breakends remains stable across varying SV tumor purities in 1,000 clonal tumor simulations (six SVs per simulation). Scatter plots show total read depth as a function of SV tumor purity, with fitted linear trend lines and corresponding Pearson correlation coefficients. (A) Total read depth at the right (downstream) breakpoint of each SV. (B) Total read depth at the left (upstream) breakpoint of each SV.

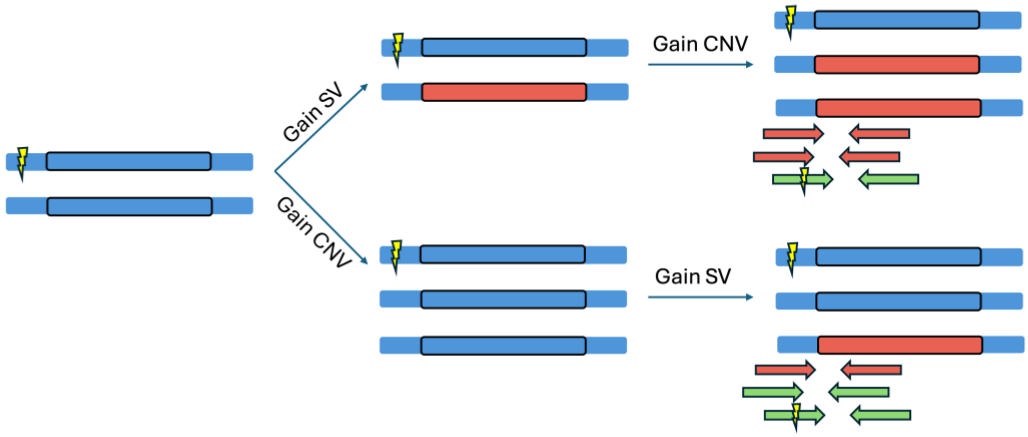


**Figure S2. Schematic of relative SV-CNV ordering inference.** Distinguishing the relative ordering of somatic SVs and copy-number variants (CNVs). The schematic illustrates expected differences in read counts depending on whether an SV occurred before or after a copy-number gain. In the early SV scenario (top), an SV precedes genomic amplification, resulting in an increased proportion of SV-supporting reads. In the late SV scenario (bottom), a copy-number gain occurs first, followed by an SV on a single allele, producing a diluted read profile in which SV-supporting reads constitute a smaller fraction of total coverage. The sign of main-text Eq. 5 distinguishes these two cases analytically (Note S2.3); in practice this distinction depends on having sufficient breakpoint and breakend coverage.

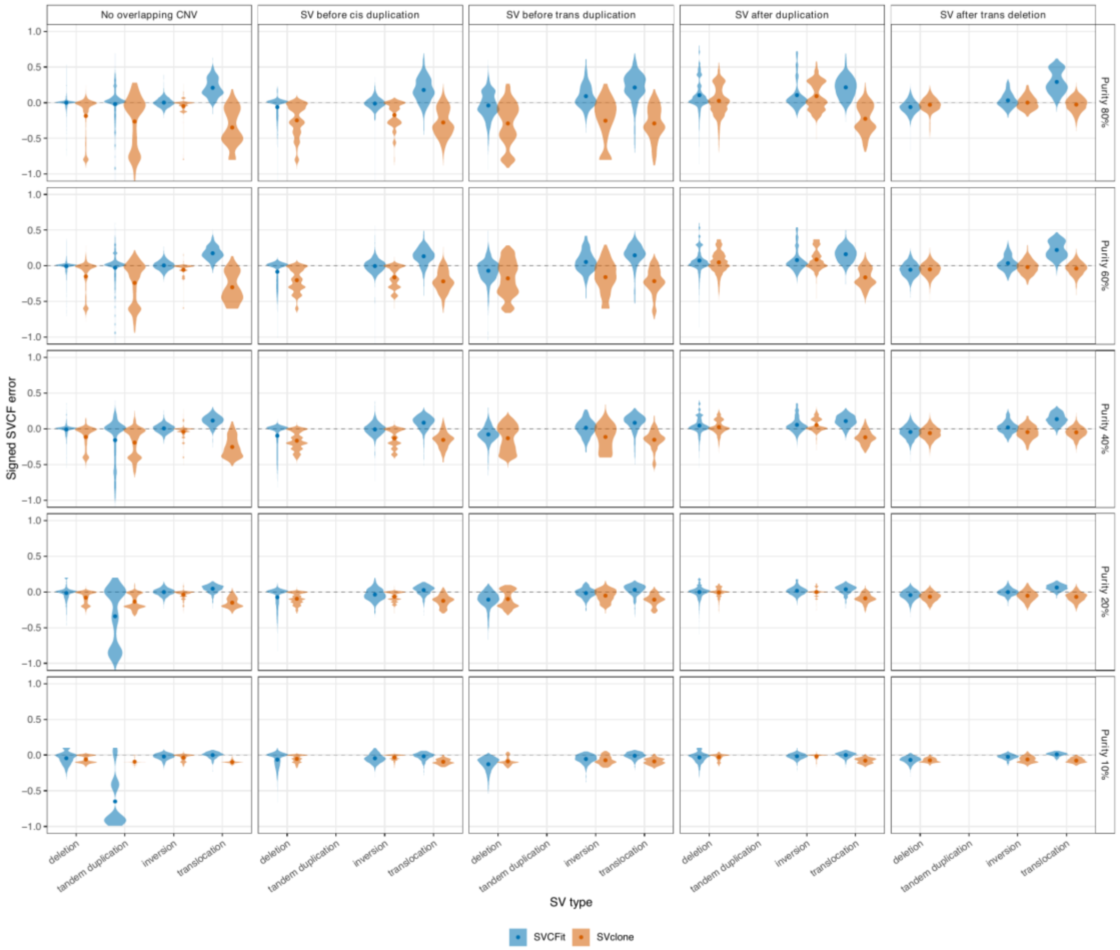


**Figure S3.** Signed SVCF error by structural-variant type across purity and SV-CNV overlap (autosomes + chrX). Per-SV signed-error distributions (violins) for SVCFit (blue) and SVclone (orange), faceted by tumor purity (rows) and SV-CNV-overlap experiment (columns: no overlapping CNV, SV before cis duplication, SV before trans duplication, SV after duplication, SV after trans deletion), split by SV type on the x-axis. chrX (hemizygous) deletions, tandem duplications, and inversions are pooled into the baseline, before-cis-duplication and after-duplication columns; the trans experiments stay autosome-only. Overlaid points are each method's mean signed error with bootstrap 95% CI (across 30 replicates); the dashed line at zero is exact. Signed SVCF error is truth minus estimate, so a positive value means the method underestimates the cancer-cell fraction. Per-cell effect sizes, confidence intervals, and p-values are in the companion statistics table (Supplementary Table S7).

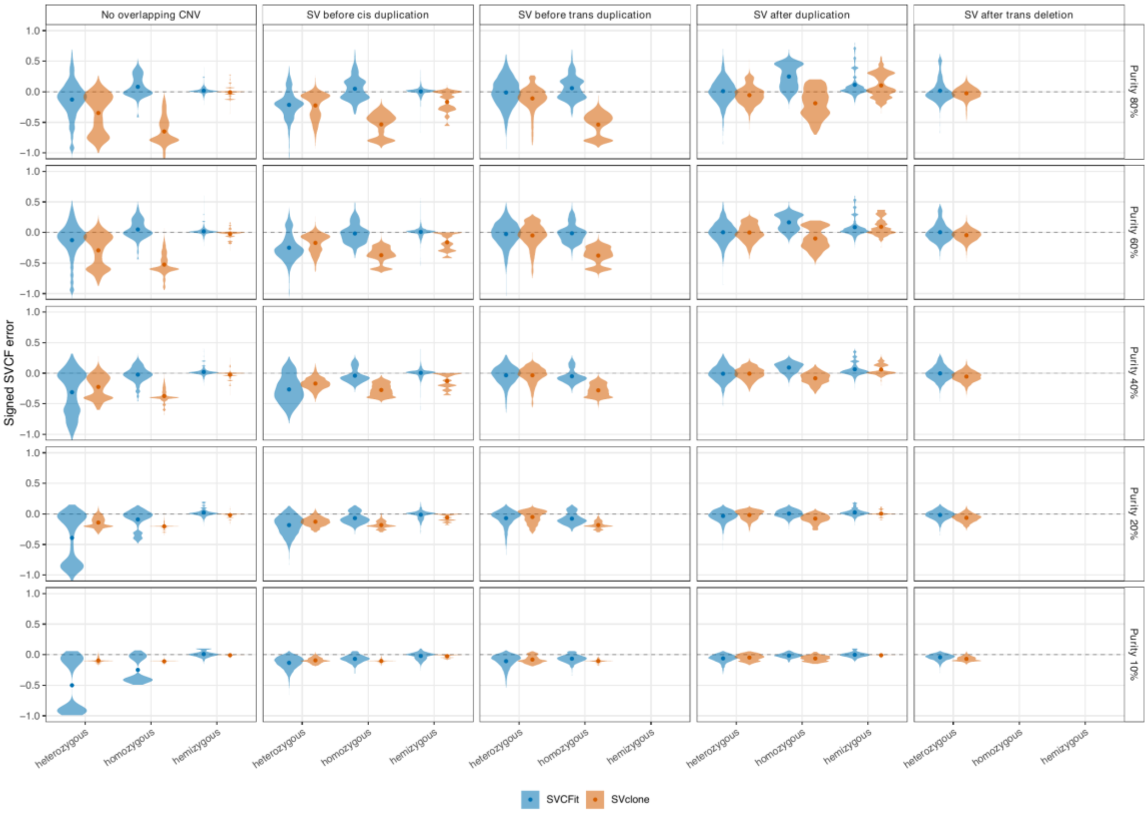


**Figure S4.** Signed SVCF error by zygosity across the full purity range, including hemizygous (chrX) loci. As in Supplementary Fig. S3 but split on the x-axis by zygosity, heterozygous, homozygous (autosomal), and hemizygous (chrX), faceted by purity (rows) and SV-CNV-overlap experiment (columns). Hemizygous is a single-copy locus for which heterozygous versus homozygous is undefined; it is the chrX analog. Violins are the per-SV signed-error distributions; overlaid points are each method's mean signed error with bootstrap 95% CI across 30 replicates; the dashed line at zero is exact. Per-cell effect sizes, confidence intervals, and p-values are in the companion statistics table (Supplementary Table S8).

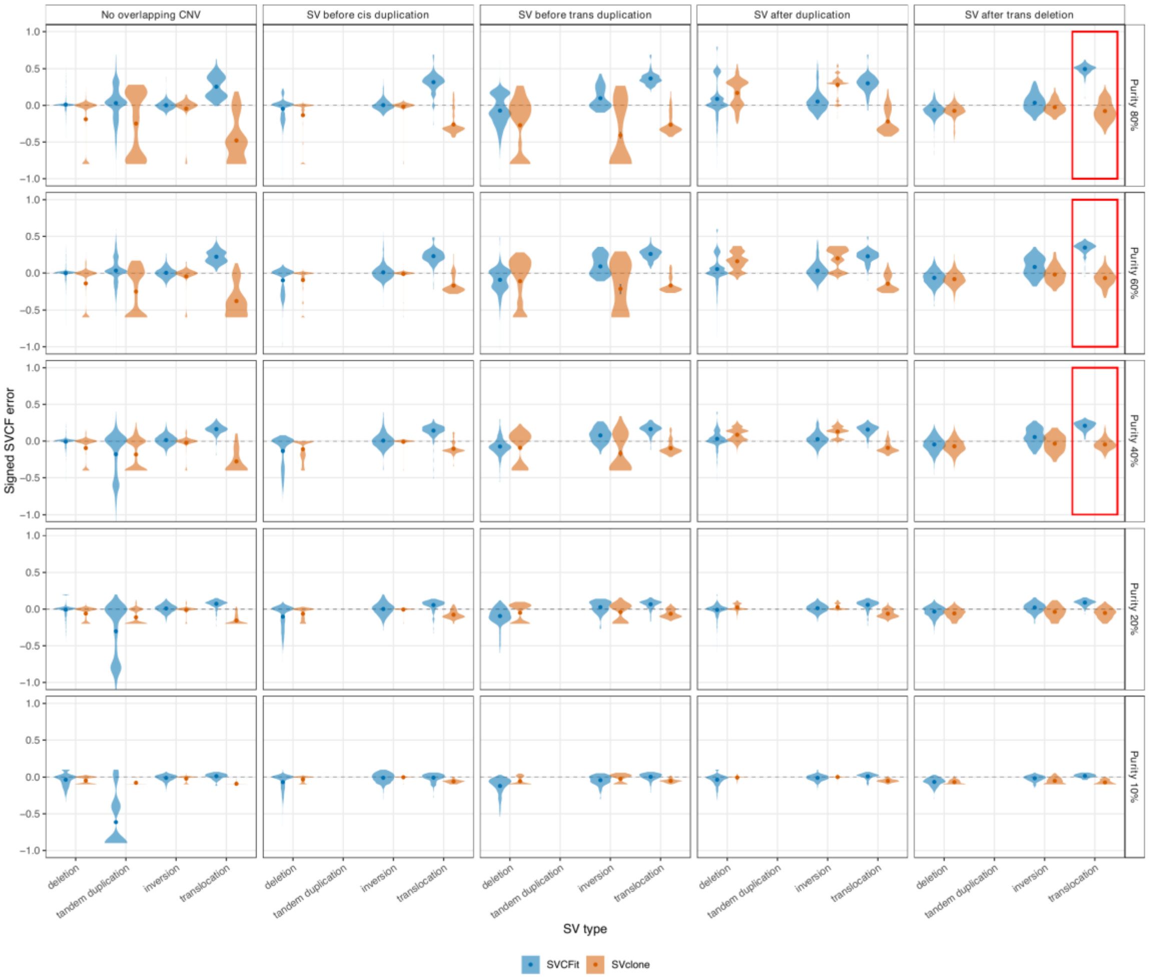


**Figure S5.** Signed SVCF error for clonal SVs only (autosomes + chrX). As in Supplementary Fig. S3 but restricted to clonal (truncal) structural variants, split by SV type across purity (rows) and SV-CNV overlap (columns). Red rectangles mark SVCFit's largest clonal failure: translocations in the SV-after-trans-deletion configuration at 40-80% purity, where SVCFit increasingly underestimates the cellular fraction as purity rises while SVclone remains near the truth. Violins are per-SV signed error; overlaid points are each method's mean signed error with bootstrap 95% CI across 30 replicates; the dashed line at zero is exact. Per-cell effect sizes, confidence intervals, and p-values are in the companion statistics table (Supplementary Table S9).

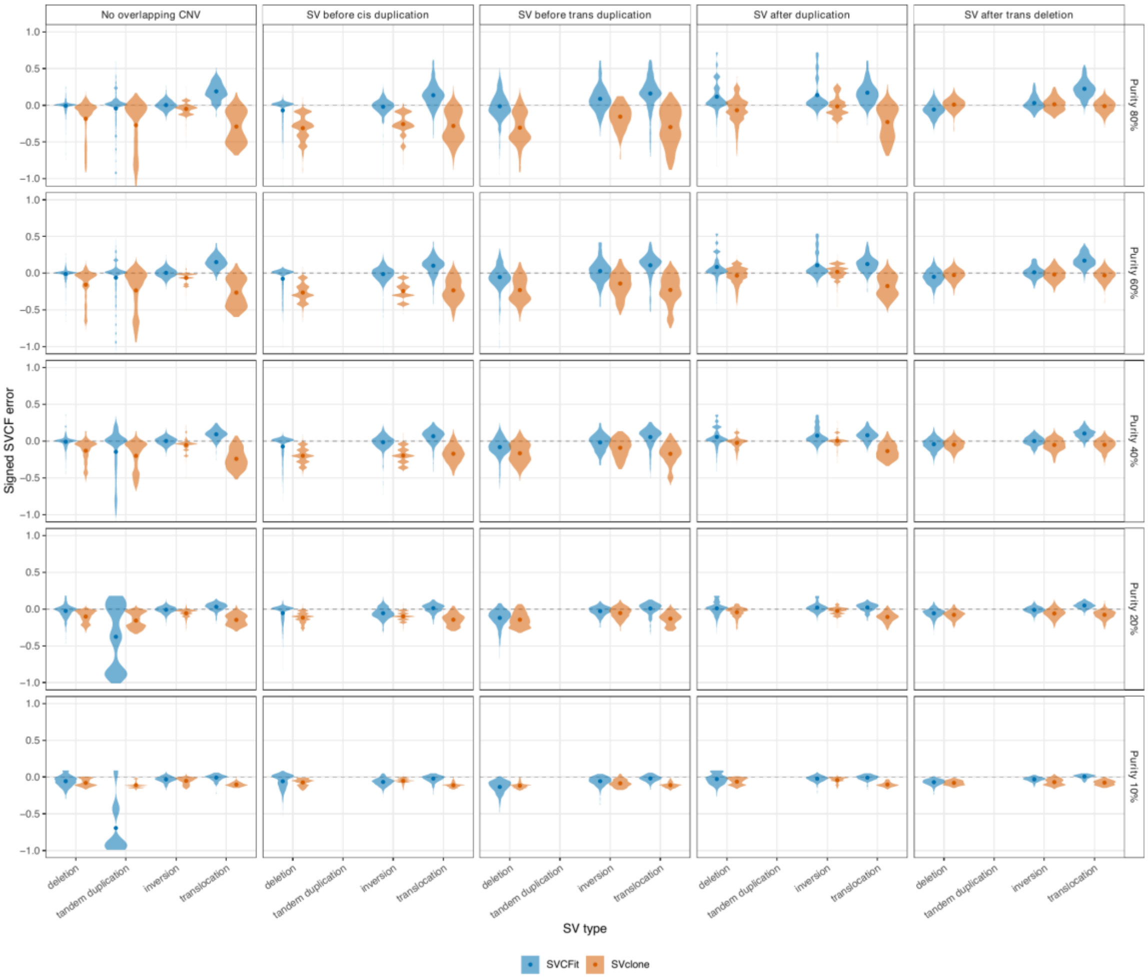


**Figure S6.** Signed SVCF error for subclonal SVs only (autosomes + chrX). As in Supplementary Fig. S5 but restricted to subclonal structural variants, the tight-margin decisions where CCF errors most affect downstream clone assignment. Split by SV type across purity (rows) and SV-CNV overlap (columns); violins are per-SV signed error; overlaid points are each method's mean signed error with bootstrap 95% CI across 30 replicates; the dashed line at zero is exact. Per-cell effect sizes, confidence intervals, and p-values are in the companion statistics table (Supplementary Table S10).

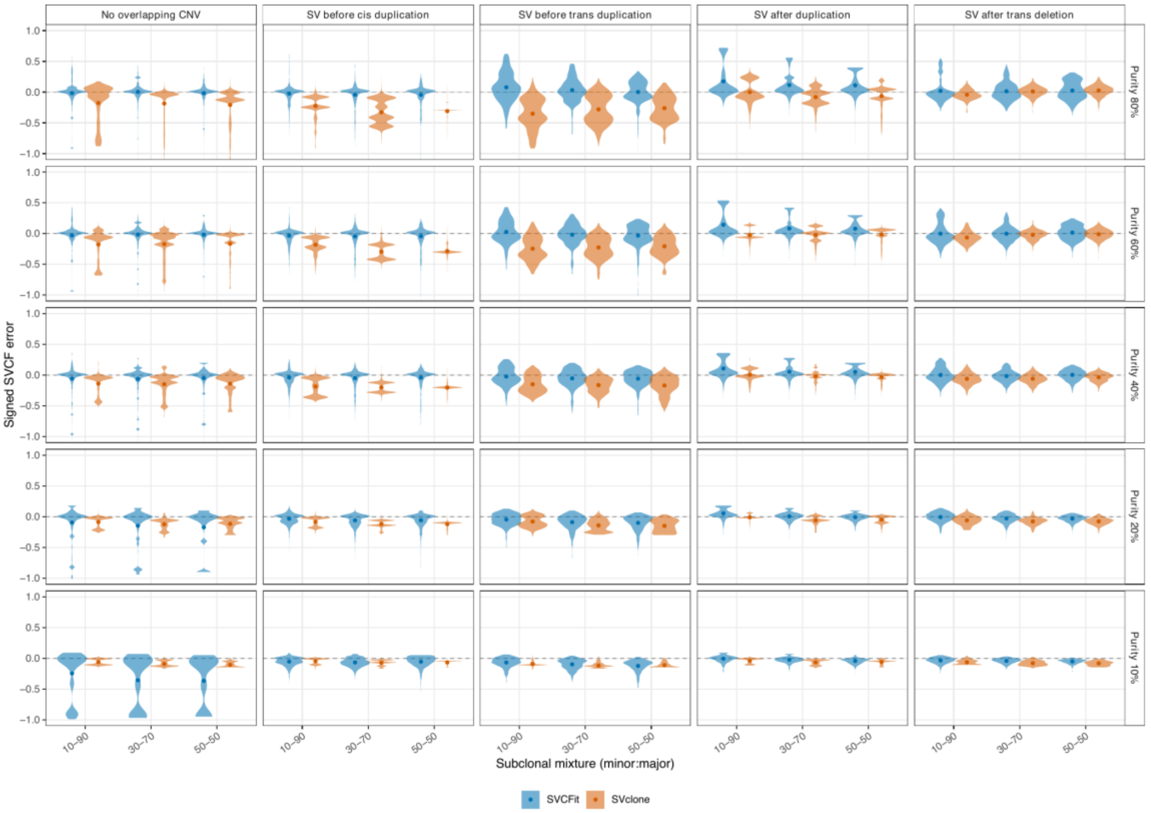


**Figure S7.** Signed SVCF error for subclonal SVs by subclonal mixture setting (autosomes + chrX). Subclonal structural variants split on the x-axis by the minor:major subclonal mixture (10-90, 30-70, 50-50), faceted by purity (rows) and SV-CNV-overlap experiment (columns). Tighter mixtures place the two subclones' CCFs closer together, shrinking the assignment margin. Violins are per-SV signed error; overlaid points are each method's mean signed error with bootstrap 95% CI across 30 replicates; the dashed line at zero is exact. Per-cell effect sizes, confidence intervals, and p-values are in the companion statistics table (Supplementary Table S11).

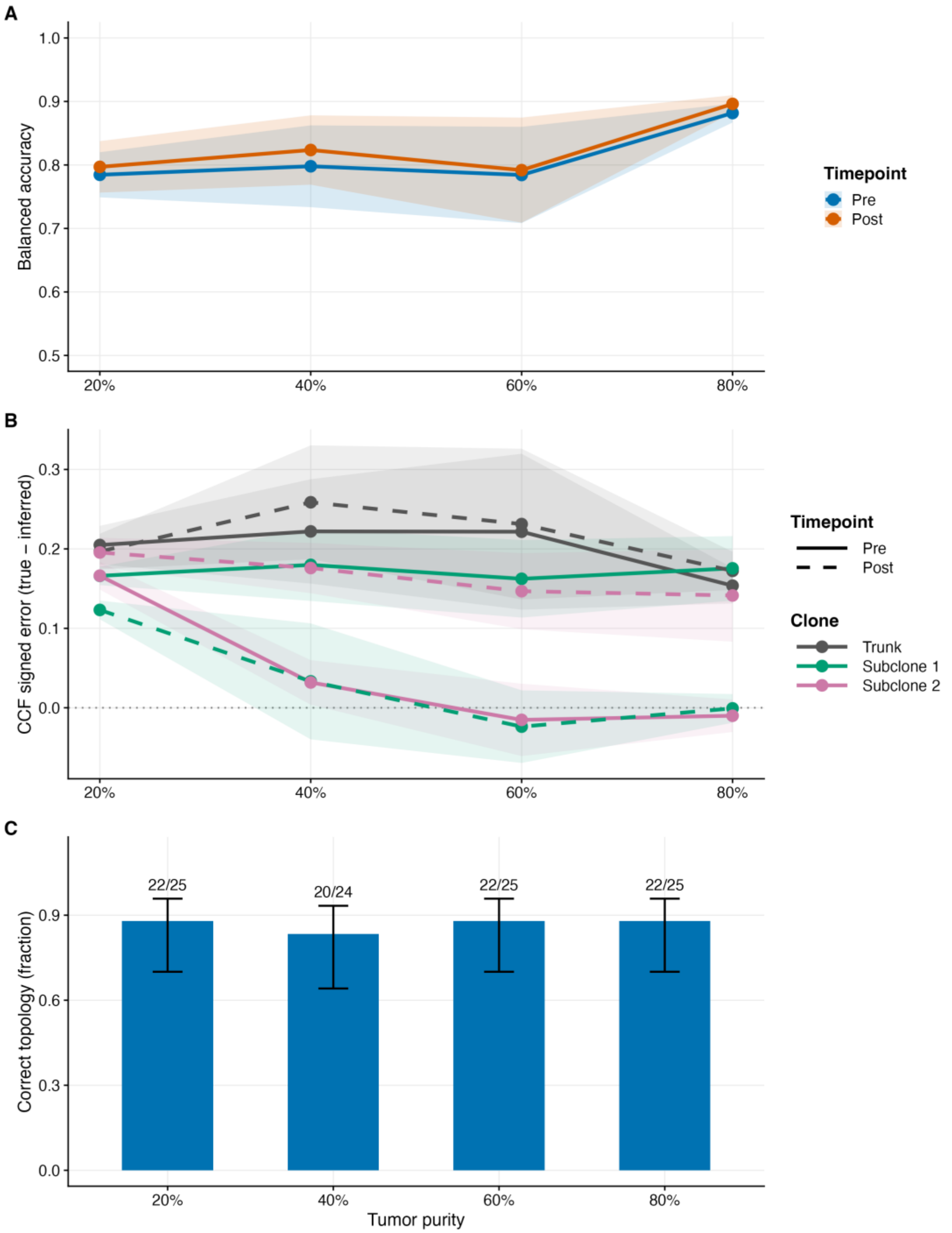
A
Balanced accuracy
Timepoint
Pre
Post
B
CCF signed error (true − inferred)
Timepoint
Pre
Post
Clone
Trunk
Subclone 1
Subclone 2
C
Correct topology (fraction)
22/25
20/24
22/25
22/25
20%
40%
60%
80%
Tumor purity

D

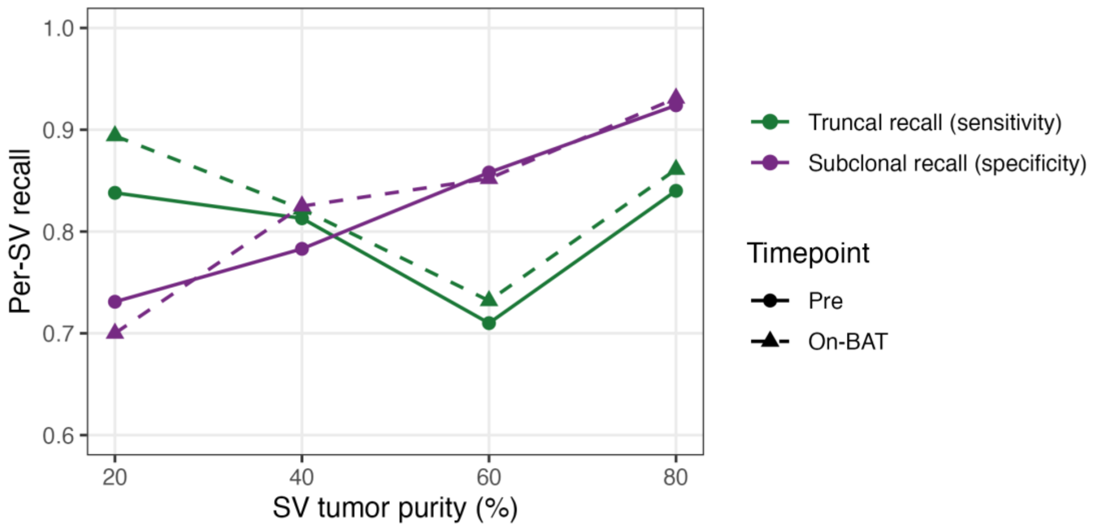


**Figure S8. Operating range of SVCFit's downstream phylogenetic inferences across the 20-80% SV tumor purity gradient.** S1 scenario, paired pre/post-treatment simulations, n = 25 paired replicates per purity, n = 24 at 40% (bootstraps 0-4 × the five overlap configurations).

(A) Per-SV truncal/subclonal balanced accuracy vs SV tumor purity, shown as two overlaid lines for pre-treatment (blue) and post-treatment (orange) timepoints; shaded ribbons indicate 95% CI.

(B) Per-clone signed CCF error (true − inferred) vs SV tumor purity, shown as three colored lines for the trunk (gray), Subclone 1 (green), and Subclone 2 (pink); pre-treatment is drawn as a solid line and post-treatment as dashed; shaded ribbons indicate 95% CI; positive values indicate underestimation.

(C) Bar plot of the fraction of replicates per purity in which the inferred rooted tree topology exactly matches ground truth; counts above each bar (e.g., 22/25) give the number of correct-topology replicates over the total replicates evaluated; error bars are bootstrap 95% CI.

(D) The two components whose average is the panel-A balanced accuracy, shown separately across the same purity gradient: per-SV truncal recall (sensitivity, green) and subclonal recall (specificity, purple), for pre-treatment (solid) and on-BAT (dashed) timepoints. The two trade off with purity, truncal recall higher at 20% purity and subclonal recall higher from 60% purity onward; this is the component-level breakdown, across all replicates, of the truncal/subclonal calling summarized as a single illustrative confusion matrix in Fig. S9D. Values are the sensitivity and specificity recovered from the same confusion-matrix evaluation used for panel A (no new simulation). Default SVCFit parameters were used in all replicates with no per-scenario tuning.

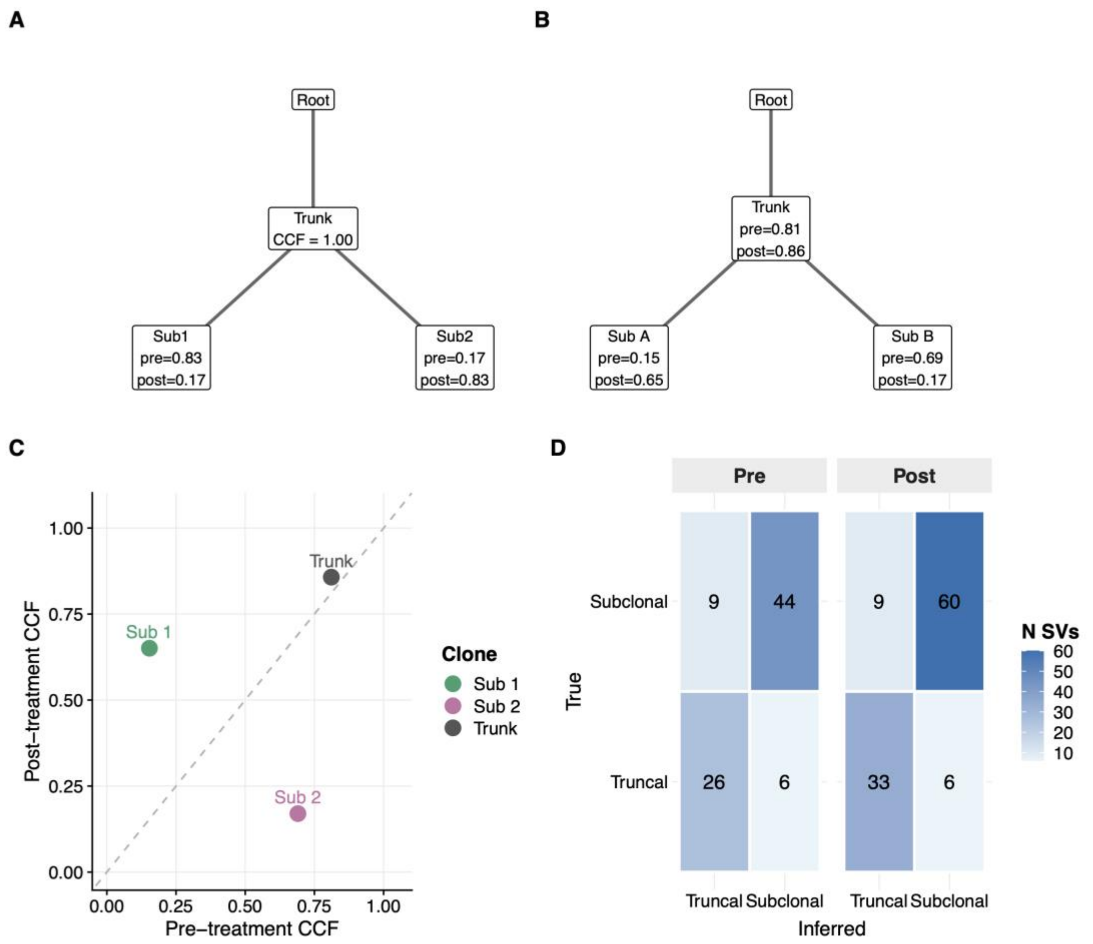


**Figure S9. Illustrative replicate from the S1 scenario at 60% tumor purity (no-overlapping-CNV configuration, bootstrap 0)**. (A) Ground-truth clonal tree. A truncal clone (CCF = 1.00 at both timepoints) gives rise to two subclones whose CCFs cross under treatment: Sub1 dominant pre-treatment (pre = 0.83, post = 0.17), Sub2 dominant post-treatment (pre = 0.17, post = 0.83). (B) Tree inferred by SVCFit and downstream clustering. The branching topology is recovered correctly. Consistent with the systematic positive signed error in Fig. S8B, per-clone CCF point estimates are pulled below the true values: the truncal clone is inferred at 0.81 (pre) / 0.86 (post) rather than 1.00, and the post-treatment dominant subclone is inferred at 0.65 rather than 0.83. (C) Paired pre/post CCFs of the inferred cluster centroids; the crossing-subclone pattern is preserved, with each centroid displaced toward the diagonal in proportion to the per-clone CCF underestimation visible in (B). (D) Confusion matrix of truncal vs. subclonal SV calls at the pre and post timepoints (counts of SVs per cell). Across both timepoints, balanced accuracy is approximately 0.85; the small number of misclassifications is dominated by truncal SVs called subclonal, the same direction of bias as in (B).

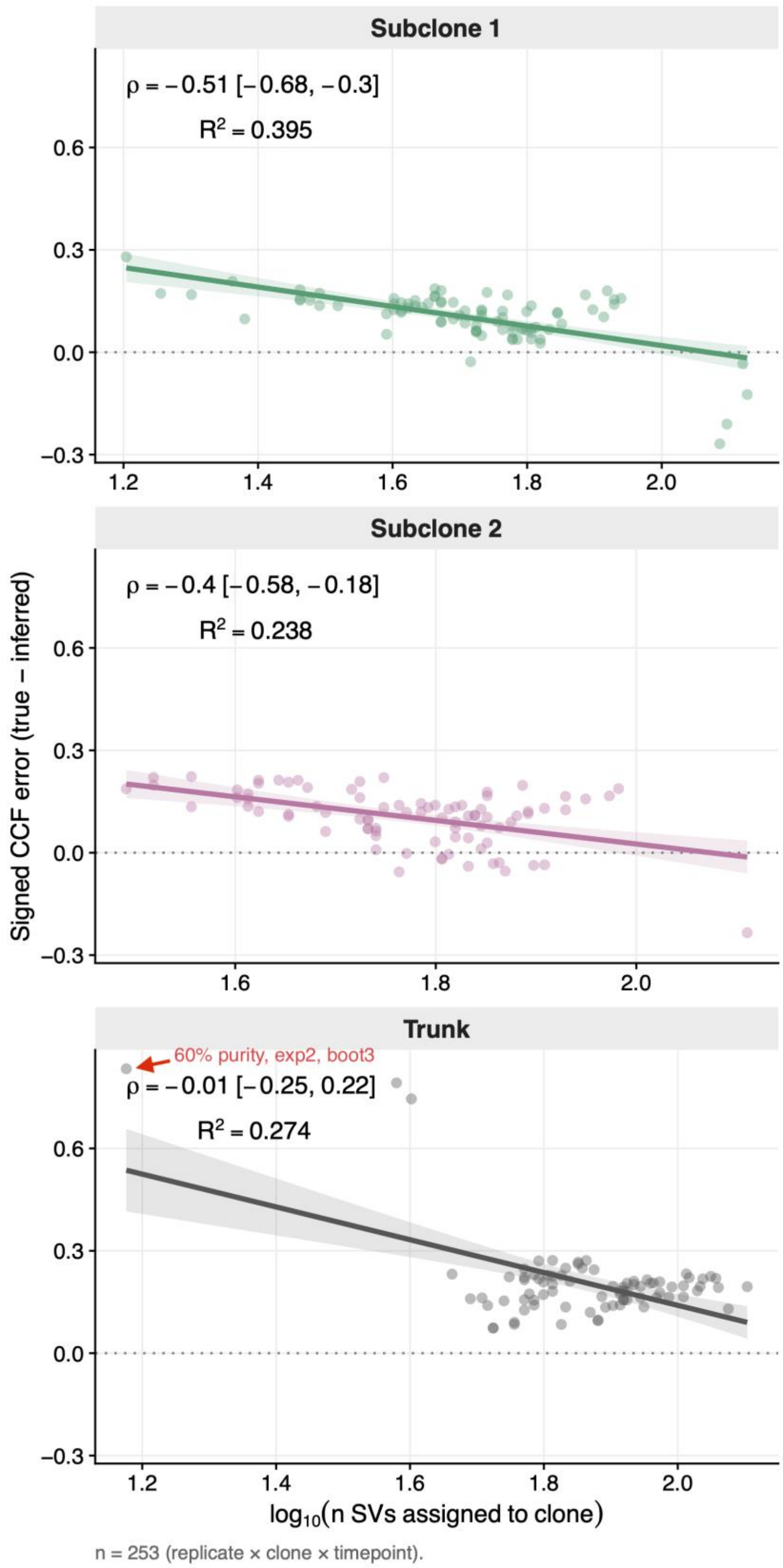
Subclone 1
ρ = −0.51 [−0.68, −0.3]
R² = 0.395
Subclone 2
ρ = −0.4 [−0.58, −0.18]
R² = 0.238
Trunk
60% purity, exp2, boot3
ρ = −0.01 [−0.25, 0.22]
R² = 0.274
Signed CCF error (true − inferred)
log10(n SVs assigned to clone)
n = 253 (replicate × clone × timepoint).

**Figure S10. Per-clone signed CCF error vs. number of SVs assigned to the cluster.** Three panels (top to bottom: Subclone 1, Subclone 2, Trunk) show signed CCF error (true − inferred CCF; positive = underestimation) as a function of $\log_{10}$(number of SVs assigned to the cluster). Each point is one (replicate × clone) pair (signed error averaged over the two timepoints), drawn from the bootstraps 0-4 × five-configuration simulation series at SV tumor purities 20%, 40%, 60%, 80% with default SVCFit parameters (n = 253 total). Cluster size is computed as the pair-total count (combining pre- and post-treatment timepoints), matching SVCFit's `enforce_min_cluster_size(min_n = 5)` invariant. In-panel annotations report Spearman ρ with bootstrap 95% CI and the linear-fit $R^2$. Subclonal CCF error decreases as cluster size grows (Subclone 1 ρ = −0.51 [−0.68, −0.30]; Subclone 2 ρ = −0.40 [−0.58, −0.18]), consistent with the expected sampling-noise effect that more SVs per cluster support a tighter cluster-level CCF estimate. The truncal cluster shows no monotone relationship with cluster size (ρ = −0.01 [−0.25, +0.22]); the higher linear-fit $R^2$ of the truncal panel (0.274) is driven by a single high-leverage replicate (a 60% purity bootstrap; $\log_{10}$(n) =1.17, signed error = 0.8) in which the DP-GMM split the truncal SVs into multiple components, producing a small truncal cluster with an unusually large CCF residual; the rank-based Spearman ρ is the robust summary statistic for the per-cluster size effect.

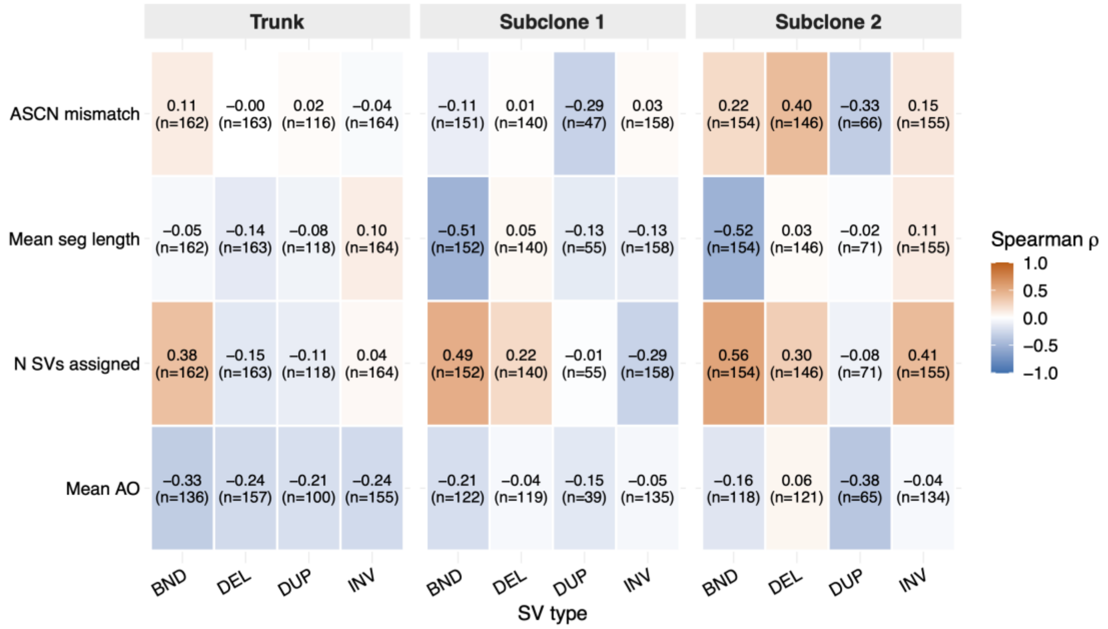


**Figure S11. SV-type-stratified covariate analysis of per-clone signed CCF error.** Heatmap of Spearman ρ between four candidate cluster-level covariates (rows, top to bottom: ACR mismatch fraction; mean ACR segment length; $\log_{10}$ of the number of SVs assigned to the cluster; mean SVtyper alternate-allele read support, AO) and the per-clone signed CCF error (true − inferred CCF). Columns are organized as three clone-group blocks (Trunk; Subclone 1; Subclone 2), each subdivided into four SV-type sub-columns (BND, DEL, DUP, INV; no INS-class SVs were generated in the simulation design). Each cell reports ρ with the number of contributing observations. Three patterns localize the SV-type contribution to per-clone CCF residuals. (i) Cluster-size × BND is positive in both subclones (ρ = +0.49 in Subclone 1; ρ = +0.56 in Subclone 2), in apparent contrast to the aggregate negative cluster-size correlation in Fig. S10, indicating that the aggregate sampling-noise effect (more SVs in a cluster supporting a tighter centroid estimate) masks a smaller BND-specific contribution in which BND-rich clusters carry larger errors. (ii) Mean-segment-length × BND is strongly negative in the subclones (ρ = −0.51 and −0.52), consistent with shorter BNDs carrying lower mappability. (iii) Mean-AO × BND is negative across all clones (most strongly in the trunk, ρ = −0.33), so clusters with lower mean AO carry larger residuals, consistent with noisier alternate-allele counts at BND breakpoints. Non-BND SV types show weaker or absent correlations across the four covariates, consistent with mappability-driven miscounting being specific to breakend-resolved SVs.

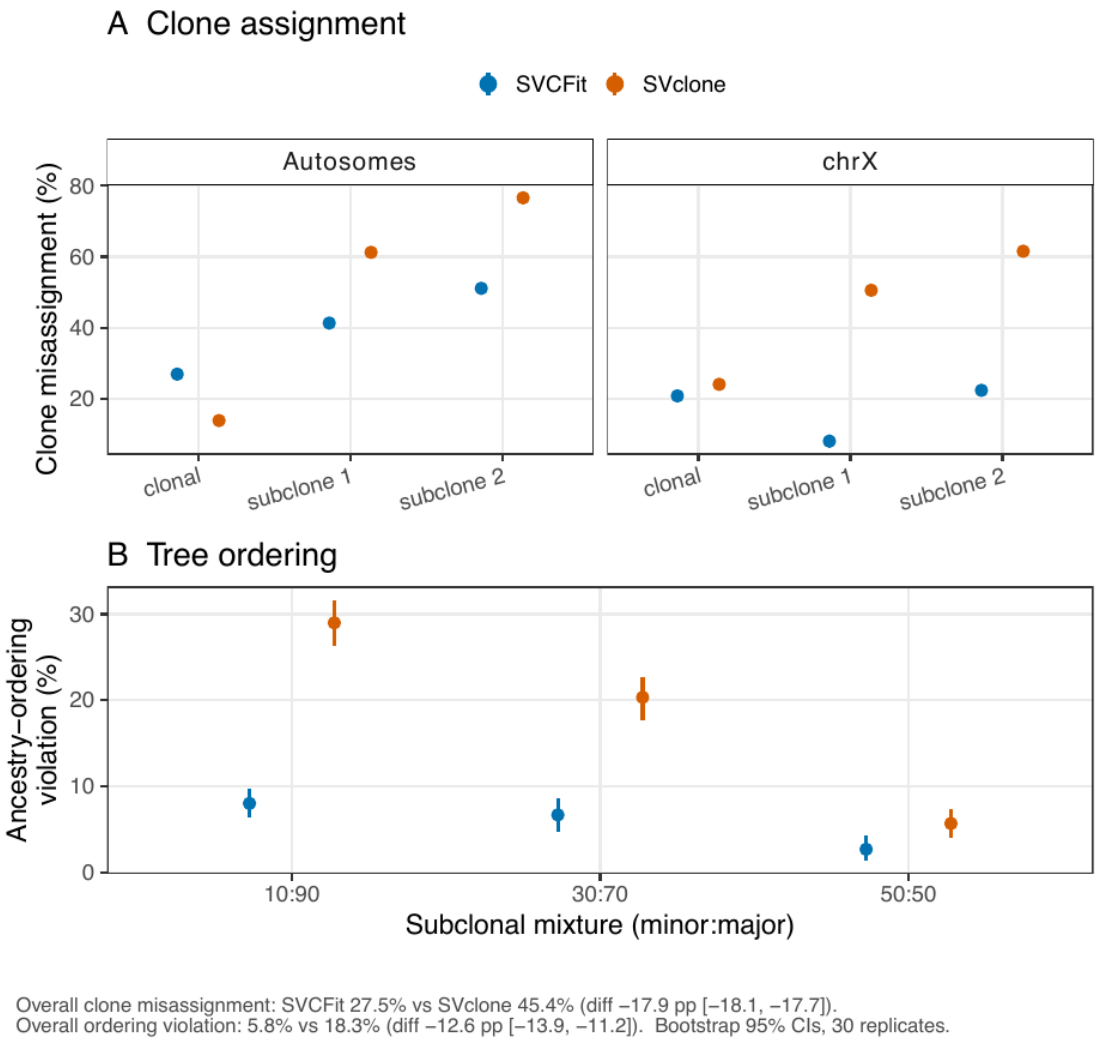


**Figure S12.** Downstream decision-boundary consequences of SVCF error: SVCFit versus SVclone (autosomes + chrX). Practical impact of CCF-estimation error, scored against the known simulation truth head-to-head, where an error matters only if it crosses a discrete downstream boundary. (A) Clone assignment: each SV is assigned to the clone whose true CCF level is nearest (oracle centroids, isolating the CCF-error effect from clustering-estimation error); a misassignment is when the nearest level is not the SV's own clone. Point ranges are the misassignment rate with bootstrap 95% CI (30 replicates), by clone and chromosome source. Overall SVCFit 27.5% versus SVclone 45.4% (Δ = −17.9 percentage points [−18.1, −17.7]); the advantage is larger on chrX (−25.8 points) than on the autosomes (−8.2 points), owing to SVCFit's hemizygous handling, and is concentrated in the fragile subclonal decisions. (B) Tree ordering: because the simulated topology is clonal to {subclone 1, subclone 2}, a subclone whose estimated mean CCF exceeds the clonal CCF would be inferred as ancestral to the truncal clone, an impossible topology. Ordering-violation rate with bootstrap 95% CI by subclonal mixture: overall SVCFit 5.8% versus SVclone 18.3% (Δ = −12.6 points [−13.9, −11.2]), worst at the tightest (10:90) margin. This is the head-to-head, chrX-inclusive complement to the SVCFit-only, autosome-only phylogeny benchmark (Supplementary Figs. S8-S11), which is kept separate. Caveat: the oracle centroids isolate the CCF-error effect, so absolute rates are high; the robust signal is the between-method difference, scored identically for both tools. Per-metric statistics are in Supplementary Table S12.

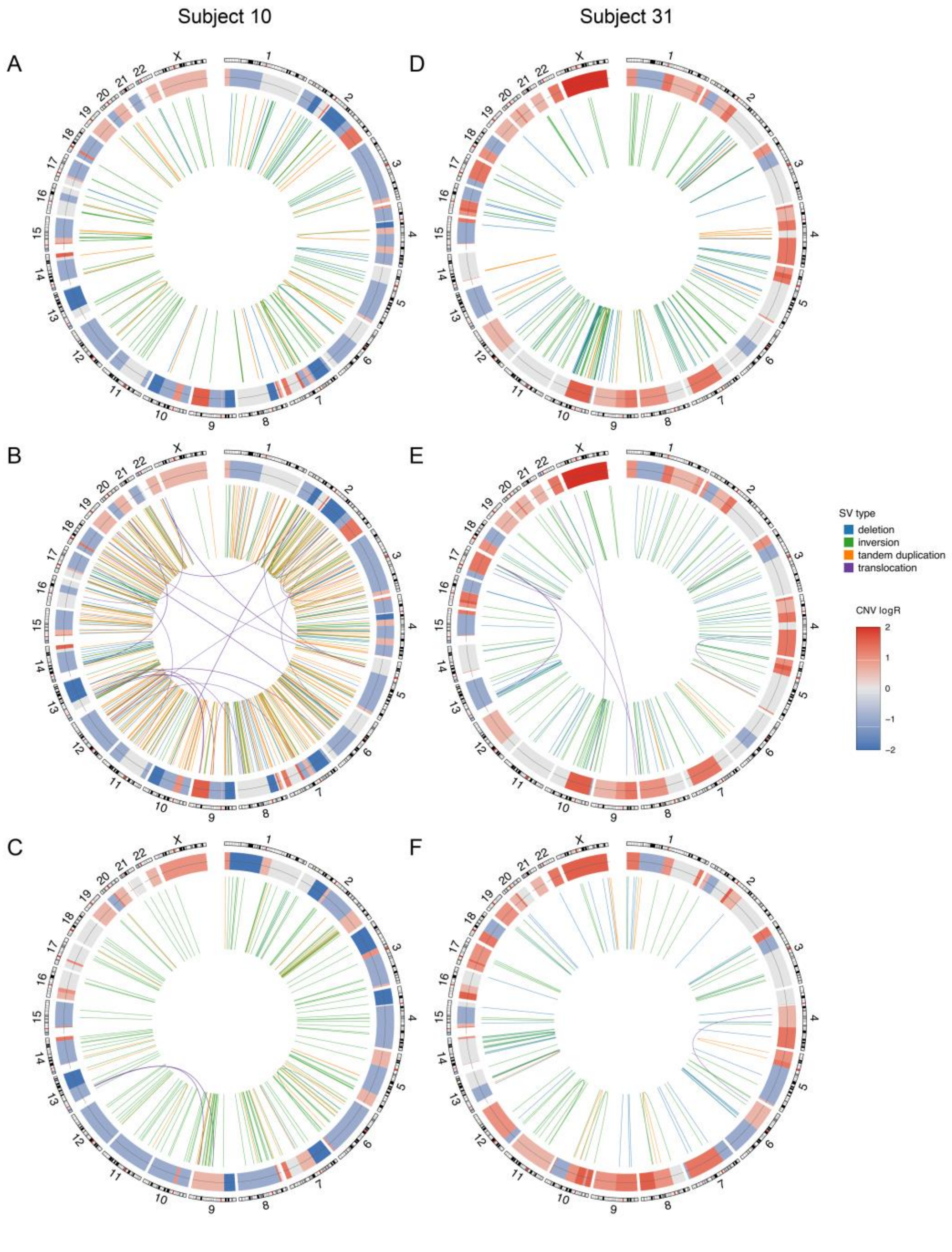


**Figure S13. Genome-wide structural variation in pre- and on-BAT samples.** Subject 10 (A-C); Subject 31 (D-F). (A,D) Present at both time points; (B,E) pre-BAT only; (C,F) on-BAT only. The outermost track is the chromosomal ideogram (chromosomes 1-22 and X). The inner heatmap track shows somatic copy number (red=gain, blue=loss). The central links show SVs: deletions (blue), inversions (green), tandem duplications (orange), and translocations (purple).

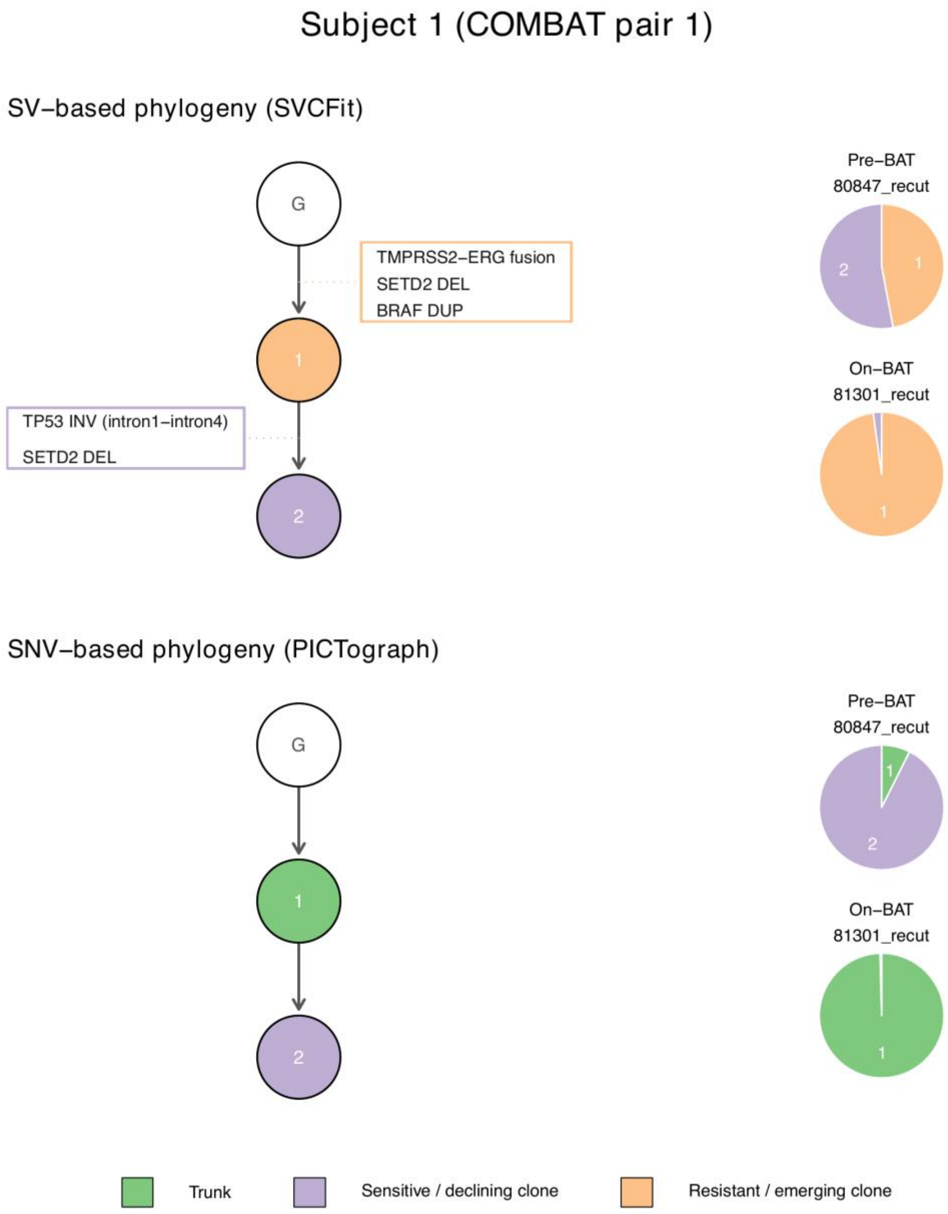


**Figure S14-1.** Subject 1 (COMBAT pair 1): SV-based phylogeny (SVCFit, top) and SNV-based phylogeny (PICTograph, bottom).

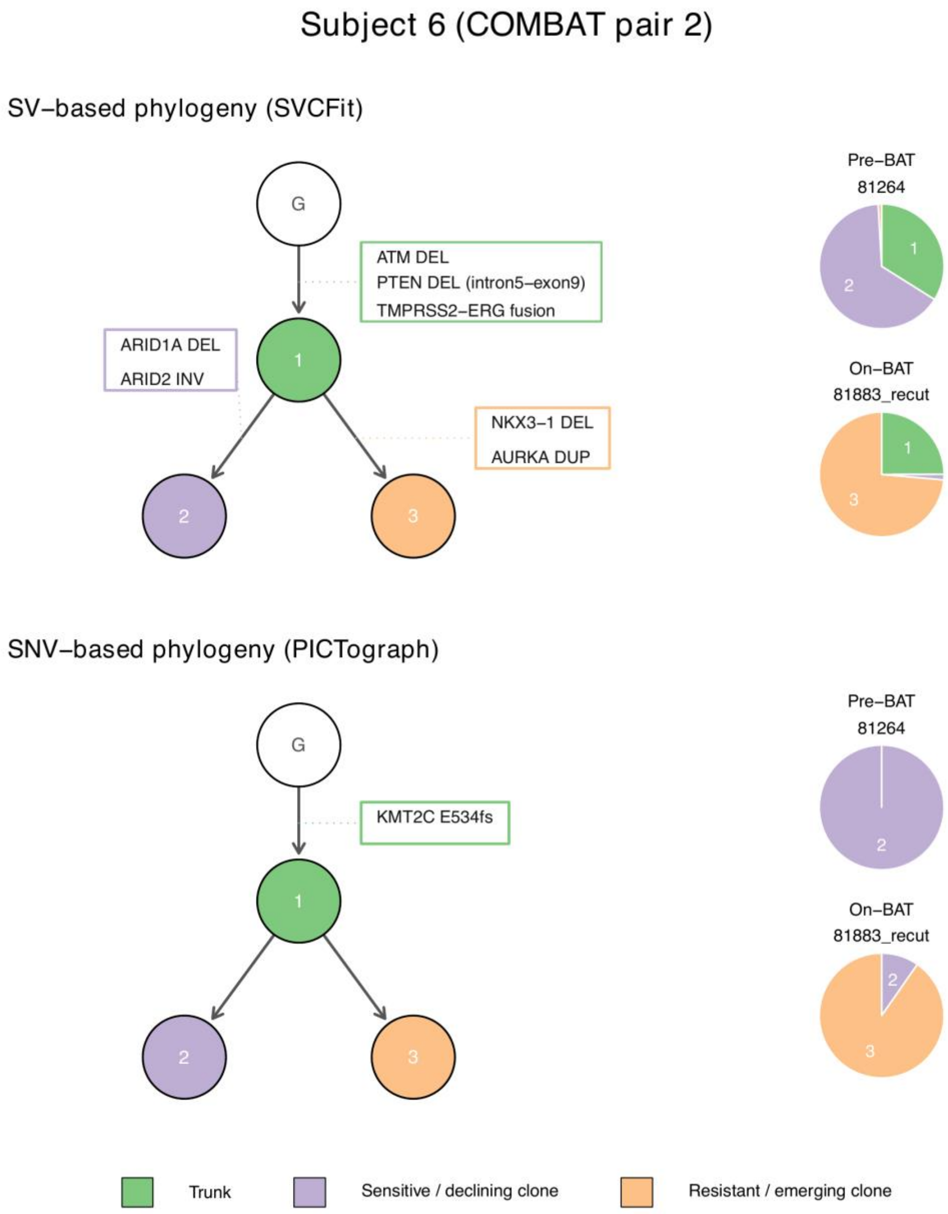


**Figure S14-2.** Subject 6 (COMBAT pair 2): SV-based phylogeny (SVCFit, top) and SNV-based phylogeny (PICTograph, bottom).

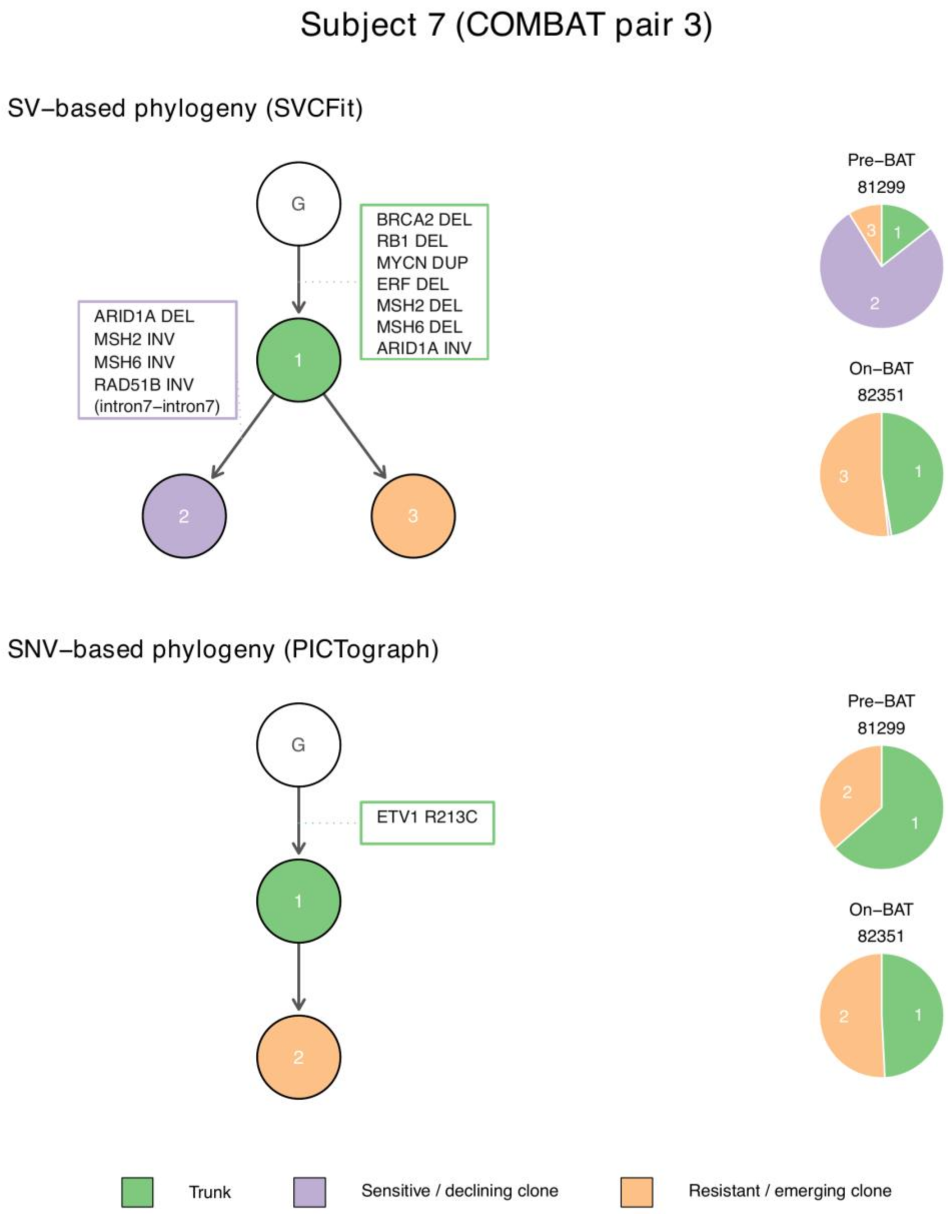


**Figure S14-3.** Subject 7 (COMBAT pair 3): SV-based phylogeny (SVCFit, top) and SNV-based phylogeny (PICTograph, bottom).

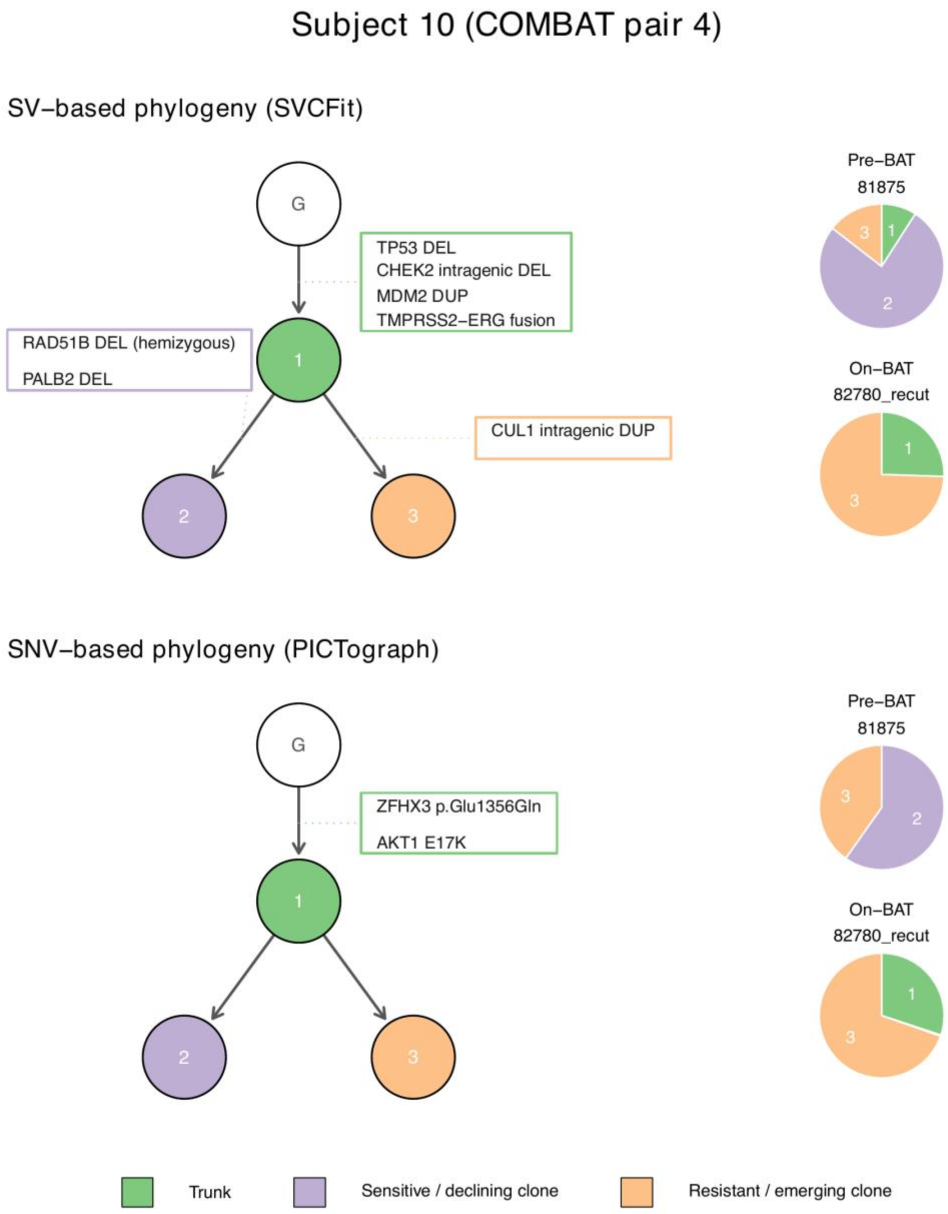


**Figure S14-4.** Subject 10 (COMBAT pair 4): SV-based phylogeny (SVCFit, top) and SNV-based phylogeny (PICTograph, bottom).

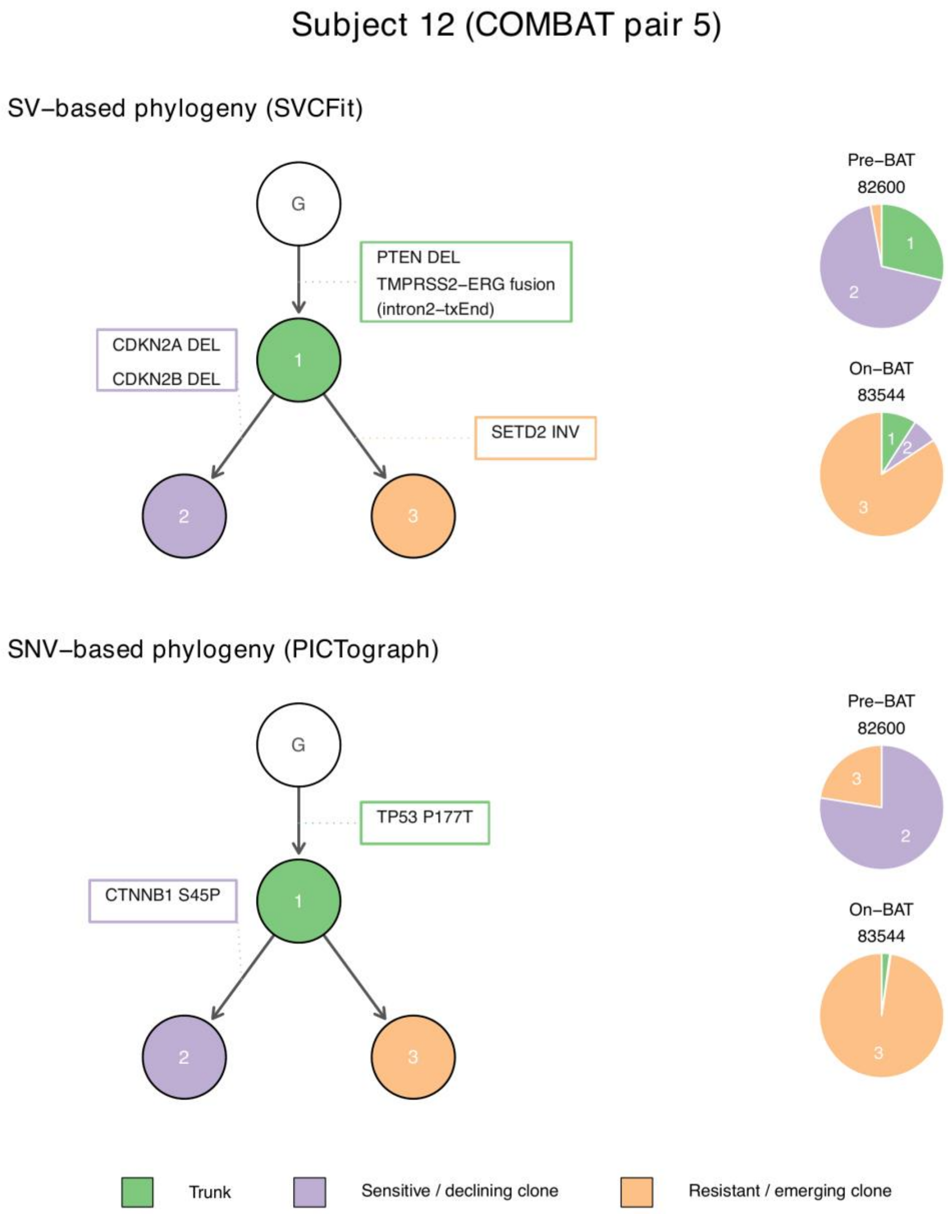


**Figure S14-5.** Subject 12 (COMBAT pair 5): SV-based phylogeny (SVCFit, top) and SNV-based phylogeny (PICTograph, bottom).

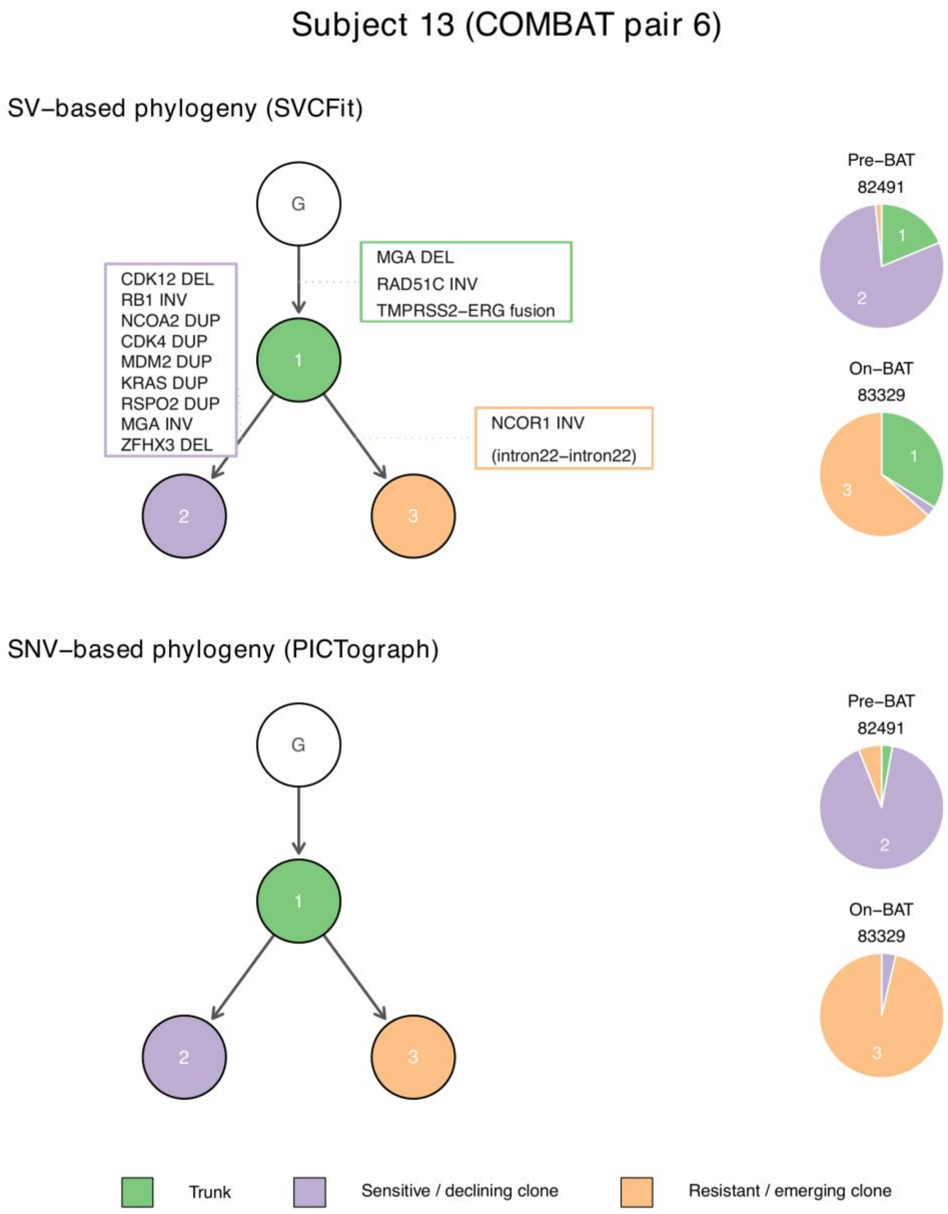


**Figure S14-6.** Subject 13 (COMBAT pair 6): SV-based phylogeny (SVCFit, top) and SNV-based phylogeny (PICTograph, bottom).

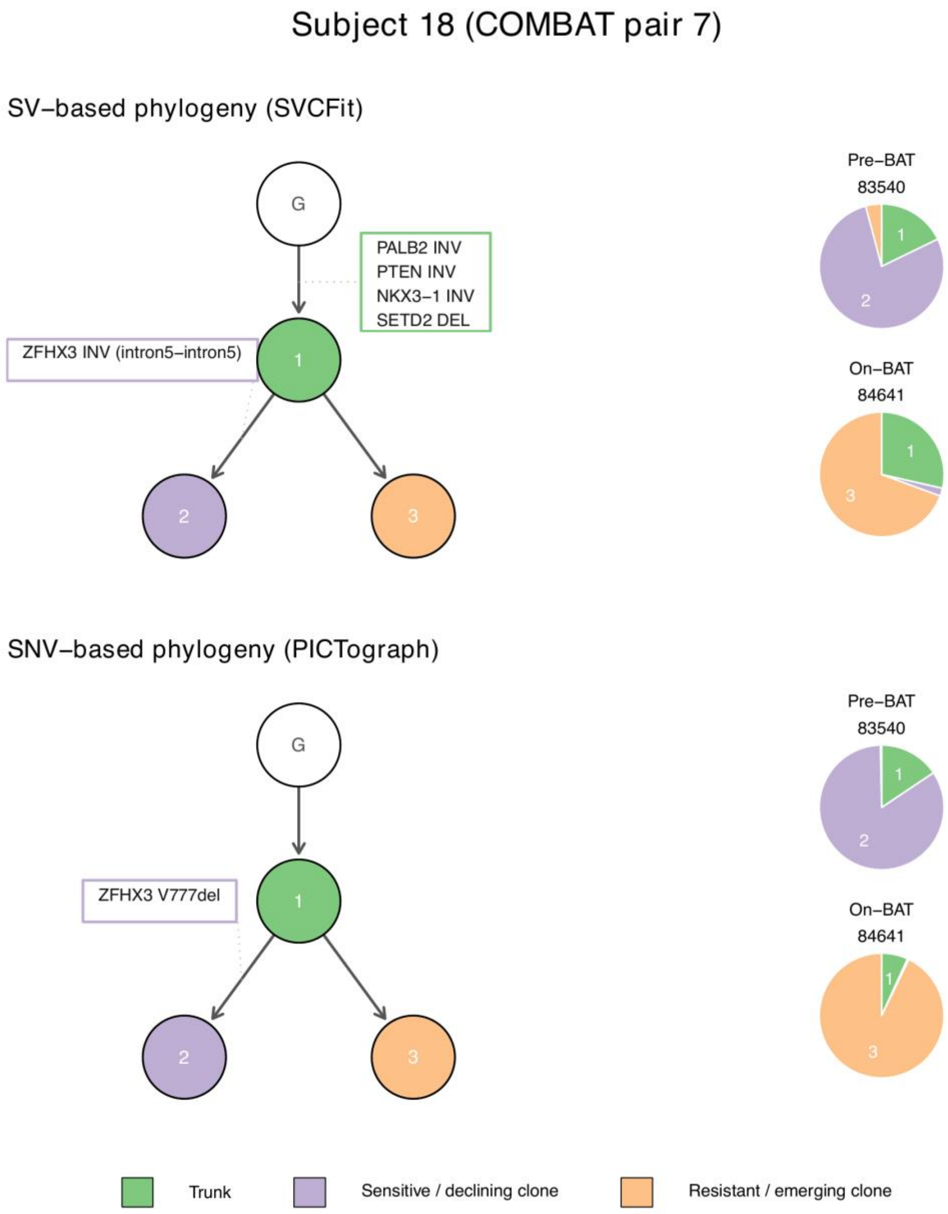


**Figure S14-7.** Subject 18 (COMBAT pair 7): SV-based phylogeny (SVCFit, top) and SNV-based phylogeny (PICTograph, bottom).

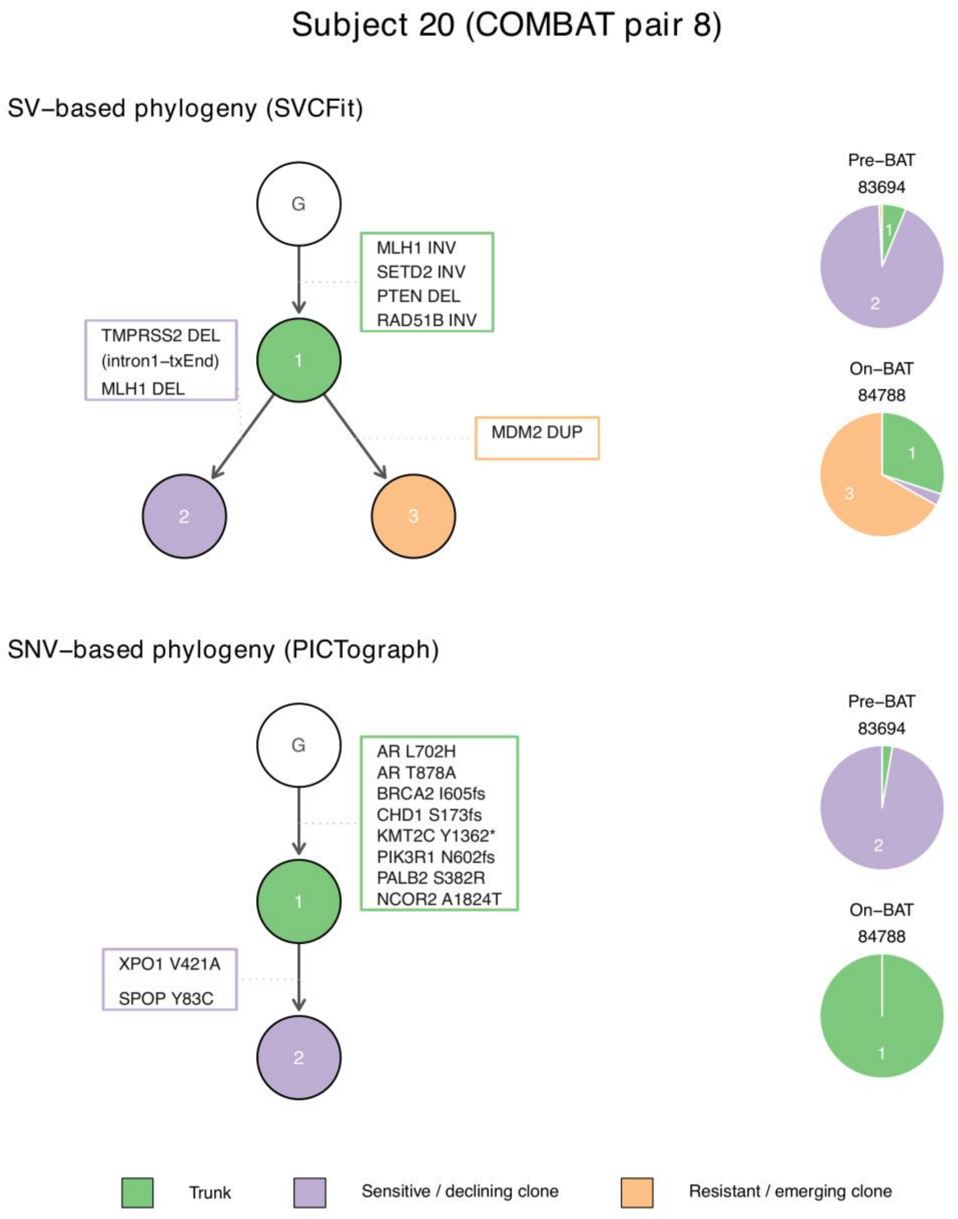


**Figure S14-8.** Subject 20 (COMBAT pair 8): SV-based phylogeny (SVCFit, top) and SNV-based phylogeny (PICTograph, bottom).

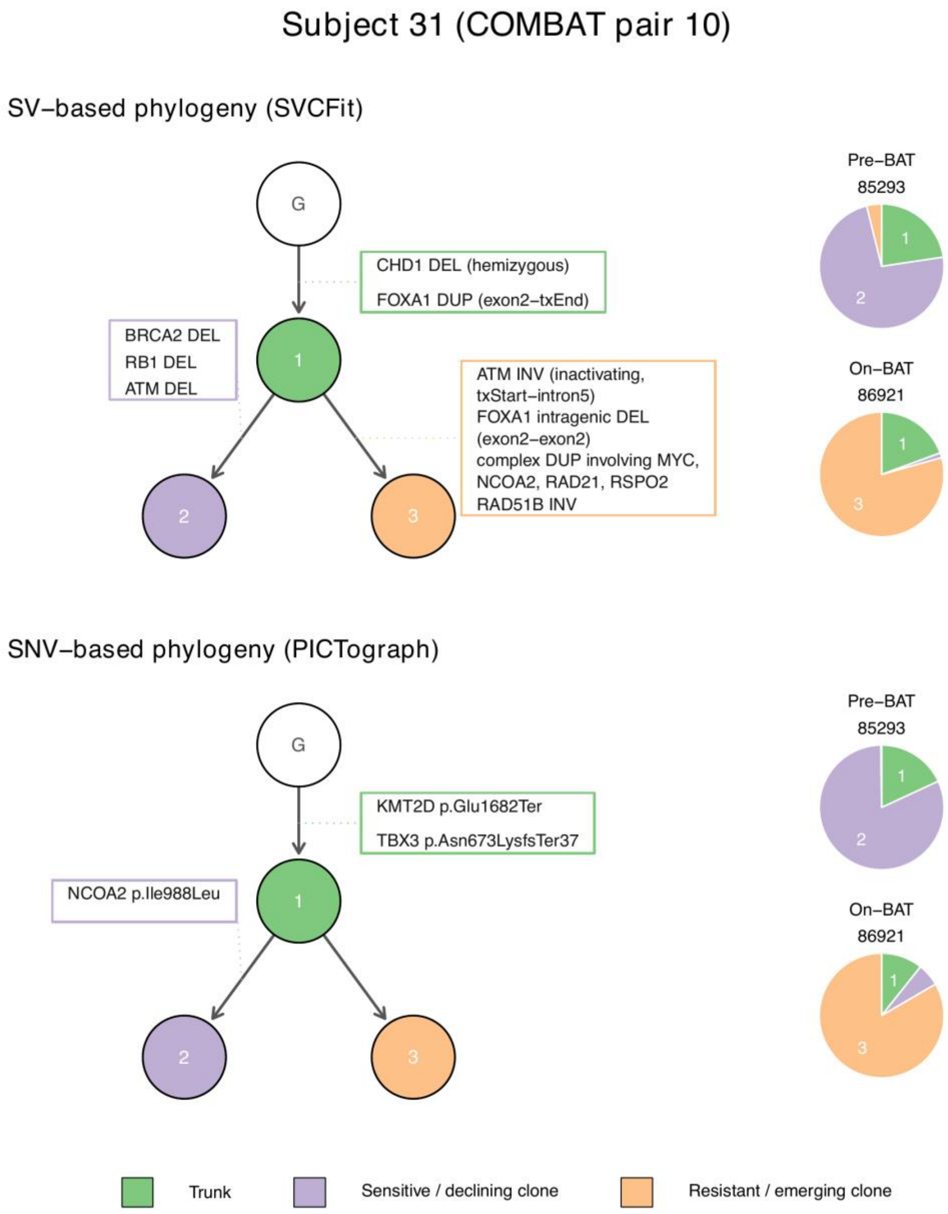


**Figure S14-9.** Subject 31 (COMBAT pair 10): SV-based phylogeny (SVCFit, top) and SNV-based phylogeny (PICTograph, bottom).

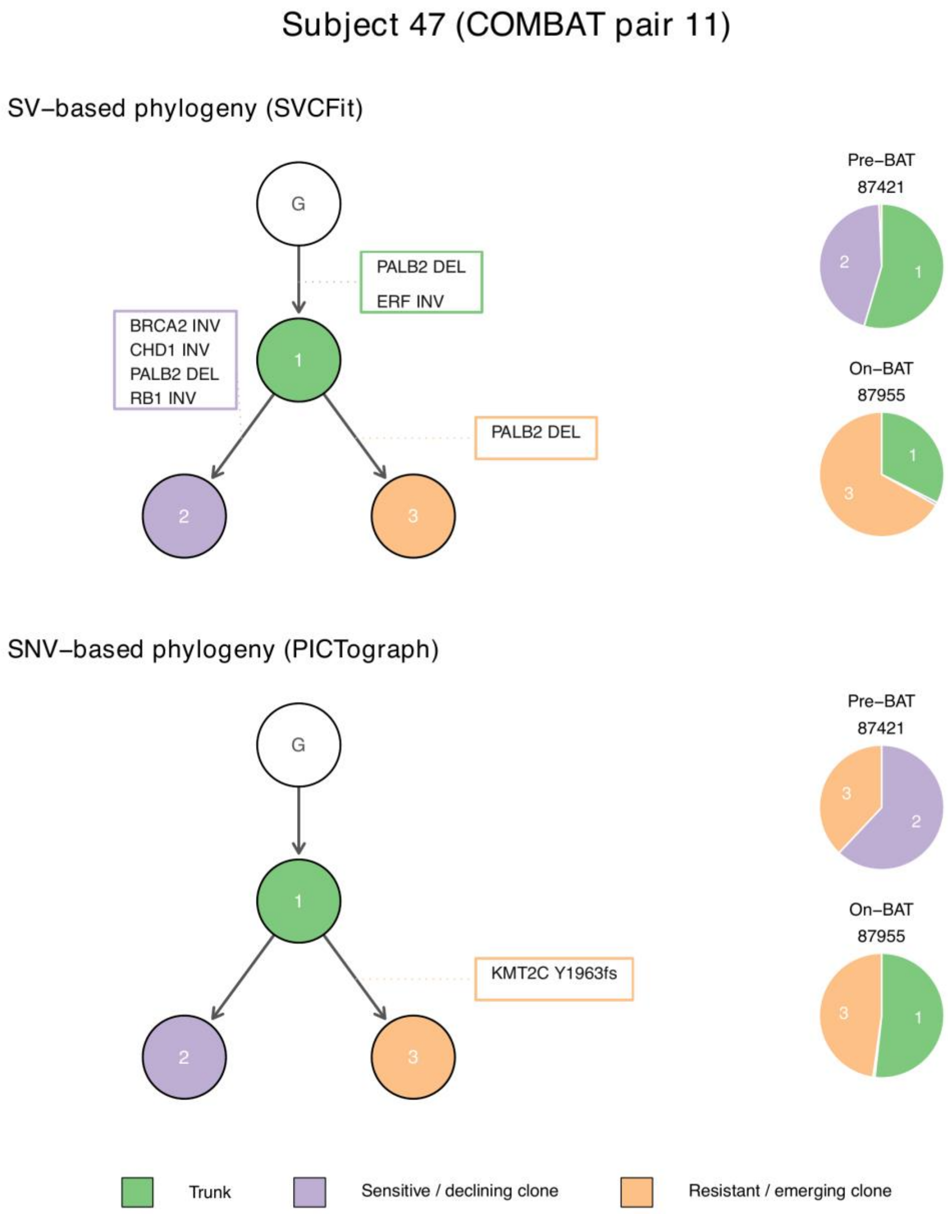


**Figure S14-10.** Subject 47 (COMBAT pair 11): SV-based phylogeny (SVCFit, top) and SNV-based phylogeny (PICTograph, bottom).

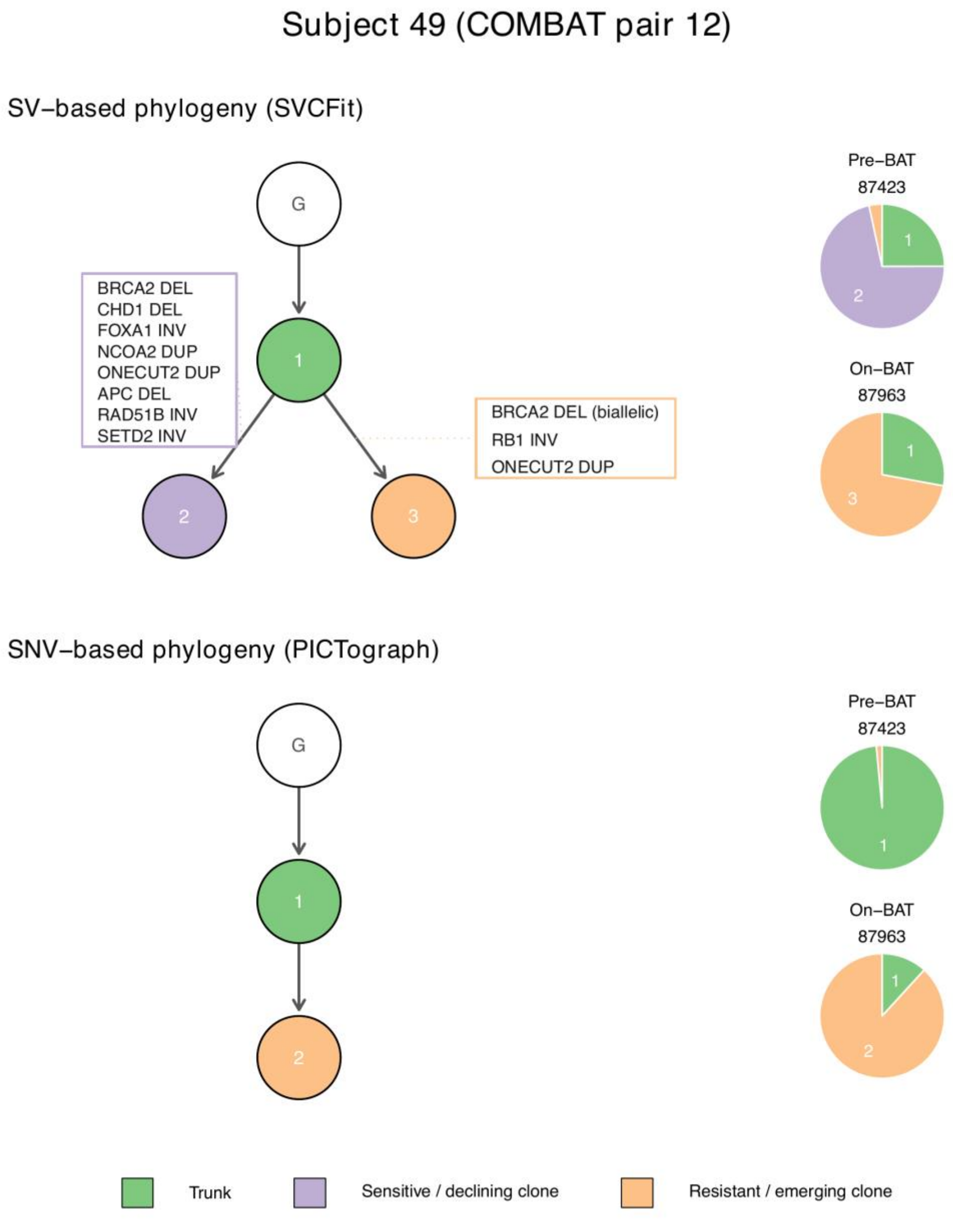


**Figure S14-11.** Subject 49 (COMBAT pair 12): SV-based phylogeny (SVCFit, top) and SNV-based phylogeny (PICTograph, bottom).

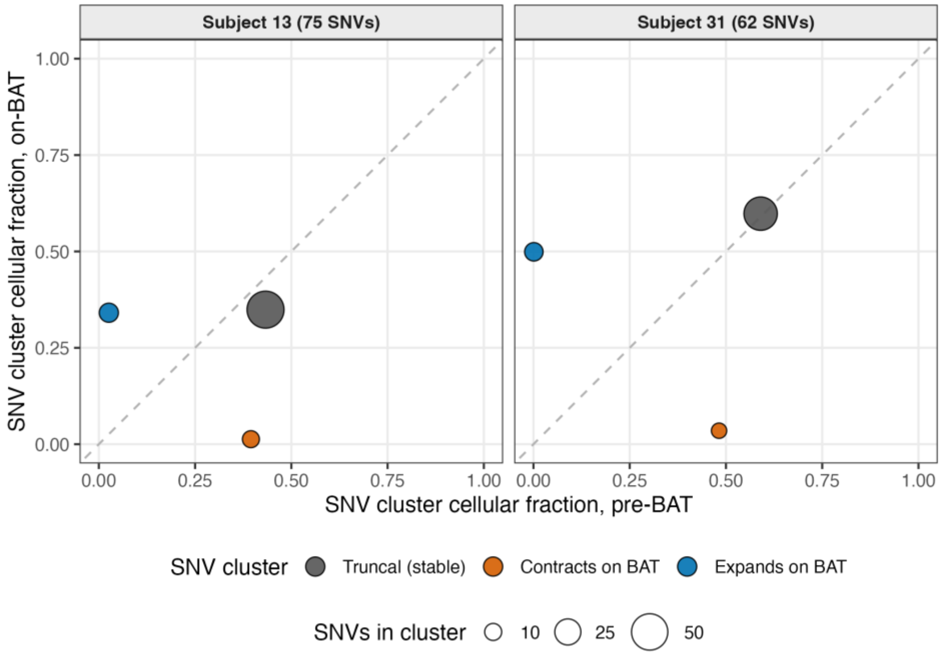


**Figure S15.** SNV cluster cellular fraction across the paired pre- and on-BAT samples for Subjects 13 and 31 (co-clustering). PICTograph estimates cellular fraction per SNV cluster rather than per individual SNV, so each point is one cluster, positioned at its estimated cellular fraction in the pre-BAT sample (x-axis) and the on-BAT sample (y-axis), with point area proportional to the number of SNVs assigned to the cluster (Subject 13: 75 SNVs in three clusters; Subject 31: 62 SNVs in three clusters). The dashed line marks equal pre- and on-BAT fraction. PICTograph performed multi-sample reconstruction, using both timepoints jointly. Both subjects resolve into three clusters rather than a single (monoclonal) population: a stable truncal cluster present at both timepoints (grey), a cluster that contracts on BAT (orange), and a cluster that expands on BAT (blue), recapitulating the clonal-replacement dynamics inferred from the SV-based trees.

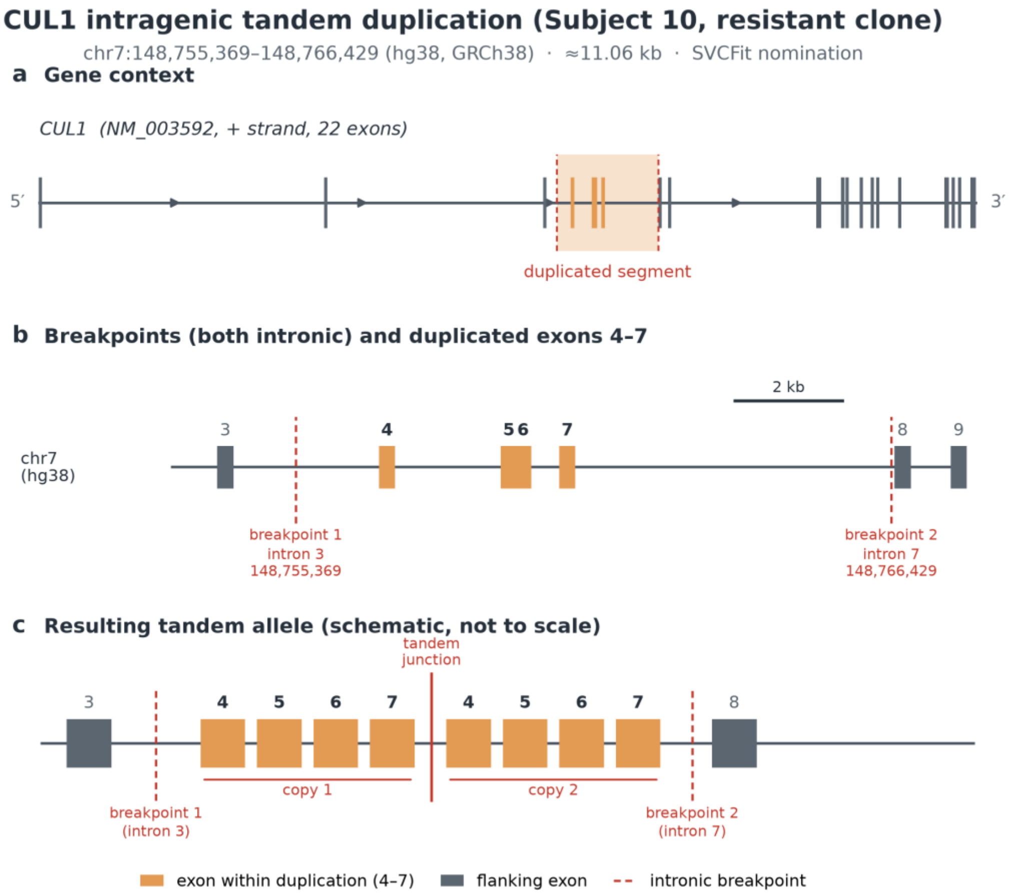


**Figure S16.** CUL1 intragenic tandem duplication nominated by SVCFit in the resistant clone (Clone 3) of Subject 10. (a) CUL1 gene model (RefSeq NM_003592; chr7, plus strand; 22 exons; GRCh38/hg38, UCSC RefGene) with the duplicated interval chr7:148,755,369-148,766,429 (approximately 11.06 kb) shaded; exons within the interval are colored. (b) Genomic zoom (drawn to scale; 2 kb bar) spanning exons 3-9. Both duplication breakpoints lie in introns: breakpoint 1 (148,755,369) in intron 3 and breakpoint 2 (148,766,429) in intron 7. The duplicated segment therefore has intronic breakpoints but encompasses coding exons 4-7; it is intragenic rather than purely intronic, and "intragenic duplication" in the main text refers to this configuration. (c) Schematic of the resulting tandem allele (not to scale), in which exons 4-7 are present in two head-to-tail copies separated by the tandem junction. The event is defined from SVCFit breakpoint evidence and is reported as a candidate resistance-associated structural alteration; its transcriptional consequence was not experimentally assessed and no regulatory-element annotation is asserted.

#### Supplementary Tables

**Table S1. Bioinformatics tools, versions, and parameters used in the analysis.**

| Step | Tool | Ver. | Key parameters | Inputs | Outputs | Where used |
|---|---|---|---|---|---|---|
| Genome simulation | VISOR | v1.1.3 | Coverage, purity, clonal proportion | BED | BAM | VISOR benchmark |
| Read trimming | trim_galore | v0.6.1 | Default | FASTQ | Trimmed FASTQ | COMBAT |
| Alignment | bwa mem | v0.7.19 | Defaults | FASTQ, Refs^a | BAM | COMBAT; Prostate mix |
| Mark duplicates | GATK4 | v4.6.2 | Defaults | BAM | Processed BAM | COMBAT |
| Germline SNP | GATK4 HC | v4.6.2 | Defaults | Normal BAM | Germline VCF | All datasets |
| Het SNP filter | bcftools | v1.20 | `-i 'GT="0/1"'` | Germline VCF | Het-SNP VCF | All datasets |
| Somatic SV | Manta | v1.6.0 | Default | Tumor & Normal BAM | VCF | Prostate; COMBAT |
| Somatic SV | Delly | v1.5.0 | Defaults | Tumor & Normal BAM | VCF | COMBAT |
| Somatic SV | GRIDSS | v2.13.2 | Defaults | Tumor & Normal BAM | VCF | COMBAT |
| SV merge | SURVIVOR | v1.0.7 | Merge multi-caller | VCFs | Consensus VCF | COMBAT |
| SV genotyping | SVtyper | v0.7.1 | Lumpy fmt; added CI^b | SV VCF, BAMs | Genotyped VCF | All datasets |
| CNV calling | FACETS | v0.6.2 | `-g -q15 -Q20 -P100 -r25,0` | Tumor & Normal BAM | CNV profile | All datasets |
| BP read filter | samtools | v1.21 | `-f 1 -F 2` | Tumor BAM | Filtered BAM | All datasets |

| Step | Tool | Ver. | Key parameters | Inputs | Outputs | Where used |
|---|---|---|---|---|---|---|
| SNP pileup | bcftools | v1.20 | mpileup on filtered BAM | BAM, SNP VCF | SNP pileup | All datasets |

Notes: ^a References included GRCh38 and hs37d5. ^b Added CIPOS and CIEND tags. Abbreviations: HC (HaplotypeCaller), BP (Breakpoint), CI (Confidence Interval).

**Table S2. Citations supporting driver-gene status for genes annotated on the COMBAT cohort SV phylogenies.**

This table compiles literature support for the driver-gene status of every gene annotated on the SV-based or SNV-based phylogenies in the main-text Results, the Subject 10 and 31 detailed analyses (Fig. 5 and Methods "Rescue of a key driver event"), and the Supplementary Note S4 per-subject tables. Genes are organized by pathway. Hosseini et al. (2025)[37] is the source of the prostate-cancer driver gene list used for tree edge annotation (Methods, "SV Tumor Evolution Tree Construction and Annotation"); inclusion of a gene in that list constitutes the methodological basis for its annotation on the trees. Additional pathway- or context-specific references are listed where they exist in the bibliography. Three gene-specific functional-resistance claims (BRAF duplication on the resistant clone of Subject 1; *RB1* loss in Subjects 7 and 49; *MDM2 duplication on the resistant clone of Subject 20)* carry citation clusters in their respective per-gene rows below. The *RB1* row Notes column clarifies that *RB1* loss is HR-supporting but not equivalent to *BRCA1/2* loss and distinguishes the Subject 7 (possible 13q co-deletion) from the Subject 49 (independent inversion) configurations.

| Gene | Pathway / role | Subject(s) | reference(s) | Notes |
|---|---|---|---|---|
| ***AR*** | Androgen receptor; ligand-binding domain | 20 (truncal L702H and T878A SNVs, recovered from filtered calls) | Hosseini 2025[33] | Ligand-binding-domain mutations of the type selected by AR-directed therapy, which this subject had received before enrollment. Both are recorded as somatic prostate-cancer variants in ClinVar and as recurrent prostate hotspots in |

| Gene | Pathway / role | Subject(s) | reference(s) | Notes |
|---|---|---|---|---|
| | | | | COSMIC. Both were removed by the variant caller's germline-risk filter and were recovered as described in Methods, "Rescue of key driver events". Truncal, so present in every tumor cell at both timepoints, marking the founder genotype rather than a BAT-resistant branch. |
| ***TMPRSS2-ERG (fusion)*** | AR-axis founder fusion | 1 (truncal, IHC-confirmed), 6 (truncal, IHC-confirmed), 10 (truncal), 12 (truncal, IHC-confirmed ERG expression in both timepoints), 13 (truncal, IHC-confirmed) | Tomlins 2005[29], Weier 2013[30], TCGA 2015[31], Yoshimoto 2008[42] | Canonical early prostate-cancer fusion |
| ***ERG*** (rearrangement / IHC marker) | AR-axis | 10 (IHC pre-/on-BAT), 20 (subclonal fs) | Tomlins 2005[29], Morais 2015[32] | IHC interpreted as proxy for *ERG* rearrangement |

| Gene | Pathway / role | Subject(s) | reference(s) | Notes |
|---|---|---|---|---|
| ***FOXA1*** | AR cistrome | 31 (truncal focal dup + resistant intragenic del), 49 (sensitive inv) | Hosseini 2025[33], Quigley 2018[34], Viswanathan 2018[35] | mCRPC AR-cistrome remodeling |
| ***NCOA2*** | AR coactivator | 13 (sensitive dup), 31 (resistant dup), 49 (sensitive dup) | Hosseini 2025[33], Quigley 2018[34] | Recurrent duplication in mCRPC |
| ***NCOR1*** | AR co-repressor | 13 (resistant inv) | Hosseini 2025[33], Lopez 2016[14] (NCOR1 expression and output declines with prostate cancer progression; loss of NCOR function results in enhanced AR signaling and is found in ~16% of metastatic prostate tumors; predicts castration-therapy resistance) | Sole resistant-clone branch-specific event in Subject 13; both breakpoints within intron 22; protein-coding disruption cannot be inferred from SV breakpoint evidence; candidate *NCOR1* locus alteration but functional significance is uncertain and requires transcriptomic validation |
| ***SPOP*** | E3-ligase substrate adaptor; AR turnover | 20 (sensitive Y83C SNV) | TCGA 2015[31], Quigley 2018[34] | Canonical primary prostate-cancer driver |
| ***TP53*** | Tumor suppressor | 10 (truncal del), 12 | Hosseini 2025[33], Quigley 2018[34], | Canonical mCRPC truncal driver |

| Gene | Pathway / role | Subject(s) | reference(s) | Notes |
|---|---|---|---|---|
| | | (truncal P177T) | Viswanathan 2018[35] | |
| ***RB1*** | Tumor suppressor; cell cycle; HR-supporting via CtIP-dependent end resection and BRG1 recruitment | 7 (truncal del → resistant in HRD context), 13 (sensitive inv), 31 (sensitive hem del), 47 (sensitive inv), 49 (resistant inv) | Sondka 2018[7] (canonical Tier 1 tumor suppressor), Hamid 2019[12] and Nyquist 2020[13] (*RB1* loss in localized and metastatic prostate cancer; *TP53* + *RB1* compound loss confers resistance to a spectrum of therapeutics), Vélez-Cruz 2016[11] (RB localizes to DNA double-strand breaks and promotes DNA end resection and homologous recombination through recruitment of BRG1), Chakraborty 2021[22] (*BRCA2/RB1* co-loss as aggressive HRD-associated CRPC phenotype) | **Caveat 1 (*RB1* is HR-supporting, not strictly an HR gene).** *RB1* loss compromises HR repair through mechanistic studies (CtIP end resection, BRG1 recruitment, 53BP1 binding, NHEJ regulation[11]) but is **not equivalent to *BRCA1/2* loss**. The strongest framing is that *RB1* loss *contributes to* HRD-associated genomic phenotypes rather than equating to bona fide HRD. **Caveat 2 (the *BRCA2/RB1* co-loss signature reflects 13q proximity, applies to Subject 7 only).** *RB1* (13q14) and *BRCA2* (13q13) are physically adjacent on chromosome 13q, so a single |

| Gene | Pathway / role | Subject(s) | reference(s) | Notes |
|---|---|---|---|---|
| | | | | deletion event can hit both loci. In Subject 7, both BRCA2 deletion and RB1 deletion are truncal and may reflect a single 13q deletion event, making the apparent HRD contribution largely attributable to BRCA2 deletion. In Subject 49, BRCA2 deletion occurs on the declining clone and independently on the resistant clone, while RB1 is altered by a resistant-clone inversion; these are independent events on the SV tree, not a 13q-proximity artifact. |
| ***PTEN*** | Tumor suppressor; PI3K | 6 (truncal del), 12 (truncal del), 18 (truncal inv), 20 (truncal del) | Hosseini 2025[33], Quigley 2018[34] | Canonical mCRPC tumor-suppressor loss |
| ***NKX3-1*** | Prostate-lineage tumor suppressor | 6 (resistant del), 18 (truncal inv) | Hosseini 2025[33], Quigley 2018[34] | |

| Gene | Pathway / role | Subject(s) | reference(s) | Notes |
| --- | --- | --- | --- | --- |
| ***CHEK2*** | DDR / cell-cycle checkpoint | 10 (truncal intragenic del) | Hosseini 2025[33] | DDR-pathway driver |
| ***BRCA2*** | Homologous recombination | 7 (truncal del), 20 (truncal fs), 31 (sensitive del), 47 (sensitive inv), 49 (sensitive del + resistant del/inv) | Hosseini 2025[33] | PARP-inhibitor target; HRD biomarker |
| ***ATM*** | Homologous recombination | 31 (sensitive del + resistant inv) | Hosseini 2025[33]. | DDR-pathway driver |
| ***RAD51B*** | Homologous recombination (BCDX2 complex) | 7 (sensitive inv), 10 (sensitive hem del), 20 (truncal inv), 49 (sensitive inv) | Hosseini 2025[33], Setton 2021[15] (biallelic RAD51B alteration is confirmed to produce a measurable HRD phenotype by functional assay; RAD51B is a member of the BCDX2 complex required for stabilizing the RAD51 presynaptic filament during strand invasion) | HR-pathway driver; RAD51B disruption contributes to HRD in sensitive clones (Subjects 7, 10, 49) and as a truncal event in all clones (Subject 20) |
| ***RAD51C*** | Homologous recombination | 13 (truncal inv) | Hosseini 2025[33] | HR-pathway driver |
| ***PALB2*** | Homologous recombination | 10 (sensitive del), 18 | Hosseini 2025[33] | BRCA2-binding partner |

| Gene | Pathway / role | Subject(s) | reference(s) | Notes |
|---|---|---|---|---|
| | | (truncal inv), 47 (truncal del + sensitive del + resistant del) | | |
| ***CHD1*** | Chromatin remodeling; HR repair | 31 (truncal hem del), 47 (sensitive inv), 49 (sensitive del) | Hosseini 2025[33], Quigley 2018[34], Kari 2016[19] (CHD1 loss causes DNA repair defects by impairing nucleosome eviction at DSBs; sensitizes prostate cancer to DNA damaging therapy in vitro, in vivo, and in patient-derived organoids; synthetic lethality with PARP inhibition); Shenoy 2017[18] (CHD1 loss promotes error-prone NHEJ over HR in DSB repair) | Recurrent prostate-cancer driver; HRD-contributing event in the Subject 47 and 49 sensitive clones |
| ***SETD2*** | Histone H3K36 methyltransferase; replication-stress tolerance | 1, 12 (resistant inv), 18 (truncal del), 20 (truncal inv), 49 | Hosseini 2025[33], Kanu 2015[36] (SETD2 loss promotes replication fork slowing, altered origin | Chromatin-modifier driver; *SETD2 inv/del on resistant clones* (Subjects 1, 12) is interpreted as an epigenetic |

| Gene | Pathway / role | Subject(s) | reference(s) | Notes |
|---|---|---|---|---|
| | | (sensitive inv) | firing, and impaired DNA damage repair; associated with branched clonal evolution through replication stress, paradoxically conferring stress tolerance in adapted cells rather than uniform sensitivity) | adaptation that enables replication-stress tolerance under BAT cycling |
| ***ARID1A*** | SWI/SNF chromatin remodeling | 6 (sensitive del), 7 (truncal inv + sensitive del) | Hosseini 2025[33] | SWI/SNF complex driver |
| ***ARID2*** | SWI/SNF chromatin remodeling | 6 (sensitive inv) | Hosseini 2025[33] | |
| ***KMT2C*** | Histone H3K4 methyltransferase | 6 (truncal SNV fs), 20 (truncal SNV nonsense), 47 (truncal SNV fs) | Hosseini 2025[33] | Recurrent SNV across cohort |
| ***KMT2D*** | Histone H3K4 methyltransferase | 31 (truncal SNV nonsense) | Hosseini 2025[33] | |
| ***MLH1*** | Mismatch repair | 20 (truncal inv + sensitive del) | Hosseini 2025[33] | MMR-pathway driver |
| ***MSH2*** | Mismatch repair | 7 (truncal del + sensitive inv) | Hosseini 2025[33] | |

| Gene | Pathway / role | Subject(s) | reference(s) | Notes |
|---|---|---|---|---|
| ***MSH6*** | Mismatch repair | 7 (truncal del + sensitive inv) | Hosseini 2025[33] | |
| ***MYC*** | Oncogene; transcription factor | 31 (resistant dup); 10 (context only: regulator of CUL1[34]) | Hosseini 2025[33], Quigley 2018[34], Viswanathan 2018[35] | Major mCRPC driver |
| ***MYCN*** | Oncogene; MYC family | 7 (truncal dup) | Hosseini 2025[33] | Lineage-plasticity driver |
| ***MDM2*** | p53 negative regulator | 10 (truncal dup), 13 (sensitive dup), 20 (resistant dup) | Sondka 2018[7] (canonical Tier 1 oncogene), Oliner 2016[16] (MDM2 amplification as a bona fide cancer driver), Chopra 2018[17] (MDM2 amplified in ~25% of CRPC, MDMX co-amplified in ~32%, mostly *TP53*-WT) | Amplification is the canonical activating mechanism for *MDM2*; CRPC-specific evidence supports MDM2 duplication as oncogenic on a wild-type *TP53* background |
| ***AURKA*** | Mitotic kinase; AR-V7 regulator | 6 (resistant dup) | Hosseini 2025[33], Jones 2017[9] (Aurora Kinase A reconfigures AR pre-mRNA splicing to upregulate AR-V7 transcripts selectively, without affecting full-length AR mRNA; AURKA | In Subject 6, AURKA duplication on the resistant clone provides a ligand-independent AR-V7-driven survival program that decouples the clone from BAT's testosterone- |

| Gene | Pathway / role | Subject(s) | reference(s) | Notes |
|---|---|---|---|---|
| | | | knockdown markedly reduces AR-V7-driven proliferation and survival), Antonarakis 2014[10] (AR-V7 detection in circulating tumor cells associates with resistance to enzalutamide and abiraterone in CRPC) | cycling mechanism |
| ***CDK4*** | Cyclin-dependent kinase | 13 (sensitive dup) | Hosseini 2025[33] | Cell-cycle driver |
| ***CDK12*** | Transcriptional CDK; DDR; genomic instability | 13 (sensitive del) | Hosseini 2025[33], Frank 2025[37] (acute CDK12 loss causes premature intronic polyadenylation of HRR genes including ATM, inducing an HRD phenotype; chronic CDK12 loss in established tumors produces the hallmark focal tandem duplication signature with | mCRPC subtype marker; branch-specific CDK12 deletion on the sensitive clone (Clone 2) in Subject 13 creates a branch-specific genomic instability background (tandem duplications, potential HRR gene downregulation) |

| Gene | Pathway / role | Subject(s) | reference(s) | Notes |
|---|---|---|---|---|
| | | | partial HRR adaptation, cells may not respond to PARP inhibition despite apparent HRD) | |
| ***CDKN2A*** | Tumor suppressor; cell cycle checkpoint (p16, p14ARF) | 12 (sensitive del), 20 (truncal SNV missense) | Hosseini 2025[33] | In Subject 12, co-loss of *CDKN2A* and *CDKN2B* on the sensitive clone abolishes G1/S (p16 → CDK4/6) and p53-stabilization (p14ARF → MDM2) checkpoints simultaneously, preventing G1 arrest in response to BAT-induced DSBs and driving cells into mitotic catastrophe |
| ***CDKN2B*** | Tumor suppressor; cell cycle checkpoint (p15) | 12 (sensitive del) | Hosseini 2025[33] | p15 provides a backup CDK4/6 inhibitory function for p16; combined *CDKN2A/B* loss removes both primary and backup G1 checkpoints |
| ***CUL1*** | SCF E3 ubiquitin ligase catalytic subunit | 10 (resistant intragenic dup) | O'Hagan 2000[38] (*CUL1* = direct c-MYC transcriptional target), Yang 2002[39] | Subject 10 Clone 3 candidate-resistance event |

| Gene | Pathway / role | Subject(s) | reference(s) | Notes |
|---|---|---|---|---|
| | | | (Skp2/p27 axis in prostate-cancer progression) | |
| ***BRAF*** | RAS/RAF/MEK pathway | 1 (resistant dup) | Sondka 2018[7] (canonical Tier 1 oncogene; copy-number gains explicitly recognized alongside point mutations and fusions); Chehrazi-Raffle 2023[8] (activating *BRAF* alterations, including class II mutations, fusions, and copy-number changes, as a unique spectrum of oncogenic events in prostate cancer) | *BRAF duplication has functional consequences* for MAPK pathway activity, including in prostate cancer (and as a known mechanism of acquired resistance to MAPK inhibitors in melanoma) |
| ***KRAS*** | RAS/RAF/MEK pathway | 13 (sensitive dup) | Hosseini 2025[33] | |
| ***AKT1*** | PI3K/AKT pathway | 10 (truncal SNV E17K hotspot) | Hosseini 2025[33] | E17K is a canonical AKT1 hotspot; truncal in Subject 10 |
| ***PIK3R1*** | PI3K regulatory subunit | 20 (truncal SNV fs) | Hosseini 2025[33] | PI3K-pathway driver |
| ***CTNNB1*** | Wnt pathway / AR coactivator | 12 (sensitive S45P SNV) | Hosseini 2025[33], Chesire & Isaacs 2002[40] *Oncogene* (β- | S45P stabilizes β-catenin, which coactivates AR; under |

| Gene | Pathway / role | Subject(s) | reference(s) | Notes |
|---|---|---|---|---|
| | | | catenin physically interacts with the AR ligand-binding domain and potentiates AR-driven transcription); Truica et al. 2000[41] *Cancer Res* (β-catenin acts as a direct AR coactivator) | supraphysiologic testosterone (BAT), enhanced AR transcriptional output amplifies Topoisomerase IIβ-mediated DSB burden, contributing to BAT sensitivity rather than resistance as would be expected under AR-targeted therapy |
| ***APC*** | Wnt pathway tumor suppressor | 49 (sensitive del) | Hosseini 2025[33] | |
| ***ONECUT2*** | Lineage-plasticity transcription factor; AR-axis suppressor | 49 (sensitive + resistant dup, parallel branch-specific events) | Hosseini 2025[33], Rotinen 2018[20] (ONECUT2 is a targetable master regulator of lethal prostate cancer: directly activates the glucocorticoid receptor *GR/NR3C1* and NE splicing factor *SRRM4*, suppresses the androgen axis, and pharmacologic inhibition restores enzalutamide | In Subject 49, ONECUT2 duplication arises independently on both branches (parallel evolution); on the resistant clone it enables complete AR-axis bypass via neuroendocrine reprogramming, overriding HRD-mediated BAT sensitivity; the sensitive clone carries the same duplication but is eliminated by its co-occurring |

| Gene | Pathway / role | Subject(s) | reference(s) | Notes |
|---|---|---|---|---|
| | | | sensitivity), Guo 2019[21] (ONECUT2 is a validated driver of neuroendocrine prostate cancer; selection for ONECUT2-expressing cells occurs under AR-targeted therapy) | triple HRR deficiency (*BRCA2* + *CHD1* + *RAD51B*) and hyper-AR-dependent state |
| ***RSPO2*** | Wnt-pathway agonist | 13 (sensitive dup), 31 (resistant dup) | Hosseini 2025[33] | |
| ***ZFHX3*** | Tumor suppressor | 13 (sensitive del) | Hosseini 2025[33] | Recurrent prostate-cancer driver |
| ***MGA*** | MAX-network transcription factor | 13 (truncal del + sensitive inv) | Hosseini 2025[33] | |
| ***ERF*** | ETS-family transcription factor | 7 (truncal del), 47 (truncal inv) | Hosseini 2025[33] | mCRPC driver |
| ***NCOR2*** | AR co-repressor | 20 (truncal SNV missense) | Hosseini 2025[33] | |
| ***XPO1*** | Nuclear export | 20 (sensitive SNV missense) | Hosseini 2025[33] | |
| ***TMPRSS2 (deletion without ERG fusion)*** | AR-axis | 20 (sensitive del) | Hosseini 2025[33] | Distinct from the *TMPRSS2-ERG* fusion: deletion of TMPRSS2 itself rather than fusion to ERG |

**Table S7.** Per-cell statistics for Supplementary Fig. S3. Mean signed SVCF error (truth minus estimate) for each method, the paired effect size Δ (the difference in absolute mean signed error, |SVCFit| minus |SVclone|) with its bootstrap 95% confidence interval, and the exact and Benjamini-Hochberg-adjusted paired Wilcoxon p-values, for all structural variants, by tumor purity, SV-CNV overlap configuration, and SV type.

| Purity | Configuration | SV type | SVCFit mean | SVclone mean | Δ \|error\| | Δ 95% CI | p (exact) | p (BH) |
|---|---|---|---|---|---|---|---|---|
| 80% | No overlapping CNV | deletion | 2.0e-04 | -0.1858 | -0.1816 | [-0.1841, -0.179] | 1.9e-09 | 3.1e-09 |
| 80% | No overlapping CNV | tandem duplication | -0.0183 | -0.2635 | -0.2445 | [-0.2483, -0.2407] | 1.9e-09 | 3.1e-09 |
| 80% | No overlapping CNV | inversion | 0.002 | -0.048 | -0.0429 | [-0.0456, -0.0401] | 1.9e-09 | 3.1e-09 |
| 80% | No overlapping CNV | translocation | 0.209 | -0.3472 | -0.1383 | [-0.1535, -0.1235] | 1.9e-09 | 3.1e-09 |
| 80% | SV before cis duplication | deletion | -0.0616 | -0.2471 | -0.1854 | [-0.1881, -0.183] | 1.9e-09 | 3.1e-09 |
| 80% | SV before cis duplication | inversion | -0.012 | -0.1732 | -0.1613 | [-0.1635, -0.159] | 1.9e-09 | 3.1e-09 |
| 80% | SV before cis duplication | translocation | 0.1783 | -0.2773 | -0.099 | [-0.1158, -0.0824] | 1.3e-08 | 1.8e-08 |
| 80% | SV before trans duplication | deletion | -0.0389 | -0.2903 | -0.2514 | [-0.2573, -0.2454] | 1.9e-09 | 3.1e-09 |
| 80% | SV before trans duplication | inversion | 0.0914 | -0.2519 | -0.159 | [-0.1893, -0.1306] | 1.9e-09 | 3.1e-09 |
| 80% | SV before trans duplication | translocation | 0.213 | -0.2903 | -0.0773 | [-0.1017, -0.0531] | 2.8e-06 | 3.6e-06 |
| 80% | SV after duplication | deletion | 0.1051 | 0.0265 | 0.0785 | [0.0738, 0.0839] | 1.9e-09 | 3.1e-09 |
| 80% | SV after duplication | inversion | 0.1063 | 0.0925 | 0.0138 | [0.0075, 0.0199] | 2.8e-04 | 3.3e-04 |
| 80% | SV after duplication | translocation | 0.2149 | -0.225 | -0.0101 | [-0.0237, 0.0039] | 0.191 | 0.198 |
| 80% | SV after trans deletion | deletion | -0.061 | -0.0289 | 0.032 | [0.0301, 0.034] | 1.9e-09 | 3.1e-09 |
| 80% | SV after trans deletion | inversion | 0.0319 | 0.0014 | 0.0195 | [0.0111, 0.0279] | 2.3e-04 | 2.8e-04 |
| 80% | SV after trans deletion | translocation | 0.2914 | -0.0259 | 0.2654 | [0.2563, 0.2753] | 1.9e-09 | 3.1e-09 |
| 60% | No overlapping CNV | deletion | -0.0048 | -0.1488 | -0.1435 | [-0.1449, -0.142] | 1.9e-09 | 3.1e-09 |
| 60% | No overlapping CNV | tandem duplication | -0.0272 | -0.2395 | -0.2123 | [-0.2169, -0.2078] | 1.9e-09 | 3.1e-09 |
| 60% | No overlapping CNV | inversion | 0.0053 | -0.0558 | -0.0485 | [-0.0505, -0.0465] | 1.9e-09 | 3.1e-09 |
| 60% | No overlapping CNV | translocation | 0.1756 | -0.3009 | -0.1253 | [-0.1397, -0.1116] | 1.9e-09 | 3.1e-09 |
| 60% | SV before cis duplication | deletion | -0.0836 | -0.2015 | -0.118 | [-0.1201, -0.1157] | 1.9e-09 | 3.1e-09 |
| 60% | SV before cis duplication | inversion | -0.0038 | -0.1624 | -0.1554 | [-0.1575, -0.1534] | 1.9e-09 | 3.1e-09 |
| 60% | SV before cis duplication | translocation | 0.1334 | -0.2177 | -0.0843 | [-0.0936, -0.0742] | 1.9e-09 | 3.1e-09 |
| 60% | SV before trans duplication | deletion | -0.0695 | -0.176 | -0.1065 | [-0.111, -0.1021] | 1.9e-09 | 3.1e-09 |
| 60% | SV before trans duplication | inversion | 0.0534 | -0.1591 | -0.1023 | [-0.138, -0.0683] | 8.0e-06 | 1.0e-05 |
| 60% | SV before trans duplication | translocation | 0.1466 | -0.2147 | -0.0681 | [-0.0871, -0.049] | 4.7e-07 | 6.2e-07 |
| 60% | SV after duplication | deletion | 0.073 | 0.0478 | 0.0252 | [0.0216, 0.0286] | 1.9e-09 | 3.1e-09 |
| 60% | SV after duplication | inversion | 0.0799 | 0.0878 | -0.0078 | [-0.0118, -0.0038] | 0.00219 | 0.00243 |
| 60% | SV after duplication | translocation | 0.1623 | -0.1624 | -2.0e-04 | [-0.0118, 0.0116] | 0.871 | 0.871 |
| 60% | SV after trans deletion | deletion | -0.0555 | -0.0504 | 0.0051 | [0.0028, 0.0074] | 2.6e-04 | 3.1e-04 |
| 60% | SV after trans deletion | inversion | 0.0361 | -0.0183 | 0.0112 | [-0.0013, 0.0233] | 0.124 | 0.132 |
| 60% | SV after trans deletion | translocation | 0.2214 | -0.038 | 0.1834 | [0.1734, 0.1933] | 1.9e-09 | 3.1e-09 |
| 40% | No overlapping CNV | deletion | -0.0074 | -0.113 | -0.1031 | [-0.1051, -0.1011] | 1.9e-09 | 3.1e-09 |
| 40% | No overlapping CNV | tandem duplication | -0.158 | -0.1924 | -0.0343 | [-0.0407, -0.028] | 9.3e-09 | 1.3e-08 |
| 40% | No overlapping CNV | inversion | 0.0069 | -0.0403 | -0.0321 | [-0.035, -0.0293] | 1.9e-09 | 3.1e-09 |
| 40% | No overlapping CNV | translocation | 0.1159 | -0.2533 | -0.1374 | [-0.149, -0.1249] | 1.9e-09 | 3.1e-09 |
| 40% | SV before cis duplication | deletion | -0.0954 | -0.1661 | -0.0706 | [-0.0728, -0.0684] | 1.9e-09 | 3.1e-09 |
| 40% | SV before cis duplication | inversion | -0.0069 | -0.1317 | -0.1241 | [-0.1273, -0.1205] | 1.9e-09 | 3.1e-09 |
| 40% | SV before cis duplication | translocation | 0.0861 | -0.1547 | -0.0685 | [-0.0788, -0.0575] | 1.9e-09 | 3.1e-09 |
| 40% | SV before trans duplication | deletion | -0.0775 | -0.1308 | -0.0532 | [-0.0564, -0.0498] | 1.9e-09 | 3.1e-09 |
| 40% | SV before trans duplication | inversion | 0.015 | -0.113 | -0.0815 | [-0.1012, -0.0611] | 1.3e-07 | 1.7e-07 |
| 40% | SV before trans duplication | translocation | 0.085 | -0.1528 | -0.0679 | [-0.0802, -0.0549] | 3.7e-09 | 5.6e-09 |
| 40% | SV after duplication | deletion | 0.0458 | 0.0258 | 0.02 | [0.0176, 0.0225] | 1.9e-09 | 3.1e-09 |
| 40% | SV after duplication | inversion | 0.0561 | 0.0542 | 0.0019 | [-9.0e-04, 0.0048] | 0.245 | 0.251 |
| 40% | SV after duplication | translocation | 0.1092 | -0.1184 | -0.0092 | [-0.0171, -0.0017] | 0.0277 | 0.0304 |
| 40% | SV after trans deletion | deletion | -0.043 | -0.0592 | -0.0162 | [-0.0181, -0.0145] | 1.9e-09 | 3.1e-09 |
| 40% | SV after trans deletion | inversion | 0.02 | -0.045 | -0.0214 | [-0.0328, -0.0091] | 9.5e-04 | 0.0011 |
| 40% | SV after trans deletion | translocation | 0.1365 | -0.0487 | 0.0878 | [0.0781, 0.0968] | 1.9e-09 | 3.1e-09 |

| Purity | Configuration | SV type | SVCFit mean | SVclone mean | Δ \|error\| | Δ 95% CI | p (exact) | p (BH) |
|---|---|---|---|---|---|---|---|---|
| 20% | No overlapping CNV | deletion | -0.0151 | -0.0798 | -0.0603 | [-0.0646, -0.056] | 1.9e-09 | 3.1e-09 |
| 20% | No overlapping CNV | tandem duplication | -0.3409 | -0.1352 | 0.2057 | [0.1945, 0.2175] | 1.9e-09 | 3.1e-09 |
| 20% | No overlapping CNV | inversion | -0.0015 | -0.0369 | -0.0332 | [-0.0357, -0.0311] | 1.9e-09 | 3.1e-09 |
| 20% | No overlapping CNV | translocation | 0.0451 | -0.1502 | -0.1051 | [-0.1125, -0.0979] | 1.9e-09 | 3.1e-09 |
| 20% | SV before cis duplication | deletion | -0.074 | -0.0954 | -0.0214 | [-0.0237, -0.0189] | 1.9e-09 | 3.1e-09 |
| 20% | SV before cis duplication | inversion | -0.0363 | -0.066 | -0.0297 | [-0.032, -0.0274] | 1.9e-09 | 3.1e-09 |
| 20% | SV before cis duplication | translocation | 0.0258 | -0.1242 | -0.0982 | [-0.1052, -0.0908] | 1.9e-09 | 3.1e-09 |
| 20% | SV before trans duplication | deletion | -0.1075 | -0.099 | 0.0084 | [0.0038, 0.013] | 0.00158 | 0.00178 |
| 20% | SV before trans duplication | inversion | -0.0159 | -0.052 | -0.0347 | [-0.0402, -0.0291] | 3.7e-09 | 5.6e-09 |
| 20% | SV before trans duplication | translocation | 0.0296 | -0.1079 | -0.0778 | [-0.0859, -0.0702] | 1.9e-09 | 3.1e-09 |
| 20% | SV after duplication | deletion | -0.0011 | -0.0066 | -7.0e-04 | [-0.0026, 0.0011] | 0.299 | 0.303 |
| 20% | SV after duplication | inversion | 0.0162 | -9.0e-04 | 0.0131 | [0.011, 0.0152] | 9.3e-09 | 1.3e-08 |
| 20% | SV after duplication | translocation | 0.037 | -0.0893 | -0.0522 | [-0.0581, -0.0455] | 1.9e-09 | 3.1e-09 |
| 20% | SV after trans deletion | deletion | -0.0435 | -0.067 | -0.0235 | [-0.0247, -0.0223] | 1.9e-09 | 3.1e-09 |
| 20% | SV after trans deletion | inversion | -0.0019 | -0.0531 | -0.0446 | [-0.0497, -0.0392] | 3.7e-09 | 5.6e-09 |
| 20% | SV after trans deletion | translocation | 0.0625 | -0.0684 | -0.0059 | [-0.0137, 0.0025] | 0.177 | 0.187 |
| 10% | No overlapping CNV | deletion | -0.0447 | -0.0617 | -0.017 | [-0.0217, -0.0124] | 3.5e-08 | 4.8e-08 |
| 10% | No overlapping CNV | tandem duplication | -0.6505 | -0.0937 | 0.5568 | [0.5458, 0.5687] | 1.9e-09 | 3.1e-09 |
| 10% | No overlapping CNV | inversion | -0.0213 | -0.036 | -0.0147 | [-0.0175, -0.012] | 3.7e-09 | 5.6e-09 |
| 10% | No overlapping CNV | translocation | -5.0e-04 | -0.0985 | -0.0874 | [-0.0925, -0.0823] | 3.7e-09 | 5.6e-09 |
| 10% | SV before cis duplication | deletion | -0.0637 | -0.0542 | 0.0095 | [0.0074, 0.0116] | 9.3e-09 | 1.3e-08 |
| 10% | SV before cis duplication | inversion | -0.0455 | -0.0338 | 0.0118 | [0.0102, 0.0134] | 1.9e-09 | 3.1e-09 |
| 10% | SV before cis duplication | translocation | -0.0178 | -0.0941 | -0.0759 | [-0.0801, -0.0714] | 1.9e-09 | 3.1e-09 |
| 10% | SV before trans duplication | deletion | -0.1288 | -0.0885 | 0.0402 | [0.0361, 0.0445] | 1.9e-09 | 3.1e-09 |
| 10% | SV before trans duplication | inversion | -0.0557 | -0.0729 | -0.0172 | [-0.0243, -0.0096] | 1.9e-04 | 2.3e-04 |
| 10% | SV before trans duplication | translocation | -0.0102 | -0.0889 | -0.0764 | [-0.0813, -0.0716] | 1.9e-09 | 3.1e-09 |
| 10% | SV after duplication | deletion | -0.0338 | -0.0293 | 0.0045 | [0.0021, 0.0069] | 0.00158 | 0.00178 |
| 10% | SV after duplication | inversion | -0.017 | -0.0186 | -0.0015 | [-0.0032, 2.0e-04] | 0.105 | 0.113 |
| 10% | SV after duplication | translocation | -4.0e-04 | -0.0777 | -0.0674 | [-0.073, -0.0616] | 1.9e-09 | 3.1e-09 |
| 10% | SV after trans deletion | deletion | -0.0686 | -0.0742 | -0.0057 | [-0.0081, -0.0032] | 7.9e-05 | 9.9e-05 |
| 10% | SV after trans deletion | inversion | -0.0254 | -0.0618 | -0.0364 | [-0.0399, -0.0326] | 1.9e-09 | 3.1e-09 |
| 10% | SV after trans deletion | translocation | 0.0096 | -0.0775 | -0.0603 | [-0.0697, -0.0517] | 7.4e-09 | 1.1e-08 |

**Table S8.** Per-cell statistics for Supplementary Fig. S4. Mean signed SVCF error (truth minus estimate) for each method, the paired effect size Δ (the difference in absolute mean signed error, |SVCFit| minus |SVclone|) with its bootstrap 95% confidence interval, and the exact and Benjamini-Hochberg-adjusted paired Wilcoxon p-values, for all structural variants, by tumor purity, configuration, and zygosity (heterozygous, homozygous, hemizygous).

| Purity | Configuration | Zygosity | SVCFit mean | SVclone mean | Δ \|error\| | Δ 95% CI | p (exact) | p (BH) |
|---|---|---|---|---|---|---|---|---|
| 80% | No overlapping CNV | heterozygous | -0.1287 | -0.348 | -0.2193 | [-0.2267, -0.2119] | 1.9e-09 | 2.9e-09 |
| 80% | No overlapping CNV | homozygous | 0.0796 | -0.6504 | -0.5707 | [-0.5777, -0.5636] | 1.9e-09 | 2.9e-09 |
| 80% | No overlapping CNV | hemizygous | 0.0173 | -0.0093 | 0.008 | [0.0065, 0.0096] | 3.7e-09 | 5.6e-09 |
| 80% | SV before cis duplication | heterozygous | -0.2152 | -0.2252 | -0.01 | [-0.0173, -0.0029] | 0.0106 | 0.0116 |
| 80% | SV before cis duplication | homozygous | 0.0471 | -0.535 | -0.4875 | [-0.4969, -0.4776] | 1.9e-09 | 2.9e-09 |
| 80% | SV before cis duplication | hemizygous | 0.005 | -0.1679 | -0.1629 | [-0.1644, -0.1613] | 1.9e-09 | 2.9e-09 |
| 80% | SV before trans duplication | heterozygous | -0.0129 | -0.1101 | -0.092 | [-0.1014, -0.0829] | 1.9e-09 | 2.9e-09 |
| 80% | SV before trans duplication | homozygous | 0.0577 | -0.5382 | -0.4804 | [-0.4889, -0.4718] | 1.9e-09 | 2.9e-09 |
| 80% | SV after duplication | heterozygous | 0.0091 | -0.0568 | -0.0399 | [-0.0471, -0.0323] | 9.3e-09 | 1.3e-08 |
| 80% | SV after duplication | homozygous | 0.2479 | -0.1889 | 0.059 | [0.0514, 0.0666] | 1.9e-09 | 2.9e-09 |
| 80% | SV after duplication | hemizygous | 0.1127 | 0.1024 | 0.0103 | [0.0049, 0.0159] | 0.00237 | 0.00273 |
| 80% | SV after trans deletion | heterozygous | 0.016 | -0.0257 | -0.0096 | [-0.0143, -0.0045] | 5.6e-04 | 6.7e-04 |
| 60% | No overlapping CNV | heterozygous | -0.1264 | -0.2934 | -0.1669 | [-0.1753, -0.1592] | 1.9e-09 | 2.9e-09 |
| 60% | No overlapping CNV | homozygous | 0.0482 | -0.5262 | -0.478 | [-0.4832, -0.473] | 1.9e-09 | 2.9e-09 |
| 60% | No overlapping CNV | hemizygous | 0.0175 | -0.0242 | -0.0067 | [-0.0081, -0.0052] | 9.3e-09 | 1.3e-08 |
| 60% | SV before cis duplication | heterozygous | -0.2503 | -0.1711 | 0.0792 | [0.0748, 0.0837] | 1.9e-09 | 2.9e-09 |
| 60% | SV before cis duplication | homozygous | -0.0177 | -0.3727 | -0.3533 | [-0.3588, -0.3479] | 1.9e-09 | 2.9e-09 |
| 60% | SV before cis duplication | hemizygous | 0.0087 | -0.1616 | -0.1529 | [-0.1547, -0.151] | 1.9e-09 | 2.9e-09 |
| 60% | SV before trans duplication | heterozygous | -0.026 | -0.0495 | -0.0221 | [-0.0284, -0.0159] | 5.7e-07 | 7.8e-07 |
| 60% | SV before trans duplication | homozygous | -0.014 | -0.3791 | -0.3646 | [-0.3705, -0.359] | 1.9e-09 | 2.9e-09 |
| 60% | SV after duplication | heterozygous | 0.0023 | -0.0014 | 0.0033 | [-0.0014, 0.0077] | 0.299 | 0.315 |
| 60% | SV after duplication | homozygous | 0.1649 | -0.0995 | 0.0654 | [0.0601, 0.0713] | 1.9e-09 | 2.9e-09 |
| 60% | SV after duplication | hemizygous | 0.0853 | 0.0925 | -0.0072 | [-0.0101, -0.0041] | 9.9e-05 | 1.2e-04 |
| 60% | SV after trans deletion | heterozygous | 0.0033 | -0.0449 | -0.0372 | [-0.0421, -0.0317] | 3.7e-09 | 5.6e-09 |
| 40% | No overlapping CNV | heterozygous | -0.312 | -0.2248 | 0.0872 | [0.077, 0.0977] | 1.9e-09 | 2.9e-09 |
| 40% | No overlapping CNV | homozygous | -0.0236 | -0.3744 | -0.3508 | [-0.3562, -0.3459] | 1.9e-09 | 2.9e-09 |
| 40% | No overlapping CNV | hemizygous | 0.0196 | -0.0254 | -0.0058 | [-0.0091, -0.0022] | 0.00123 | 0.00145 |
| 40% | SV before cis duplication | heterozygous | -0.2658 | -0.1698 | 0.096 | [0.0913, 0.1008] | 1.9e-09 | 2.9e-09 |
| 40% | SV before cis duplication | homozygous | -0.0411 | -0.2754 | -0.2343 | [-0.2376, -0.231] | 1.9e-09 | 2.9e-09 |
| 40% | SV before cis duplication | hemizygous | 0.0053 | -0.1287 | -0.1227 | [-0.1246, -0.1203] | 1.9e-09 | 2.9e-09 |
| 40% | SV before trans duplication | heterozygous | -0.0346 | -0.0351 | -5.0e-04 | [-0.0051, 0.0046] | 0.7 | 0.7 |
| 40% | SV before trans duplication | homozygous | -0.0502 | -0.2806 | -0.2304 | [-0.2342, -0.2269] | 1.9e-09 | 2.9e-09 |
| 40% | SV after duplication | heterozygous | -0.0093 | -0.0077 | 0.0016 | [-0.0013, 0.0047] | 0.503 | 0.511 |
| 40% | SV after duplication | homozygous | 0.0924 | -0.0834 | 0.009 | [0.0044, 0.014] | 0.00299 | 0.00332 |
| 40% | SV after duplication | hemizygous | 0.0637 | 0.0592 | 0.0045 | [0.0019, 0.007] | 0.00256 | 0.0029 |
| 40% | SV after trans deletion | heterozygous | -0.0037 | -0.0553 | -0.0491 | [-0.0517, -0.0458] | 1.9e-09 | 2.9e-09 |
| 20% | No overlapping CNV | heterozygous | -0.3919 | -0.1417 | 0.2502 | [0.2393, 0.2605] | 1.9e-09 | 2.9e-09 |
| 20% | No overlapping CNV | homozygous | -0.0903 | -0.2035 | -0.1132 | [-0.1185, -0.1079] | 1.9e-09 | 2.9e-09 |
| 20% | No overlapping CNV | hemizygous | 0.0227 | -0.0197 | 0.003 | [-5.0e-04, 0.0066] | 0.129 | 0.139 |
| 20% | SV before cis duplication | heterozygous | -0.1827 | -0.1274 | 0.0554 | [0.0506, 0.0601] | 1.9e-09 | 2.9e-09 |
| 20% | SV before cis duplication | homozygous | -0.0691 | -0.1824 | -0.1133 | [-0.1163, -0.1103] | 1.9e-09 | 2.9e-09 |
| 20% | SV before cis duplication | hemizygous | -0.0126 | -0.0591 | -0.0465 | [-0.0481, -0.045] | 1.9e-09 | 2.9e-09 |
| 20% | SV before trans duplication | heterozygous | -0.0695 | -0.0503 | 0.0192 | [0.0133, 0.0244] | 1.7e-06 | 2.2e-06 |
| 20% | SV before trans duplication | homozygous | -0.0774 | -0.1827 | -0.1053 | [-0.1099, -0.1011] | 1.9e-09 | 2.9e-09 |
| 20% | SV after duplication | heterozygous | -0.0337 | -0.018 | 0.0158 | [0.0111, 0.0199] | 1.0e-06 | 1.3e-06 |
| 20% | SV after duplication | homozygous | 0.0052 | -0.0766 | -0.0678 | [-0.0722, -0.0634] | 1.9e-09 | 2.9e-09 |
| 20% | SV after duplication | hemizygous | 0.0254 | 0.0052 | 0.02 | [0.0183, 0.0217] | 1.9e-09 | 2.9e-09 |
| 20% | SV after trans deletion | heterozygous | -0.0173 | -0.0637 | -0.0464 | [-0.0485, -0.0441] | 1.9e-09 | 2.9e-09 |
| 10% | No overlapping CNV | heterozygous | -0.5004 | -0.0946 | 0.4059 | [0.3926, 0.419] | 1.9e-09 | 2.9e-09 |

| Purity | Configuration | Zygosity | SVCFit mean | SVclone mean | Δ \|error\| | Δ 95% CI | p (exact) | p (BH) |
|---|---|---|---|---|---|---|---|---|
| 10% | No overlapping CNV | homozygous | -0.249 | -0.1083 | 0.1406 | [0.1305, 0.1513] | 1.9e-09 | 2.9e-09 |
| 10% | No overlapping CNV | hemizygous | 0.0103 | -0.0085 | 0.0025 | [-3.0e-04, 0.0055] | 0.328 | 0.34 |
| 10% | SV before cis duplication | heterozygous | -0.1334 | -0.0927 | 0.0407 | [0.037, 0.0443] | 1.9e-09 | 2.9e-09 |
| 10% | SV before cis duplication | homozygous | -0.0688 | -0.1031 | -0.0343 | [-0.0386, -0.03] | 1.9e-09 | 2.9e-09 |
| 10% | SV before cis duplication | hemizygous | -0.0213 | -0.024 | -0.0027 | [-0.0039, -0.0013] | 5.1e-04 | 6.2e-04 |
| 10% | SV before trans duplication | heterozygous | -0.1062 | -0.0795 | 0.0267 | [0.0228, 0.0307] | 1.9e-09 | 2.9e-09 |
| 10% | SV before trans duplication | homozygous | -0.0655 | -0.1039 | -0.0385 | [-0.0428, -0.034] | 1.9e-09 | 2.9e-09 |
| 10% | SV after duplication | heterozygous | -0.0606 | -0.0472 | 0.0133 | [0.0098, 0.0168] | 2.5e-07 | 3.6e-07 |
| 10% | SV after duplication | homozygous | -0.0149 | -0.0623 | -0.0473 | [-0.0514, -0.0425] | 1.9e-09 | 2.9e-09 |
| 10% | SV after duplication | hemizygous | -0.0011 | -0.009 | -0.0041 | [-0.0055, -0.0027] | 9.2e-06 | 1.2e-05 |
| 10% | SV after trans deletion | heterozygous | -0.0427 | -0.0687 | -0.0261 | [-0.0283, -0.0236] | 1.9e-09 | 2.9e-09 |

**Table S9.** Per-cell statistics for Supplementary Fig. S5. Mean signed SVCF error (truth minus estimate) for each method, the paired effect size Δ (the difference in absolute mean signed error, |SVCFit| minus |SVclone|) with its bootstrap 95% confidence interval, and the exact and Benjamini-Hochberg-adjusted paired Wilcoxon p-values, for clonal structural variants only, by tumor purity, configuration, and SV type.

| Purity | Configuration | SV type | SVCFit mean | SVclone mean | Δ \|error\| | Δ 95% CI | p (exact) | p (BH) |
|---|---|---|---|---|---|---|---|---|
| 80% | No overlapping CNV | deletion | 0.0077 | -0.1895 | -0.181 | [-0.1852, -0.1767] | 1.9e-09 | 4.4e-09 |
| 80% | No overlapping CNV | tandem duplication | 0.0297 | -0.2488 | -0.216 | [-0.2229, -0.2092] | 1.9e-09 | 4.4e-09 |
| 80% | No overlapping CNV | inversion | 3.0e-04 | -0.0458 | -0.0381 | [-0.0428, -0.034] | 1.9e-09 | 4.4e-09 |
| 80% | No overlapping CNV | translocation | 0.2527 | -0.4783 | -0.2256 | [-0.2629, -0.1894] | 1.9e-09 | 4.4e-09 |
| 80% | SV before cis duplication | deletion | -0.0479 | -0.1347 | -0.0868 | [-0.0905, -0.0832] | 1.9e-09 | 4.4e-09 |
| 80% | SV before cis duplication | inversion | 0.0025 | -0.0176 | -0.0107 | [-0.0141, -0.0072] | 1.2e-05 | 1.7e-05 |
| 80% | SV before cis duplication | translocation | 0.3148 | -0.2606 | 0.0543 | [0.0369, 0.0734] | 8.3e-07 | 1.3e-06 |
| 80% | SV before trans duplication | deletion | -0.0723 | -0.2713 | -0.199 | [-0.2108, -0.1867] | 1.9e-09 | 4.4e-09 |
| 80% | SV before trans duplication | inversion | 0.0961 | -0.4084 | -0.3121 | [-0.3667, -0.2527] | 5.6e-09 | 1.2e-08 |
| 80% | SV before trans duplication | translocation | 0.3645 | -0.2648 | 0.0997 | [0.0798, 0.1193] | 1.3e-08 | 2.5e-08 |
| 80% | SV after duplication | deletion | 0.0873 | 0.1682 | -0.0809 | [-0.0878, -0.074] | 1.9e-09 | 4.4e-09 |
| 80% | SV after duplication | inversion | 0.0505 | 0.2791 | -0.2286 | [-0.2379, -0.219] | 1.9e-09 | 4.4e-09 |
| 80% | SV after duplication | translocation | 0.2995 | -0.2203 | 0.0792 | [0.0625, 0.0943] | 3.5e-08 | 6.2e-08 |
| 80% | SV after trans deletion | deletion | -0.0652 | -0.0752 | -0.01 | [-0.0132, -0.0068] | 6.9e-06 | 1.0e-05 |
| 80% | SV after trans deletion | inversion | 0.0334 | -0.0264 | 0.0018 | [-0.0129, 0.018] | 0.626 | 0.634 |
| 80% | SV after trans deletion | translocation | 0.4929 | -0.0787 | 0.4143 | [0.395, 0.4345] | 1.9e-09 | 4.4e-09 |
| 60% | No overlapping CNV | deletion | 0.0017 | -0.1389 | -0.1324 | [-0.1349, -0.1297] | 1.9e-09 | 4.4e-09 |
| 60% | No overlapping CNV | tandem duplication | 0.0355 | -0.2497 | -0.2142 | [-0.2202, -0.2086] | 1.9e-09 | 4.4e-09 |
| 60% | No overlapping CNV | inversion | 0.0056 | -0.0425 | -0.0331 | [-0.0377, -0.0282] | 1.9e-09 | 4.4e-09 |
| 60% | No overlapping CNV | translocation | 0.2241 | -0.3788 | -0.1548 | [-0.1866, -0.1243] | 1.3e-08 | 2.5e-08 |
| 60% | SV before cis duplication | deletion | -0.0974 | -0.0925 | 0.0049 | [0.0014, 0.0086] | 0.0277 | 0.0336 |
| 60% | SV before cis duplication | inversion | 0.0119 | -0.0092 | 0.0038 | [0, 0.0076] | 0.114 | 0.127 |
| 60% | SV before cis duplication | translocation | 0.2321 | -0.168 | 0.0641 | [0.0423, 0.0862] | 1.0e-06 | 1.6e-06 |
| 60% | SV before trans duplication | deletion | -0.0897 | -0.1084 | -0.0187 | [-0.028, -0.0096] | 5.1e-04 | 6.7e-04 |
| 60% | SV before trans duplication | inversion | 0.0928 | -0.2126 | -0.1254 | [-0.211, -0.0431] | 0.0137 | 0.0168 |
| 60% | SV before trans duplication | translocation | 0.2617 | -0.1663 | 0.0954 | [0.0796, 0.1112] | 1.9e-09 | 4.4e-09 |
| 60% | SV after duplication | deletion | 0.0563 | 0.1629 | -0.1066 | [-0.1115, -0.1019] | 1.9e-09 | 4.4e-09 |
| 60% | SV after duplication | inversion | 0.0343 | 0.2024 | -0.1681 | [-0.1727, -0.1637] | 1.9e-09 | 4.4e-09 |
| 60% | SV after duplication | translocation | 0.2304 | -0.1434 | 0.087 | [0.0694, 0.1036] | 1.9e-08 | 3.4e-08 |
| 60% | SV after trans deletion | deletion | -0.0628 | -0.0802 | -0.0175 | [-0.0213, -0.0138] | 3.7e-09 | 8.3e-09 |
| 60% | SV after trans deletion | inversion | 0.085 | -0.0177 | 0.0412 | [0.0153, 0.0665] | 0.00538 | 0.00673 |
| 60% | SV after trans deletion | translocation | 0.3482 | -0.0667 | 0.2799 | [0.2608, 0.2981] | 1.9e-09 | 4.4e-09 |
| 40% | No overlapping CNV | deletion | -0.0066 | -0.0944 | -0.0871 | [-0.0889, -0.0854] | 1.9e-09 | 4.4e-09 |
| 40% | No overlapping CNV | tandem duplication | -0.1787 | -0.1816 | -0.0029 | [-0.0122, 0.007] | 0.404 | 0.42 |
| 40% | No overlapping CNV | inversion | 0.0142 | -0.0216 | -0.0058 | [-0.0109, -6.0e-04] | 0.0767 | 0.0877 |
| 40% | No overlapping CNV | translocation | 0.1633 | -0.2757 | -0.1124 | [-0.1432, -0.079] | 9.2e-06 | 1.3e-05 |
| 40% | SV before cis duplication | deletion | -0.1341 | -0.1095 | 0.0247 | [0.0219, 0.0273] | 1.9e-09 | 4.4e-09 |
| 40% | SV before cis duplication | inversion | 0.0072 | -0.0093 | 0.0029 | [-9.0e-04, 0.0068] | 0.221 | 0.232 |
| 40% | SV before cis duplication | translocation | 0.1433 | -0.1044 | 0.0389 | [0.0196, 0.0567] | 3.4e-04 | 4.7e-04 |
| 40% | SV before trans duplication | deletion | -0.0729 | -0.0907 | -0.0178 | [-0.0249, -0.0111] | 6.3e-05 | 8.7e-05 |
| 40% | SV before trans duplication | inversion | 0.0781 | -0.1652 | -0.0839 | [-0.1355, -0.0352] | 0.00322 | 0.00409 |
| 40% | SV before trans duplication | translocation | 0.1634 | -0.0964 | 0.067 | [0.0489, 0.0842] | 1.6e-07 | 2.8e-07 |
| 40% | SV after duplication | deletion | 0.0316 | 0.0873 | -0.0557 | [-0.0588, -0.0528] | 1.9e-09 | 4.4e-09 |
| 40% | SV after duplication | inversion | 0.0266 | 0.1304 | -0.1038 | [-0.1081, -0.0997] | 1.9e-09 | 4.4e-09 |
| 40% | SV after duplication | translocation | 0.1571 | -0.0908 | 0.0663 | [0.057, 0.0757] | 1.9e-09 | 4.4e-09 |
| 40% | SV after trans deletion | deletion | -0.0446 | -0.0703 | -0.0256 | [-0.0284, -0.0227] | 1.9e-09 | 4.4e-09 |
| 40% | SV after trans deletion | inversion | 0.0558 | -0.0335 | 0.0214 | [8.0e-04, 0.0416] | 0.0473 | 0.0556 |

| Purity | Configuration | SV type | SVCFit mean | SVclone mean | Δ \|error\| | Δ 95% CI | p (exact) | p (BH) |
|---|---|---|---|---|---|---|---|---|
| 40% | SV after trans deletion | translocation | 0.209 | -0.0448 | 0.1639 | [0.1504, 0.1775] | 1.9e-09 | 4.4e-09 |
| 20% | No overlapping CNV | deletion | -0.0065 | -0.0613 | -0.0422 | [-0.0465, -0.038] | 1.9e-09 | 4.4e-09 |
| 20% | No overlapping CNV | tandem duplication | -0.3025 | -0.1125 | 0.19 | [0.1809, 0.1995] | 1.9e-09 | 4.4e-09 |
| 20% | No overlapping CNV | inversion | 0.008 | -0.0128 | -0.003 | [-0.0057, -4.0e-04] | 0.0384 | 0.0459 |
| 20% | No overlapping CNV | translocation | 0.0718 | -0.1563 | -0.0846 | [-0.0971, -0.0718] | 3.7e-09 | 8.3e-09 |
| 20% | SV before cis duplication | deletion | -0.1034 | -0.0632 | 0.0402 | [0.0373, 0.0432] | 1.9e-09 | 4.4e-09 |
| 20% | SV before cis duplication | inversion | 7.0e-04 | -0.0062 | 0.0033 | [7.0e-04, 0.0061] | 0.0919 | 0.104 |
| 20% | SV before cis duplication | translocation | 0.0552 | -0.0773 | -0.0219 | [-0.0347, -0.0099] | 0.00219 | 0.00282 |
| 20% | SV before trans duplication | deletion | -0.0948 | -0.0507 | 0.0441 | [0.0374, 0.0505] | 1.9e-09 | 4.4e-09 |
| 20% | SV before trans duplication | inversion | 0.0262 | -0.0413 | -0.0188 | [-0.0388, 0.0018] | 0.129 | 0.142 |
| 20% | SV before trans duplication | translocation | 0.0657 | -0.0643 | 0.0015 | [-0.0081, 0.0117] | 0.839 | 0.839 |
| 20% | SV after duplication | deletion | -0.0103 | 0.0231 | -0.0116 | [-0.0144, -0.0087] | 2.0e-07 | 3.4e-07 |
| 20% | SV after duplication | inversion | 0.0116 | 0.0278 | -0.0157 | [-0.0182, -0.0133] | 1.9e-09 | 4.4e-09 |
| 20% | SV after duplication | translocation | 0.058 | -0.063 | -0.005 | [-0.0158, 0.0056] | 0.529 | 0.543 |
| 20% | SV after trans deletion | deletion | -0.0335 | -0.0588 | -0.0253 | [-0.0275, -0.0232] | 1.9e-09 | 4.4e-09 |
| 20% | SV after trans deletion | inversion | 0.0206 | -0.0376 | -0.0137 | [-0.0281, -5.0e-04] | 0.164 | 0.177 |
| 20% | SV after trans deletion | translocation | 0.0885 | -0.0526 | 0.0333 | [0.0195, 0.0458] | 5.0e-05 | 7.0e-05 |
| 10% | No overlapping CNV | deletion | -0.0373 | -0.0491 | -0.0115 | [-0.0172, -0.0057] | 8.0e-04 | 0.00105 |
| 10% | No overlapping CNV | tandem duplication | -0.6145 | -0.0794 | 0.535 | [0.519, 0.5509] | 1.9e-09 | 4.4e-09 |
| 10% | No overlapping CNV | inversion | -0.0124 | -0.0223 | -0.0098 | [-0.013, -0.0066] | 2.3e-06 | 3.5e-06 |
| 10% | No overlapping CNV | translocation | 0.0111 | -0.0924 | -0.0749 | [-0.0799, -0.0699] | 3.0e-08 | 5.3e-08 |
| 10% | SV before cis duplication | deletion | -0.0694 | -0.0334 | 0.036 | [0.0334, 0.0384] | 1.9e-09 | 4.4e-09 |
| 10% | SV before cis duplication | inversion | -0.0111 | -0.0038 | 0.0074 | [0.0052, 0.0094] | 5.7e-07 | 9.3e-07 |
| 10% | SV before cis duplication | translocation | -0.0096 | -0.061 | -0.0398 | [-0.0466, -0.0323] | 1.9e-08 | 3.4e-08 |
| 10% | SV before trans duplication | deletion | -0.1219 | -0.057 | 0.0649 | [0.0589, 0.0709] | 1.9e-09 | 4.4e-09 |
| 10% | SV before trans duplication | inversion | -0.0428 | -0.0254 | 0.0071 | [-0.0045, 0.0192] | 0.204 | 0.218 |
| 10% | SV before trans duplication | translocation | 0.0024 | -0.0532 | -0.0345 | [-0.041, -0.0268] | 1.7e-06 | 2.6e-06 |
| 10% | SV after duplication | deletion | -0.037 | -0.0078 | 0.0292 | [0.026, 0.0327] | 1.9e-09 | 4.4e-09 |
| 10% | SV after duplication | inversion | -0.0123 | -6.0e-04 | 0.0108 | [0.0085, 0.013] | 9.3e-09 | 1.9e-08 |
| 10% | SV after duplication | translocation | 0.0087 | -0.0528 | -0.0351 | [-0.0395, -0.0309] | 1.9e-09 | 4.4e-09 |
| 10% | SV after trans deletion | deletion | -0.0669 | -0.0707 | -0.0037 | [-0.0076, -1.0e-04] | 0.0732 | 0.0849 |
| 10% | SV after trans deletion | inversion | -0.0173 | -0.0518 | -0.0343 | [-0.0392, -0.0291] | 5.6e-09 | 1.2e-08 |
| 10% | SV after trans deletion | translocation | 0.0144 | -0.0744 | -0.052 | [-0.0624, -0.0425] | 1.5e-08 | 2.8e-08 |

**Table S10.** Per-cell statistics for Supplementary Fig. S6. Mean signed SVCF error (truth minus estimate) for each method, the paired effect size Δ (the difference in absolute mean signed error, |SVCFit| minus |SVclone|) with its bootstrap 95% confidence interval, and the exact and Benjamini-Hochberg-adjusted paired Wilcoxon p-values, for subclonal structural variants only, by tumor purity, configuration, and SV type.

| Purity | Configuration | SV type | SVCFit mean | SVclone mean | Δ \|error\| | Δ 95% CI | p (exact) | p (BH) |
|---|---|---|---|---|---|---|---|---|
| 80% | No overlapping CNV | deletion | -0.0056 | -0.183 | -0.1737 | [-0.1769, -0.1698] | 1.9e-09 | 2.7e-09 |
| 80% | No overlapping CNV | tandem duplication | -0.0431 | -0.2711 | -0.2281 | [-0.233, -0.223] | 1.9e-09 | 2.7e-09 |
| 80% | No overlapping CNV | inversion | 0.0029 | -0.0492 | -0.0419 | [-0.045, -0.0386] | 1.9e-09 | 2.7e-09 |
| 80% | No overlapping CNV | translocation | 0.1895 | -0.2895 | -0.1 | [-0.1138, -0.0867] | 1.9e-09 | 2.7e-09 |
| 80% | SV before cis duplication | deletion | -0.0694 | -0.3109 | -0.2415 | [-0.2448, -0.2386] | 1.9e-09 | 2.7e-09 |
| 80% | SV before cis duplication | inversion | -0.0196 | -0.2555 | -0.2359 | [-0.2392, -0.2324] | 1.9e-09 | 2.7e-09 |
| 80% | SV before cis duplication | translocation | 0.1372 | -0.2809 | -0.1436 | [-0.1624, -0.1219] | 5.6e-09 | 7.4e-09 |
| 80% | SV before trans duplication | deletion | -0.0129 | -0.305 | -0.2895 | [-0.2942, -0.2846] | 1.9e-09 | 2.7e-09 |
| 80% | SV before trans duplication | inversion | 0.087 | -0.1551 | -0.0525 | [-0.0856, -0.0207] | 0.00664 | 0.00672 |
| 80% | SV before trans duplication | translocation | 0.1603 | -0.2966 | -0.1363 | [-0.1644, -0.1086] | 3.5e-08 | 4.4e-08 |
| 80% | SV after duplication | deletion | 0.117 | -0.0673 | 0.0497 | [0.0406, 0.0582] | 5.6e-09 | 7.4e-09 |
| 80% | SV after duplication | inversion | 0.1387 | -0.0146 | 0.1233 | [0.1179, 0.1279] | 1.9e-09 | 2.7e-09 |
| 80% | SV after duplication | translocation | 0.1708 | -0.228 | -0.0572 | [-0.0749, -0.0388] | 4.4e-06 | 4.8e-06 |
| 80% | SV after trans deletion | deletion | -0.0574 | 0.0095 | 0.0463 | [0.0388, 0.0533] | 3.7e-09 | 5.2e-09 |
| 80% | SV after trans deletion | inversion | 0.0307 | 0.0135 | 0.013 | [0.0053, 0.0208] | 0.00299 | 0.00306 |
| 80% | SV after trans deletion | translocation | 0.2238 | -0.0118 | 0.2067 | [0.1991, 0.2144] | 1.9e-09 | 2.7e-09 |
| 60% | No overlapping CNV | deletion | -0.0098 | -0.1565 | -0.1467 | [-0.1491, -0.1443] | 1.9e-09 | 2.7e-09 |
| 60% | No overlapping CNV | tandem duplication | -0.0605 | -0.2341 | -0.1736 | [-0.1796, -0.1677] | 1.9e-09 | 2.7e-09 |
| 60% | No overlapping CNV | inversion | 0.005 | -0.0633 | -0.0552 | [-0.0576, -0.0527] | 1.9e-09 | 2.7e-09 |
| 60% | No overlapping CNV | translocation | 0.1513 | -0.264 | -0.1127 | [-0.1265, -0.0988] | 1.9e-09 | 2.7e-09 |
| 60% | SV before cis duplication | deletion | -0.0757 | -0.2635 | -0.1878 | [-0.19, -0.1853] | 1.9e-09 | 2.7e-09 |
| 60% | SV before cis duplication | inversion | -0.0121 | -0.2431 | -0.2307 | [-0.2338, -0.2277] | 1.9e-09 | 2.7e-09 |
| 60% | SV before cis duplication | translocation | 0.1022 | -0.2332 | -0.131 | [-0.1442, -0.1178] | 1.9e-09 | 2.7e-09 |
| 60% | SV before trans duplication | deletion | -0.0533 | -0.2296 | -0.1763 | [-0.1816, -0.1712] | 1.9e-09 | 2.7e-09 |
| 60% | SV before trans duplication | inversion | 0.0296 | -0.1404 | -0.1021 | [-0.1303, -0.0759] | 4.7e-07 | 5.5e-07 |
| 60% | SV before trans duplication | translocation | 0.1078 | -0.2276 | -0.1198 | [-0.142, -0.0962] | 1.9e-09 | 2.7e-09 |
| 60% | SV after duplication | deletion | 0.0847 | -0.0323 | 0.0524 | [0.0465, 0.0591] | 1.9e-09 | 2.7e-09 |
| 60% | SV after duplication | inversion | 0.1077 | 0.0183 | 0.0893 | [0.0845, 0.0943] | 1.9e-09 | 2.7e-09 |
| 60% | SV after duplication | translocation | 0.1246 | -0.1741 | -0.0495 | [-0.0635, -0.0344] | 2.0e-06 | 2.2e-06 |
| 60% | SV after trans deletion | deletion | -0.0495 | -0.0252 | 0.0239 | [0.0208, 0.027] | 1.9e-09 | 2.7e-09 |
| 60% | SV after trans deletion | inversion | 0.0121 | -0.0172 | -0.0065 | [-0.0159, 0.0027] | 0.237 | 0.237 |
| 60% | SV after trans deletion | translocation | 0.1712 | -0.0291 | 0.1413 | [0.1314, 0.1505] | 1.9e-09 | 2.7e-09 |
| 40% | No overlapping CNV | deletion | -0.0081 | -0.13 | -0.116 | [-0.1197, -0.1123] | 1.9e-09 | 2.7e-09 |
| 40% | No overlapping CNV | tandem duplication | -0.1455 | -0.1994 | -0.0539 | [-0.0622, -0.0458] | 1.9e-09 | 2.7e-09 |
| 40% | No overlapping CNV | inversion | 0.0023 | -0.0519 | -0.0455 | [-0.0483, -0.0426] | 1.9e-09 | 2.7e-09 |
| 40% | No overlapping CNV | translocation | 0.0928 | -0.241 | -0.1482 | [-0.1609, -0.1359] | 1.9e-09 | 2.7e-09 |
| 40% | SV before cis duplication | deletion | -0.0728 | -0.1991 | -0.1263 | [-0.129, -0.1233] | 1.9e-09 | 2.7e-09 |
| 40% | SV before cis duplication | inversion | -0.0144 | -0.1961 | -0.1817 | [-0.1855, -0.1776] | 1.9e-09 | 2.7e-09 |
| 40% | SV before cis duplication | translocation | 0.0669 | -0.1714 | -0.1045 | [-0.1166, -0.0921] | 1.9e-09 | 2.7e-09 |
| 40% | SV before trans duplication | deletion | -0.0814 | -0.165 | -0.0836 | [-0.0888, -0.0783] | 1.9e-09 | 2.7e-09 |
| 40% | SV before trans duplication | inversion | -0.0172 | -0.0906 | -0.0568 | [-0.0733, -0.0394] | 2.3e-06 | 2.6e-06 |
| 40% | SV before trans duplication | translocation | 0.0555 | -0.1724 | -0.1169 | [-0.1327, -0.1006] | 1.9e-09 | 2.7e-09 |
| 40% | SV after duplication | deletion | 0.0573 | -0.0233 | 0.034 | [0.0293, 0.0386] | 1.9e-09 | 2.7e-09 |
| 40% | SV after duplication | inversion | 0.0754 | 0.0049 | 0.0694 | [0.0664, 0.0724] | 1.9e-09 | 2.7e-09 |
| 40% | SV after duplication | translocation | 0.0803 | -0.1353 | -0.055 | [-0.0632, -0.0463] | 1.9e-09 | 2.7e-09 |
| 40% | SV after trans deletion | deletion | -0.0412 | -0.0481 | -0.0069 | [-0.009, -0.0049] | 2.0e-07 | 2.4e-07 |
| 40% | SV after trans deletion | inversion | 0.0011 | -0.0513 | -0.0348 | [-0.0447, -0.0247] | 5.7e-07 | 6.5e-07 |
| 40% | SV after trans deletion | translocation | 0.1044 | -0.0497 | 0.0547 | [0.0431, 0.0656] | 2.6e-08 | 3.3e-08 |

| Purity | Configuration | SV type | SVCFit mean | SVclone mean | Δ \|error\| | Δ 95% CI | p (exact) | p (BH) |
|---|---|---|---|---|---|---|---|---|
| 20% | No overlapping CNV | deletion | -0.0254 | -0.1017 | -0.0732 | [-0.0776, -0.0685] | 1.9e-09 | 2.7e-09 |
| 20% | No overlapping CNV | tandem duplication | -0.3768 | -0.156 | 0.2208 | [0.2033, 0.2403] | 1.9e-09 | 2.7e-09 |
| 20% | No overlapping CNV | inversion | -0.0097 | -0.0563 | -0.0461 | [-0.0488, -0.0434] | 1.9e-09 | 2.7e-09 |
| 20% | No overlapping CNV | translocation | 0.0311 | -0.1472 | -0.1156 | [-0.1253, -0.1058] | 1.9e-09 | 2.7e-09 |
| 20% | SV before cis duplication | deletion | -0.0547 | -0.1167 | -0.0621 | [-0.0645, -0.0596] | 1.9e-09 | 2.7e-09 |
| 20% | SV before cis duplication | inversion | -0.0563 | -0.0984 | -0.0421 | [-0.0445, -0.0397] | 1.9e-09 | 2.7e-09 |
| 20% | SV before cis duplication | translocation | 0.0125 | -0.1446 | -0.1296 | [-0.1373, -0.1223] | 1.9e-09 | 2.7e-09 |
| 20% | SV before trans duplication | deletion | -0.1198 | -0.1451 | -0.0252 | [-0.0315, -0.0189] | 2.0e-07 | 2.4e-07 |
| 20% | SV before trans duplication | inversion | -0.028 | -0.0542 | -0.0259 | [-0.0327, -0.0182] | 2.8e-06 | 3.0e-06 |
| 20% | SV before trans duplication | translocation | 0.008 | -0.131 | -0.115 | [-0.1242, -0.1058] | 1.9e-09 | 2.7e-09 |
| 20% | SV after duplication | deletion | 0.0103 | -0.0428 | -0.0296 | [-0.034, -0.0253] | 1.9e-09 | 2.7e-09 |
| 20% | SV after duplication | inversion | 0.0203 | -0.0253 | -0.005 | [-0.008, -0.002] | 0.00299 | 0.00306 |
| 20% | SV after duplication | translocation | 0.0216 | -0.1071 | -0.0824 | [-0.0884, -0.0758] | 1.9e-09 | 2.7e-09 |
| 20% | SV after trans deletion | deletion | -0.0576 | -0.0786 | -0.021 | [-0.023, -0.0192] | 1.9e-09 | 2.7e-09 |
| 20% | SV after trans deletion | inversion | -0.0116 | -0.0599 | -0.0466 | [-0.0522, -0.041] | 1.9e-09 | 2.7e-09 |
| 20% | SV after trans deletion | translocation | 0.0491 | -0.0776 | -0.0284 | [-0.0401, -0.0176] | 1.2e-04 | 1.3e-04 |
| 10% | No overlapping CNV | deletion | -0.0567 | -0.0806 | -0.0239 | [-0.029, -0.019] | 1.9e-09 | 2.7e-09 |
| 10% | No overlapping CNV | tandem duplication | -0.6948 | -0.1106 | 0.5841 | [0.5637, 0.6029] | 1.9e-09 | 2.7e-09 |
| 10% | No overlapping CNV | inversion | -0.0326 | -0.0526 | -0.02 | [-0.0243, -0.0161] | 1.9e-09 | 2.7e-09 |
| 10% | No overlapping CNV | translocation | -0.008 | -0.1031 | -0.0845 | [-0.0922, -0.076] | 7.4e-09 | 9.8e-09 |
| 10% | SV before cis duplication | deletion | -0.0583 | -0.0742 | -0.0159 | [-0.0191, -0.0126] | 5.6e-09 | 7.4e-09 |
| 10% | SV before cis duplication | inversion | -0.0674 | -0.0525 | 0.0148 | [0.013, 0.0169] | 1.9e-09 | 2.7e-09 |
| 10% | SV before cis duplication | translocation | -0.0218 | -0.1126 | -0.0908 | [-0.0958, -0.086] | 1.9e-09 | 2.7e-09 |
| 10% | SV before trans duplication | deletion | -0.1356 | -0.1182 | 0.0174 | [0.0124, 0.0226] | 1.6e-07 | 2.0e-07 |
| 10% | SV before trans duplication | inversion | -0.0563 | -0.086 | -0.0297 | [-0.036, -0.0237] | 1.3e-08 | 1.7e-08 |
| 10% | SV before trans duplication | translocation | -0.0195 | -0.1115 | -0.091 | [-0.0964, -0.0854] | 1.9e-09 | 2.7e-09 |
| 10% | SV after duplication | deletion | -0.0281 | -0.0639 | -0.0358 | [-0.0391, -0.0321] | 1.9e-09 | 2.7e-09 |
| 10% | SV after duplication | inversion | -0.0231 | -0.041 | -0.0179 | [-0.0203, -0.0156] | 1.9e-09 | 2.7e-09 |
| 10% | SV after duplication | translocation | -0.0094 | -0.1027 | -0.0864 | [-0.0926, -0.0802] | 1.9e-09 | 2.7e-09 |
| 10% | SV after trans deletion | deletion | -0.0707 | -0.0801 | -0.0094 | [-0.0133, -0.0056] | 5.6e-05 | 6.0e-05 |
| 10% | SV after trans deletion | inversion | -0.0323 | -0.0699 | -0.0376 | [-0.0415, -0.0335] | 1.9e-09 | 2.7e-09 |
| 10% | SV after trans deletion | translocation | 0.0065 | -0.077 | -0.0642 | [-0.0768, -0.0519] | 2.4e-07 | 2.8e-07 |

**Table S11.** Per-cell statistics for Supplementary Fig. S7. Mean signed SVCF error (truth minus estimate) for each method, the paired effect size Δ (the difference in absolute mean signed error, |SVCFit| minus |SVclone|) with its bootstrap 95% confidence interval, and the exact and Benjamini-Hochberg-adjusted paired Wilcoxon p-values, for subclonal structural variants, by tumor purity, configuration, and subclonal mixture setting.

| Purity | Configuration | Mixture (minor:major) | SVCFit mean | SVclone mean | Δ \|error\| | Δ 95% CI | p (exact) | p (BH) |
|---|---|---|---|---|---|---|---|---|
| 80% | No overlapping CNV | 10-90 | -0.0171 | -0.1776 | -0.1593 | [-0.1641, -0.1547] | 1.9e-09 | 2.8e-09 |
| 80% | No overlapping CNV | 30-70 | 0.0034 | -0.1859 | -0.1784 | [-0.1817, -0.1748] | 1.9e-09 | 2.8e-09 |
| 80% | No overlapping CNV | 50-50 | -0.0172 | -0.2082 | -0.1909 | [-0.1943, -0.1876] | 1.9e-09 | 2.8e-09 |
| 80% | SV before cis duplication | 10-90 | -0.0238 | -0.2253 | -0.2015 | [-0.2052, -0.1977] | 1.9e-09 | 2.8e-09 |
| 80% | SV before cis duplication | 30-70 | -0.0423 | -0.3281 | -0.2858 | [-0.2907, -0.2807] | 1.9e-09 | 2.8e-09 |
| 80% | SV before cis duplication | 50-50 | -0.0446 | -0.3105 | -0.266 | [-0.2702, -0.2615] | 1.9e-09 | 2.8e-09 |
| 80% | SV before trans duplication | 10-90 | 0.0773 | -0.3515 | -0.2742 | [-0.289, -0.2585] | 1.9e-09 | 2.8e-09 |
| 80% | SV before trans duplication | 30-70 | 0.0308 | -0.2808 | -0.2488 | [-0.2677, -0.2289] | 1.9e-09 | 2.8e-09 |
| 80% | SV before trans duplication | 50-50 | 0.0017 | -0.2612 | -0.2439 | [-0.2551, -0.2321] | 1.9e-09 | 2.8e-09 |
| 80% | SV after duplication | 10-90 | 0.1746 | -6.0e-04 | 0.1525 | [0.1412, 0.1636] | 1.9e-09 | 2.8e-09 |
| 80% | SV after duplication | 30-70 | 0.1148 | -0.0817 | 0.0331 | [0.0242, 0.0413] | 1.3e-07 | 1.5e-07 |
| 80% | SV after duplication | 50-50 | 0.1109 | -0.0646 | 0.0462 | [0.0367, 0.0552] | 3.7e-09 | 5.3e-09 |
| 80% | SV after trans deletion | 10-90 | 0.0203 | -0.0414 | -0.0152 | [-0.0273, -0.0031] | 0.0185 | 0.0199 |
| 80% | SV after trans deletion | 30-70 | 0.0136 | 0.0102 | 0.0056 | [0.0012, 0.0101] | 0.0208 | 0.022 |
| 80% | SV after trans deletion | 50-50 | 0.0253 | 0.0259 | 8.0e-04 | [-0.0032, 0.0049] | 1 | 1 |
| 60% | No overlapping CNV | 10-90 | -0.0298 | -0.1776 | -0.1478 | [-0.1534, -0.1426] | 1.9e-09 | 2.8e-09 |
| 60% | No overlapping CNV | 30-70 | -0.0151 | -0.1699 | -0.1532 | [-0.1568, -0.1494] | 1.9e-09 | 2.8e-09 |
| 60% | No overlapping CNV | 50-50 | -0.0174 | -0.1642 | -0.1458 | [-0.1499, -0.1418] | 1.9e-09 | 2.8e-09 |
| 60% | SV before cis duplication | 10-90 | -0.0333 | -0.1806 | -0.1472 | [-0.1502, -0.1444] | 1.9e-09 | 2.8e-09 |
| 60% | SV before cis duplication | 30-70 | -0.0438 | -0.2941 | -0.2503 | [-0.2528, -0.2477] | 1.9e-09 | 2.8e-09 |
| 60% | SV before cis duplication | 50-50 | -0.0432 | -0.2845 | -0.2413 | [-0.2453, -0.237] | 1.9e-09 | 2.8e-09 |
| 60% | SV before trans duplication | 10-90 | 0.0268 | -0.2451 | -0.2169 | [-0.2261, -0.2074] | 1.9e-09 | 2.8e-09 |
| 60% | SV before trans duplication | 30-70 | -0.0173 | -0.2259 | -0.2069 | [-0.2151, -0.1982] | 1.9e-09 | 2.8e-09 |
| 60% | SV before trans duplication | 50-50 | -0.032 | -0.2062 | -0.1729 | [-0.1794, -0.1664] | 1.9e-09 | 2.8e-09 |
| 60% | SV after duplication | 10-90 | 0.1442 | -0.0255 | 0.1188 | [0.1124, 0.1247] | 1.9e-09 | 2.8e-09 |
| 60% | SV after duplication | 30-70 | 0.0832 | -0.022 | 0.0612 | [0.0567, 0.0662] | 1.9e-09 | 2.8e-09 |
| 60% | SV after duplication | 50-50 | 0.0792 | -0.014 | 0.0652 | [0.0568, 0.0738] | 1.9e-09 | 2.8e-09 |
| 60% | SV after trans deletion | 10-90 | -9.0e-04 | -0.0636 | -0.0437 | [-0.0533, -0.0334] | 1.3e-07 | 1.5e-07 |
| 60% | SV after trans deletion | 30-70 | -9.0e-04 | -0.0193 | -0.0032 | [-0.0089, 0.0027] | 0.191 | 0.196 |
| 60% | SV after trans deletion | 50-50 | 0.0163 | -0.0092 | 0.0042 | [-0.0026, 0.0114] | 0.28 | 0.284 |
| 40% | No overlapping CNV | 10-90 | -0.0564 | -0.1383 | -0.082 | [-0.0901, -0.0747] | 1.9e-09 | 2.8e-09 |
| 40% | No overlapping CNV | 30-70 | -0.0615 | -0.147 | -0.0854 | [-0.0904, -0.0806] | 1.9e-09 | 2.8e-09 |
| 40% | No overlapping CNV | 50-50 | -0.0473 | -0.1388 | -0.0907 | [-0.098, -0.0838] | 1.9e-09 | 2.8e-09 |
| 40% | SV before cis duplication | 10-90 | -0.0372 | -0.1854 | -0.1482 | [-0.1544, -0.1406] | 1.9e-09 | 2.8e-09 |
| 40% | SV before cis duplication | 30-70 | -0.0473 | -0.1989 | -0.1517 | [-0.1558, -0.1474] | 1.9e-09 | 2.8e-09 |
| 40% | SV before cis duplication | 50-50 | -0.0404 | -0.203 | -0.1627 | [-0.1661, -0.1588] | 1.9e-09 | 2.8e-09 |
| 40% | SV before trans duplication | 10-90 | -0.022 | -0.1482 | -0.1225 | [-0.1301, -0.1157] | 1.9e-09 | 2.8e-09 |
| 40% | SV before trans duplication | 30-70 | -0.0524 | -0.1617 | -0.1093 | [-0.1176, -0.1011] | 1.9e-09 | 2.8e-09 |
| 40% | SV before trans duplication | 50-50 | -0.0568 | -0.1681 | -0.1114 | [-0.1198, -0.1029] | 1.9e-09 | 2.8e-09 |
| 40% | SV after duplication | 10-90 | 0.1074 | 0.0044 | 0.1013 | [0.0957, 0.1086] | 1.9e-09 | 2.8e-09 |
| 40% | SV after duplication | 30-70 | 0.0551 | -0.0178 | 0.0373 | [0.0302, 0.0445] | 1.9e-09 | 2.8e-09 |
| 40% | SV after duplication | 50-50 | 0.0517 | -0.0301 | 0.0216 | [0.0166, 0.0263] | 8.0e-08 | 9.9e-08 |
| 40% | SV after trans deletion | 10-90 | 0.0026 | -0.0596 | -0.0398 | [-0.0485, -0.0313] | 6.2e-08 | 7.8e-08 |
| 40% | SV after trans deletion | 30-70 | -0.0157 | -0.0586 | -0.0413 | [-0.0454, -0.0371] | 1.9e-09 | 2.8e-09 |
| 40% | SV after trans deletion | 50-50 | 0.0043 | -0.0348 | -0.0247 | [-0.0308, -0.0184] | 6.2e-08 | 7.8e-08 |
| 20% | No overlapping CNV | 10-90 | -0.0956 | -0.087 | 0.0086 | [-4.0e-04, 0.0176] | 0.105 | 0.109 |
| 20% | No overlapping CNV | 30-70 | -0.1485 | -0.1224 | 0.0261 | [0.0177, 0.0341] | 8.3e-07 | 9.6e-07 |
| 20% | No overlapping CNV | 50-50 | -0.173 | -0.1163 | 0.0567 | [0.0492, 0.0653] | 1.9e-09 | 2.8e-09 |
| 20% | SV before cis duplication | 10-90 | -0.0345 | -0.0837 | -0.0492 | [-0.0522, -0.0464] | 1.9e-09 | 2.8e-09 |

| Purity | Configuration | Mixture (minor:major) | SVCFit mean | SVclone mean | Δ \|error\| | Δ 95% CI | p (exact) | p (BH) |
|---|---|---|---|---|---|---|---|---|
| 20% | SV before cis duplication | 30-70 | -0.0598 | -0.1196 | -0.0598 | [-0.0621, -0.0575] | 1.9e-09 | 2.8e-09 |
| 20% | SV before cis duplication | 50-50 | -0.0581 | -0.1226 | -0.0644 | [-0.0683, -0.0608] | 1.9e-09 | 2.8e-09 |
| 20% | SV before trans duplication | 10-90 | -0.0462 | -0.0796 | -0.0334 | [-0.0415, -0.0256] | 3.5e-08 | 4.7e-08 |
| 20% | SV before trans duplication | 30-70 | -0.0889 | -0.1415 | -0.0526 | [-0.0618, -0.0438] | 3.7e-09 | 5.3e-09 |
| 20% | SV before trans duplication | 50-50 | -0.1009 | -0.149 | -0.0481 | [-0.0551, -0.0409] | 1.9e-09 | 2.8e-09 |
| 20% | SV after duplication | 10-90 | 0.0541 | -0.0094 | 0.0446 | [0.0395, 0.0495] | 1.9e-09 | 2.8e-09 |
| 20% | SV after duplication | 30-70 | 0.0056 | -0.0594 | -0.0514 | [-0.0547, -0.0479] | 1.9e-09 | 2.8e-09 |
| 20% | SV after duplication | 50-50 | -0.0072 | -0.0466 | -0.0385 | [-0.0418, -0.0353] | 1.9e-09 | 2.8e-09 |
| 20% | SV after trans deletion | 10-90 | -0.0066 | -0.0594 | -0.045 | [-0.0507, -0.0392] | 1.9e-09 | 2.8e-09 |
| 20% | SV after trans deletion | 30-70 | -0.0291 | -0.0768 | -0.0476 | [-0.053, -0.0427] | 1.9e-09 | 2.8e-09 |
| 20% | SV after trans deletion | 50-50 | -0.0317 | -0.0766 | -0.0449 | [-0.0503, -0.04] | 1.9e-09 | 2.8e-09 |
| 10% | No overlapping CNV | 10-90 | -0.2474 | -0.0627 | 0.1847 | [0.1705, 0.2001] | 1.9e-09 | 2.8e-09 |
| 10% | No overlapping CNV | 30-70 | -0.3591 | -0.0932 | 0.2658 | [0.2473, 0.2837] | 1.9e-09 | 2.8e-09 |
| 10% | No overlapping CNV | 50-50 | -0.3683 | -0.1071 | 0.2612 | [0.2409, 0.2812] | 1.9e-09 | 2.8e-09 |
| 10% | SV before cis duplication | 10-90 | -0.0555 | -0.047 | 0.0084 | [0.0056, 0.011] | 3.2e-06 | 3.7e-06 |
| 10% | SV before cis duplication | 30-70 | -0.0689 | -0.0747 | -0.0058 | [-0.0086, -0.0029] | 0.00104 | 0.00115 |
| 10% | SV before cis duplication | 50-50 | -0.0566 | -0.0691 | -0.0125 | [-0.0155, -0.0097] | 8.0e-08 | 9.9e-08 |
| 10% | SV before trans duplication | 10-90 | -0.0701 | -0.0943 | -0.0241 | [-0.0306, -0.0176] | 2.5e-07 | 3.0e-07 |
| 10% | SV before trans duplication | 30-70 | -0.1001 | -0.1186 | -0.0185 | [-0.0249, -0.0121] | 1.8e-05 | 2.0e-05 |
| 10% | SV before trans duplication | 50-50 | -0.1252 | -0.1133 | 0.0119 | [0.0049, 0.0186] | 0.00374 | 0.00407 |
| 10% | SV after duplication | 10-90 | -0.0088 | -0.0409 | -0.0306 | [-0.0335, -0.0277] | 1.9e-09 | 2.8e-09 |
| 10% | SV after duplication | 30-70 | -0.0251 | -0.0668 | -0.0418 | [-0.0449, -0.0389] | 1.9e-09 | 2.8e-09 |
| 10% | SV after duplication | 50-50 | -0.0448 | -0.063 | -0.0182 | [-0.0221, -0.0143] | 2.6e-08 | 3.5e-08 |
| 10% | SV after trans deletion | 10-90 | -0.0373 | -0.0657 | -0.0281 | [-0.034, -0.023] | 3.7e-09 | 5.3e-09 |
| 10% | SV after trans deletion | 30-70 | -0.044 | -0.081 | -0.037 | [-0.0446, -0.0297] | 9.3e-09 | 1.3e-08 |
| 10% | SV after trans deletion | 50-50 | -0.0534 | -0.0835 | -0.0301 | [-0.0367, -0.0238] | 1.1e-08 | 1.5e-08 |

**Table S12.** Downstream decision-boundary statistics for Supplementary Fig. S12. Clone-misassignment and ancestry-ordering-violation rates (%) for SVCFit and SVclone, each with bootstrap 95% confidence intervals (30 replicates), and the between-method difference (percentage points), overall and broken down by chromosome source, clone, and subclonal mixture.

| Metric | Breakdown | n SVs | SVCFit % [95% CI] | SVclone % [95% CI] | Δ (points) [95% CI] |
|---|---|---|---|---|---|
| Clone assignment | overall: All SVs | 194677 | 27.46 [27.32, 27.6] | 45.35 [45.24, 45.47] | -17.89 [-18.06, -17.72] |
| Clone assignment | source: Autosomes | 87947 | 38.78 [38.54, 39.02] | 47.02 [46.81, 47.22] | -8.24 [-8.51, -7.96] |
| Clone assignment | source: chrX | 106730 | 18.14 [17.95, 18.32] | 43.96 [43.82, 44.1] | -25.82 [-26.05, -25.59] |
| Clone assignment | source_clone: Autosomes clonal | 37156 | 26.95 [26.63, 27.27] | 13.87 [13.62, 14.12] | 13.08 [12.67, 13.49] |
| Clone assignment | source_clone: Autosomes sub1 | 19087 | 41.33 [40.81, 41.83] | 61.25 [60.75, 61.77] | -19.92 [-20.48, -19.4] |
| Clone assignment | source_clone: Autosomes sub2 | 31704 | 51.11 [50.61, 51.63] | 76.61 [76.33, 76.89] | -25.5 [-26, -24.99] |
| Clone assignment | source_clone: chrX clonal | 42203 | 20.82 [20.5, 21.14] | 24.07 [23.72, 24.4] | -3.25 [-3.62, -2.88] |
| Clone assignment | source_clone: chrX sub1 | 26998 | 8.06 [7.78, 8.34] | 50.57 [50.04, 51.11] | -42.5 [-43.09, -41.92] |
| Clone assignment | source_clone: chrX sub2 | 37529 | 22.37 [21.98, 22.77] | 61.57 [61.16, 61.95] | -39.19 [-39.72, -38.63] |
| Tree ordering | overall: All conditions | 900 | 5.78 [4.89, 6.78] | 18.33 [17.33, 19.33] | -12.56 [-13.89, -11.22] |
| Tree ordering | mixture: 0.1 | 300 | 8 [6.33, 9.67] | 29 [26.33, 31.67] | -21 [-24, -18] |
| Tree ordering | mixture: 0.3 | 300 | 6.67 [4.67, 8.67] | 20.33 [17.67, 22.67] | -13.67 [-16.67, -10.67] |
| Tree ordering | mixture: 0.5 | 300 | 2.67 [1.33, 4.33] | 5.67 [4, 7.33] | -3 [-5, -1] |

**Table S13.** Per-cell statistics for Figure 3A. Mean signed SVCF error (truth minus estimate) for each method, the paired effect size (|SVCFit| minus |SVclone| mean signed error) with its bootstrap 95% confidence interval, and the exact and Benjamini-Hochberg-adjusted paired Wilcoxon p-values, by SV type, clonality, and tumor purity, pooling autosomal and hemizygous (chrX) structural variants.

| SV type | Clonality | Purity | SVCFit mean | SVclone mean | Δ \|error\| | Δ 95% CI | p (exact) | p (BH) |
|---|---|---|---|---|---|---|---|---|
| deletion | Clonal | 80% | -0.0048 | -0.0711 | -0.0652 | [-0.0678, -0.0627] | 1.9e-09 | 2.3e-09 |
| deletion | Clonal | 40% | -0.0447 | -0.0446 | 1.0e-04 | [-0.0015, 0.0017] | 0.984 | 0.984 |
| deletion | Clonal | 10% | -0.0657 | -0.036 | 0.0297 | [0.0278, 0.0315] | 1.9e-09 | 2.3e-09 |
| deletion | Subclonal | 80% | 0.0019 | -0.1861 | -0.1807 | [-0.1831, -0.178] | 1.9e-09 | 2.3e-09 |
| deletion | Subclonal | 40% | -0.0258 | -0.123 | -0.0971 | [-0.0989, -0.0954] | 1.9e-09 | 2.3e-09 |
| deletion | Subclonal | 10% | -0.0662 | -0.0802 | -0.014 | [-0.0158, -0.0122] | 1.9e-09 | 2.3e-09 |
| inversion | Clonal | 80% | 0.0221 | 0.0756 | -0.0535 | [-0.0569, -0.0503] | 1.9e-09 | 2.3e-09 |
| inversion | Clonal | 40% | 0.0188 | 0.0348 | -0.016 | [-0.0179, -0.0141] | 1.9e-09 | 2.3e-09 |
| inversion | Clonal | 10% | -0.0126 | -0.01 | 0.0026 | [0.0014, 0.0038] | 1.9e-04 | 2.2e-04 |
| inversion | Subclonal | 80% | 0.0436 | -0.1171 | -0.0735 | [-0.0759, -0.0712] | 1.9e-09 | 2.3e-09 |
| inversion | Subclonal | 40% | 0.0187 | -0.0925 | -0.0738 | [-0.0761, -0.0714] | 1.9e-09 | 2.3e-09 |
| inversion | Subclonal | 10% | -0.0522 | -0.0537 | -0.0015 | [-0.0026, -5.0e-04] | 0.012 | 0.0131 |
| tandem duplication | Clonal | 80% | 0.0297 | -0.2488 | -0.216 | [-0.223, -0.2091] | 1.9e-09 | 2.3e-09 |
| tandem duplication | Clonal | 40% | -0.1787 | -0.1816 | -0.0029 | [-0.0126, 0.0073] | 0.404 | 0.422 |
| tandem duplication | Clonal | 10% | -0.6145 | -0.0794 | 0.535 | [0.5188, 0.5524] | 1.9e-09 | 2.3e-09 |
| tandem duplication | Subclonal | 80% | -0.0431 | -0.2711 | -0.2281 | [-0.2332, -0.2232] | 1.9e-09 | 2.3e-09 |
| tandem duplication | Subclonal | 40% | -0.1455 | -0.1994 | -0.0539 | [-0.0623, -0.0465] | 1.9e-09 | 2.3e-09 |
| tandem duplication | Subclonal | 10% | -0.6948 | -0.1106 | 0.5841 | [0.5645, 0.6028] | 1.9e-09 | 2.3e-09 |
| translocation | Clonal | 80% | 0.3456 | -0.2535 | 0.0921 | [0.0811, 0.1024] | 1.9e-09 | 2.3e-09 |
| translocation | Clonal | 40% | 0.167 | -0.1138 | 0.0533 | [0.047, 0.0597] | 1.9e-09 | 2.3e-09 |
| translocation | Clonal | 10% | 0.0039 | -0.063 | -0.054 | [-0.0568, -0.051] | 1.9e-09 | 2.3e-09 |
| translocation | Subclonal | 80% | 0.1744 | -0.2099 | -0.0355 | [-0.045, -0.0255] | 8.3e-07 | 1.0e-06 |
| translocation | Subclonal | 40% | 0.0779 | -0.1469 | -0.069 | [-0.0745, -0.0637] | 1.9e-09 | 2.3e-09 |
| translocation | Subclonal | 10% | -0.0151 | -0.1072 | -0.0921 | [-0.0953, -0.0892] | 1.9e-09 | 2.3e-09 |

**Table S14.** Statistics for Figure 3B. Mean signed SVCF error at low tumor purity (10-20%) by zygosity, including hemizygous (chrX) loci, for each method; the paired effect size with its bootstrap 95% confidence interval; the exact and Benjamini-Hochberg-adjusted paired Wilcoxon p-values; and the more accurate method.

| Zygosity | SVCFit mean | SVclone mean | Δ \|error\| | Δ 95% CI | p (exact) | p (BH) | More accurate |
|---|---|---|---|---|---|---|---|
| heterozygous | -0.166 | -0.083 | 0.083 | [0.0806, 0.0853] | 1.9e-09 | 1.9e-09 | SVclone |
| homozygous | -0.0813 | -0.147 | -0.0657 | [-0.0678, -0.0635] | 1.9e-09 | 1.9e-09 | SVCFit |
| hemizygous | -3.0e-04 | -0.0264 | -0.0246 | [-0.025, -0.0241] | 1.9e-09 | 1.9e-09 | SVCFit |

**Table S15.** Per-condition statistics for Figure 3C. Mean signed CCF error (truth minus estimate) for each method on the prostate two-tumor mixture series, the paired effect size with its bootstrap 95% confidence interval, and the exact and Benjamini-Hochberg-adjusted paired Wilcoxon p-values, by mixing condition.

| Mixture (bM:gM) | SVCFit mean | SVclone mean | Δ \|error\| | Δ 95% CI | p (exact) | p (BH) |
|---|---|---|---|---|---|---|
| 10-90 | 0.1075 | 0.1094 | -0.0019 | [-0.0062, 0.0026] | 0.382 | 0.467 |
| 20-80 | 0.0639 | 0.0851 | -0.0212 | [-0.0269, -0.0154] | 2.5e-07 | 4.0e-07 |
| 30-70 | 0.0196 | 0.0641 | -0.0429 | [-0.0489, -0.0368] | 1.9e-09 | 5.1e-09 |
| 40-60 | -0.0115 | 0.0506 | -0.0317 | [-0.0389, -0.0242] | 1.6e-07 | 3.6e-07 |
| 50-50 | -0.0396 | 0.0384 | 0.001 | [-0.0102, 0.0122] | 0.952 | 0.952 |
| 60-40 | -0.0328 | 0.0337 | -8.0e-04 | [-0.0111, 0.0093] | 0.824 | 0.906 |
| 70-30 | -0.0224 | 0.0424 | -0.0178 | [-0.0282, -0.0066] | 0.00219 | 0.00301 |
| 80-20 | 0.0025 | 0.0685 | -0.0533 | [-0.0608, -0.0463] | 1.9e-09 | 5.1e-09 |
| 90-10 | 0.046 | 0.1099 | -0.0639 | [-0.0695, -0.0587] | 1.9e-09 | 5.1e-09 |
| 4 clust | -0.0908 | -0.0019 | 0.0587 | [0.0438, 0.0726] | 2.5e-07 | 4.0e-07 |
| 5 clust | 0.0054 | 0.0724 | -0.0608 | [-0.0665, -0.0557] | 1.9e-09 | 5.1e-09 |
